# Analysis of historical selection in winter wheat

**DOI:** 10.1101/2022.01.07.475391

**Authors:** Chin Jian Yang, Olufunmilayo Ladejobi, Richard Mott, Wayne Powell, Ian Mackay

## Abstract

Winter wheat is a major crop with a rich selection history in the modern era of crop breeding. Genetic gains across economically important traits like yield have been well characterized and are the major force driving its production. Winter wheat is also an excellent model for analyzing historical genetic selection. As a proof of concept, we analyze two major collections of winter wheat varieties that were bred in western Europe from 1916 to 2010, namely the Triticeae Genome (TG) and WAGTAIL panels, which include 333 and 403 varieties respectively. We develop and apply a selection mapping approach, Regression of Alleles on Years (RALLY), in these panels, as well as in simulated populations. RALLY maps loci under sustained historical selection by using a simple logistic model to regress allele counts on years of variety release. To control for drift-induced allele frequency change, we develop a hybrid approach of genomic control and delta control. Within the TG panel, we identify 22 significant RALLY quantitative selection loci (QSLs) and estimate the local heritabilities for 12 traits across these QSLs. By correlating predicted marker effects with RALLY regression estimates, we show that alleles whose frequencies have increased over time are heavily biased towards conferring positive yield effect, but negative effects in flowering time, lodging, plant height and grain protein content. Altogether, our results (1) demonstrate the use of RALLY to identify selected genomic regions while controlling for drift, and (2) reveal key patterns in the historical selection in winter wheat and guide its future breeding.

**Key Message:** Modelling of the distribution of allele frequency over year of variety release identifies major loci involved in historical breeding of winter wheat.

## Introduction

Modern agriculture benefits from long standing breeding effort in creating new and improved crop varieties over time. Genetic gain is often used as a measure of the success in breeding for trait improvement. For example, in wheat, the genetic gains in yield and other agriculturally valuable traits have been well quantified (Mackay et al. 2011, Tadesse et al. 2019 and Shorinola et al. 2021). The introduction of genomic selection (GS) (Meuwissen et al. 2001) in breeding program further shortens breeding cycles, improves selection accuracy and intensity, and accelerates genetic gain (Voss-Fels et al. 2019). Lastly, genetic gain is further increased by the rise of knowledge exchange between plant and animal breeding through GS (Hickey et al. 2017).

In recent years, there has been a growing interest in mapping quantitative selection loci (QSLs) that are associated with genetic gain independently of any phenotype. The mapping approach typically involves correlating continuous variables, such as year of variety release and geographical parameters, to genomic markers in a historical variety dataset. Conceptually, it is similar to selection mapping which tests for selection signatures among genomic markers using population genetic models (Johnsson 2018). This approach has been variously named as Birth Date Selection Mapping (Decker et al. 2012), Generation Proxy Selection Mapping (Rowan et al. 2021) and EnvGWAS (Li et al. 2020, Sharma et al. 2021). Here, we will refer to it as EnvGWAS because the underlying mixed linear model is no different from a conventional genome-wide association study (GWAS).

Related to EnvGWAS, EigenGWAS uses eigenvectors (principal components) as the dependent variable in a mixed linear model (Li et al. 2020, Sharma et al. 2021). The EigenGWAS approach may yield similar results to EnvGWAS if the dependent variables in EnvGWAS are correlated strongly with any eigenvector. Otherwise, EigenGWAS may identify additional QSLs where it incorporates variables that have not been quantified directly. A key confounding factor for determining whether a locus has been under sustained historical selection or drift is that varieties are linked by a complex historical pedigree and unequal relatedness. By correcting for population structure using a mixed linear model (Yu et al. 2006), year effects and principal components that are associated with drift can be controlled in EnvGWAS and EigenGWAS respectively.

Here, we introduce a new application of an old method by modelling allele frequency change over years in a historical variety dataset. This method, termed Regression of Alleles on Years (RALLY), fits a logistic regression to model the allele count as a dependent variable and the year of variety release as an independent variable. A logistic model is commonly used in case-control studies where the dependent variables are binary traits of whether an individual is diseased and the independent variables are test factors (Prentice and Pyke 1979). A logistic model is appropriate because changes in allele frequencies are small when the starting frequencies are near the extrema, and large when they are intermediate. In addition, the model is bounded asymptotically by 0 and 1. The dependent and independent variables are switched between RALLY and EnvGWAS. Instead of estimating the mean of years of release for each allele in EnvGWAS, RALLY estimates the mean of allele counts for each year, which is equivalent to the allele frequency for a given year. Recently, Looseley et al. (2020) applied a similar approach to RALLY on significant GWAS markers in a historical barley variety dataset. RALLY is a genome-wide approach that employs parametric control (PC) as a correction to drift-induced allele frequency change. PC is a novel hybrid approach of genomic control (GC) (Devlin and Roeder 1999) and delta control (DC) (Gorroochurn et al. 2006), which are two common control approaches against population structure in human GWAS studies without the need for mixed linear model.

Our analyses in the simulated and historical variety datasets demonstrate the usefulness of RALLY in mapping QSLs. We begin the evaluation of RALLY in simulated populations where the truth is known, both with and without selection, to quantify RALLY detection power and limit. We use the simulations to calibrate PC, which is then applied to the two historical winter wheat datasets, namely the panels of Triticeae Genome (TG) (Bentley et al. 2014) and Wheat Association Genetics for Trait Advancement and Improvement of Lineages (WAGTAIL) (Fradgley et al. 2019). The WAGTAIL panel is used only as a replicate RALLY analysis. Within the TG panel, we identify 22 RALLY QSLs and compare them to the GWAS QTLs from Ladejobi et al. (2019). Some notable QSLs include one in 2B which coincides with *Ppd-B1* (Mohler et al. 2004), *Yr7/Yr5/YrSP* (Marchal et al. 2018) and alien introgression from *Triticum timopheevii* (Tsilo et al. 2008, Martynov et al. 2018), as well as another in 6A that coincides with *TaGW2* (Su et al. 2011), *Rht24* (Würschum et al. 2017) and *Rht25* (Mo et al. 2018). To further support the RALLY QSLs, we show that all 22 QSLs have non-zero local heritabilities for at least one trait. Next, we find clear directional selection in traits like flowering time, lodging, yield, plant height and grain protein content by comparing the signs of predicted marker allele effects with their directions of allele frequency change as given by RALLY. By extending the results to pairs of traits, we identify the selection priorities. For example, more ears with lighter grains have been preferred over fewer ears with heavier grains. Finally, we employ the multivariate breeder’s equation (Lande and Arnold 1983) to estimate selection parameters, although our results suggest a limited use in modern crops, in contrast to its original application in evolutionary studies. Overall, we have shown that many major genomic regions have been extensively used in winter wheat breeding and we suggest that future selection should emphasize on improving other unexplored genomic regions.

## Materials and Methods

### Population simulation with and without selection

We initiated our population simulation in a fictitious species with 10 chromosomes (Figure 1). The genetic lengths of the chromosomes were set from 100 to 280 centiMorgans (cM) with an increment of 20 cM in subsequent chromosomes. The populations spanned over 50 generations (years) with and without selection. All the simulations were performed using the “AlphaSimR” package (Gaynor et al. 2021) in R (R Core Team 2021). We first created 32 inbred founders using the *runMacs* function and we placed one marker (segregating site) at every 0.1 cM. Two causal quantitative trait loci (QTLs) were chosen randomly from the markers at each frequency ranging from 1/32 to 16/32, which resulted in 32 QTLs. The QTL effects were drawn from an approximately negative binomial distribution such that the rarer QTL alleles have larger effects than the more common QTL alleles (Figure 1). We standardized the QTL effects such that the total variance of additive genetic or QTL effects is 1. Phenotypic values for each line were set as a sum of QTL effects and residual effects drawn from a normal distribution of mean 0 and variance 1, which is equivalent to a heritability of 0.5.

**Figure 1.**
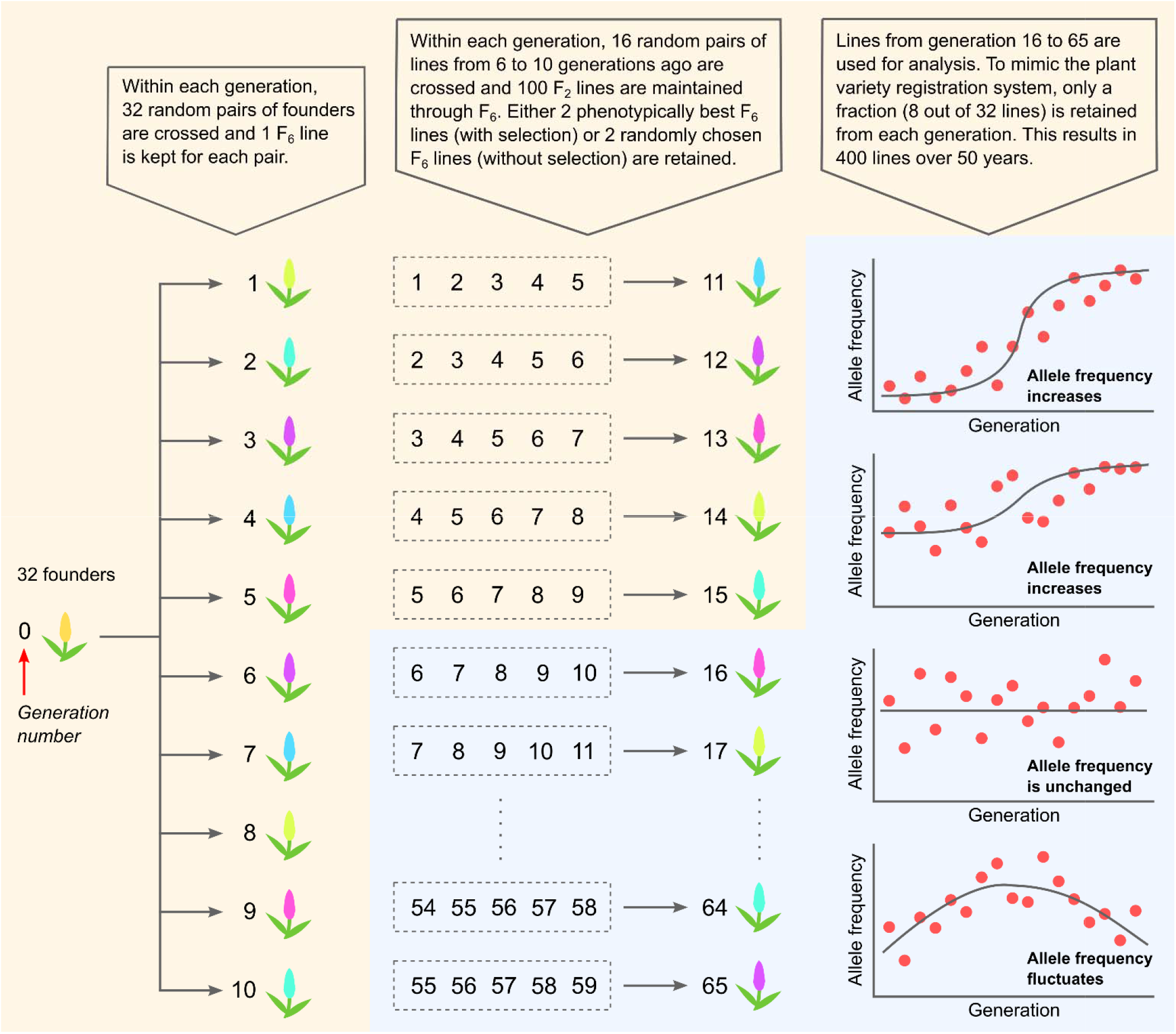
Population simulation and changes in allele frequency over time. The simulated populations with and without selection are described in detail here. The first 15 generations were used as burn-ins and discarded. 8 varieties from each generation starting at 16 and ending at 65 were randomly chosen to create a population of 400 varieties that span over 50 generations. Examples of how allele frequency changes over time are shown, with the first two examples follows a logistic distribution and thus are more likely to be significant under RALLY than the other two examples.

We created selected (S) and unselected (U) populations from the 32 founders using a simplified model that mimics new variety breeding of major crops in Europe. All varieties were derived as F_6_ recombinant inbred lines (RILs) from bi-parental crosses. This is equivalent to 4 generations of single seed descent (SSD) from an F_2_ population. The first 10 generations were created by crossing the initial 32 founders at random. In the subsequent generations, we randomly sampled 32 parents from 6 to 10 generations ago and created 16 bi-parental populations with each having 100 F_6_ RILs. By keeping 2 RILs per bi-parental population, we maintained 32 lines at each generation. The 2 RILs were chosen either from the two highest phenotypic values (selected) or randomly (unselected). This step was repeated until the population underwent 55 generations of phenotypic selection. The first 15 generations were discarded as burn-in because none of the parents of the individuals from these 15 generations have been selected, and hence there is no selection-induced allele frequency change. In our simplistic modelling of the plant variety rights (PVRs) system where only a fraction of new lines passing the PVR test, we randomly sampled and retained 8 lines per generation for a total of 400 lines that spanned over 50 generations. The simulated populations with and without selection were used in subsequent analyses.

### RALLY and GWAS in simulated populations

We compared the performances of Regression of Alleles on Years (RALLY) and Genome Wide Association Study (GWAS) when applied to the selected and unselected populations. The model for RALLY was fitted in a logistic regression using the *glm* function in R (R Core Team 2021). The model for GWAS was fitted in a mixed linear model using the *GWASpoly* function in the “GWASpoly” R package (Rosyara et al. 2016).

Briefly, the logistic regression model for RALLY can be shown as below:

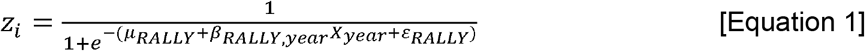

Or, alternatively, the logistic regression model can be rewritten in a linear form as:

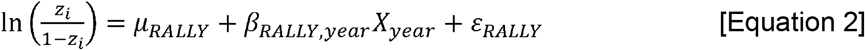

Where the model terms are described as below:

*z_i_* is a binary variable indicating the absence (0) or presence (1) of an allele at marker *i* in *n* lines.
*μ_RALLY_* is the mean allele frequency in the first year, or *y*-intercept.
*β_RALLY,year_* is the fixed year effect, or regression coefficient of the year variable.
*X_year_* is the year variable.
*ε_RALLY_* is the residual effect with a distribution of 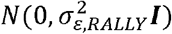 and ***I*** is the identity matrix.

The GWAS model is written as below:

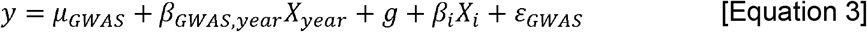

Where the model terms are described as below:

*y* is the trait values in *n* lines.
*μ_GWAS_* is the mean of trait value.
*β_GWAS,year_* is the fixed year effect.
*X_year_* is the year variable.
*g* is the random genetic background effect with a distribution of 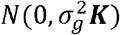 and ***K*** is the additive genetic relationship matrix.
*β_i_* is the fixed allele effect at marker *i*.
*X_i_* is the number of alleles at marker *i*.
*ε_GWAS_* is the residual effect with a distribution of 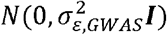 and ***I*** is the identity matrix.

In the given models, the terms of interest are *β_RALLY,year_* and *β_i_* in RALLY and GWAS, respectively. The term significances are determined by their corresponding standard normal *Z*-statistics at a Bonferroni corrected threshold of P = 0.05. Due to how the populations are simulated, some markers may not segregate in all the populations. These markers, along with the QTLs and other markers that are highly linked (r^2^ > 0.99) to QTLs, were removed from the RALLY and GWAS analyses. The simulations were repeated for 100 iterations and the models were fitted for each simulated population separately.

### Model correction by parametric control (PC)

In the previously described naïve RALLY model (Equation 1 and 2), the RALLY test statistics may be inflated by population structure arising from consanguinity and population stratification. These factors can prevent a proper separation of markers under selection or drift if they are not addressed. To control for the inflation, we used a combined approach of genomic control (GC) (Devlin and Roeder 1999) and delta control (DC) (Gorroochurn et al. 2006) which we call parametric control (PC). In the absence of confounding factors, we expect the null test statistics (Z-scores) to be distributed as *N*(0,1). However, in the presence of population structure, the distribution of null test statistics becomes *N*(*δ,σ*). As the terms imply, DC controls the inflation in mean *δ* and GC controls the inflation in standard deviation *σ*. If we can estimate *δ* and *σ*, we can adjust the test statistics as the following:

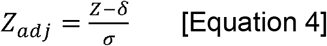

We used a maximum likelihood (ML) approach to estimate *δ* and *σ*. For any value of Z, both positive and negative signs are equally likely because the regression coefficient of one allele has the same magnitude but opposite sign of the other allele. Therefore, we can construct a composite likelihood function from two standard normal probability density functions that account for positive and negative Z values. The likelihood function is shown in Equation 5 below. To simplify the calculation, we used the log likelihood function as described in Equation 6 below. We computed *δ* and *σ* for an *n*-vector of Z values by either maximizing the log likelihood function, or equivalently, minimizing the negative log likelihood function using the “nlm” package in R (R Core Team 2021).

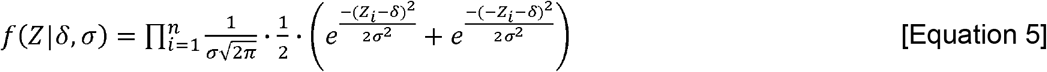

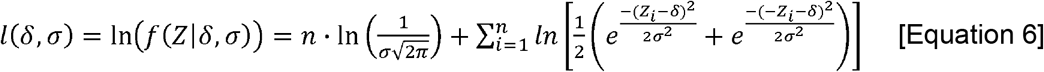

An important factor in PC is the selection of null marker sets for calculating the inflation factors for adjusting the test statistics. The most conservative approach is to use all markers as the null, but this approach is unrealistic as it results in over-correction when the selection is strong and prevalent across the whole genome. Therefore, PC is best estimated from markers that have not undergone selection, although it is paradoxical given that such markers are unknown at this stage. As a compromise, we may assume that the allele frequency differences between first and last years are larger for markers under selection than drift. This assumption is reasonable for a modern breeding population that has undergone intensive selection. We first predicted the allele frequency change for each marker using the RALLY model and then identified the null marker set from markers that fall below various thresholds of allele frequency change. We tested the thresholds ranging from 0.05 to 0.50 at an increment of 0.01, in which the thresholds of 0.05 and 0.50 correspond to 40% and 99% of the total markers respectively. Unfortunately, because the variance of allele frequency change, 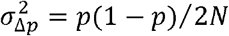, is largest when the initial allele frequencies are intermediate (Falconer and Mackay 1996), a loss in RALLY’s power to distinguish between weak selection signal and drift at those markers is unavoidable.

### Detection limits of RALLY

We estimated the detection limits of RALLY using a simple example that is based on the simulated populations as described previously. We considered a QTL marker and five other proximal markers that are 1, 2, 3, 4 and 5 cM away. The initial QTL frequencies were set to 1/32 to 16/32 with an increment of 1/32, and all possible marker-QTL haplotype frequencies were considered. We modelled selection on the QTL by increasing QTL initial frequency to the final frequency of 31/32 over 50 generations according to either a logistic or linear distribution. Consequently, the proximal markers experienced hitch-hiking effect due to the selection on QTL. Assuming an infinite population size, recombination is the sole factor that is responsible for the hitch-hiking effect, which allowed us to model the change in allele frequencies of the proximal markers. Non-recombinants are inherited at a probability of 1 – *r* and recombinants are inherited at a probability of *r*. From this, we derived the expected allele frequencies for the proximal markers at each generation. Next, we randomly sampled 8 individuals per generation using a binomial distribution with the expected frequencies as the sampling probabilities. This step was repeated for 100 times for each tested marker-QTL haplotype frequencies. A more detailed description of this is provided in Figure S1.

### RALLY in two wheat panels

We first applied the RALLY approach in the Triticeae Genome (TG) panel (Bentley et al. 2014, Ladejobi et al. 2019) as a proof of concept. The TG panel has 344 winter wheat varieties from the UK, France and Germany that were released between 1948 and 2007 (Figure S2), which is ideal for analyzing selection over time in modern wheat breeding. We retained 333 varieties that were in common between the TG panel data derived from DArT markers (Bentley et al. 2014) and genotype-by-sequencing (GBS) markers (Ladejobi et al. 2019). The DArT marker data was only used in a later analysis for estimating multivariate selection parameters. From the initial 41,861 GBS markers, we removed 3,009 markers that are in high linkage disequilibrium (LD) (*r*^2^ > 0.2) with markers from other chromosomes which left us with 38,852 markers. These markers were positioned according to the IWGSC RefSeq v1.0 genome assembly. Here, we applied a similar model to Equation 2 with an additional fixed effect to account for the country of origin. We identified the year regression coefficients, applied the same level of PC as identified from the simulation to adjust the test statistics, and determined the significance at a Bonferroni-corrected threshold of 0.05.

Next, we replicated the analysis in the WAGTAIL panel (Fradgley et al. 2019) to test RALLY performance in a different sampling panel of modern wheat varieties. The WAGTAIL panel has 403 winter wheat varieties of mostly UK origin that were released between 1916 and 2010. Of the 403 varieties, 283 originated from the UK, 51 from France, 34 from Germany and 35 from other countries including Australia, Belgium, Canada, Denmark, the Netherlands, Sweden, Switzerland, and United States. There were 99 overlapping varieties between the TG and WAGTAIL panels. Since the WAGTAIL panel was genotyped using the wheat 90k array (Wang et al. 2014) and did not immediately have physical map positions for direct comparison with the TG panel, we identified the physical map positions from the IWGSC RefSeq v1.0 annotation file. We retained 5,592 out of 26,015 markers that had matching chromosomes between the original WAGTAIL genetic map and the physical map. We also removed 319 markers that are in high LD (*r*^2^ > 0.2) with markers from other chromosomes which left us with 5,273 markers. We applied Equation 2 with an additional fixed country of origin effect to the WAGTAIL panel and computed the year regression coefficients with the same PC and multiple testing correction to the test significances.

### Estimating local heritabilities from RALLY QSLs

We clustered the significant markers identified from RALLY into groups based on the extent of LD surrounding the markers. Because genomic markers are not completely independent, some significant markers may be tagging the same QSLs. Starting with the most significant (focal) markers within each chromosome, we assigned markers that have *r*^2^ > 0.2 with the focal marker to the same group. To avoid incorrectly mapped markers, we require the groups to have a minimum of 10 markers in the TG panel and 5 markers in the WAGTAIL panel due to lower marker density. As a trade-off, there may be bias against genomic regions with sparse marker density such as the D-genome. We repeated the process for the next significant marker that has not been assigned to any group until all significant markers have been assigned. Lastly, we merged all overlapping groups.

We estimated the local heritabilities 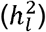 for each QSL in the TG panel using the genomic heritabilities partitioning method that was introduced by Schork (2001) and Visscher et al. (2007). QSLs with non-zero 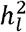 would support the hypothesis of selection over drift for the observed change in allele frequency. The TG panel includes 12 traits: Flowering Time (FT), Lodging (LODG), Yield (YLD), Plant Height (HT), Grain Protein Content (PROT), Winter Kill (WK), Awns (AWNS), Specific Weight (SPWT), Total Grain Weight (TGW), Ears per m^2^ (EM2), Tiller Number (TILL) and Maturity (MAT) (Bentley et al. 2014, Ladejobi et al. 2019). We were not able to estimate the 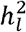 in the WAGTAIL panel since we did not have multi-trait data for the WAGTAIL panel. For each trait and QSL combination, we estimated the 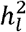 from the following mixed model fitted using the *mmer* function from the “sommer” package (Covarrubias-Pazaran et al. 2016) in R (R Core Team 2021):

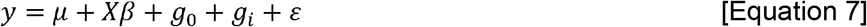

Where the model terms are described as below.

*y* is a vector of phenotypic trait values for *n* varieties.
*μ* is the trait mean.
*X* is an *n*×2 matrix of incidence matrix for the fixed year and country of origin effects.
*β* is a vector of length 2 of the fixed year and country of origin effects.
*g*_0_ is a vector of length *n* of the random genetic effect due to relationship among varieties calculated from markers not in group *i*, and it follows a distribution of 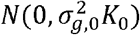.
*g_i_* is a vector of length *n* of the random genetic effect due to relationship among varieties calculated from markers in group *i*, and it follows a distribution of 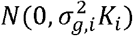.
*ε* is a vector of length *n* of the random residual effect under a distribution of 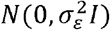.

After each model was fitted, we calculated the 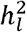 as 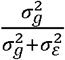. For any trait, we identified the non-zero 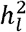 groups 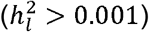 and refitted a new mixed model with all the non-zero 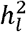 groups. The model is shown as below with the similar terms as explained in Equation 7.

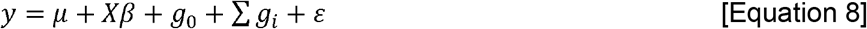

From Equation 5, we estimated the new 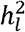 as 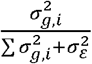 and used these as the final estimated 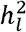 for each trait and group combination.

### Associating marker effects with alleles that are increasing over time

We estimated the marker allele effects for each trait in the TG panel using ridge regression (RR) (Hoerl and Kennard 1970) and least absolute shrinkage and selection operator (LASSO) (Tibshirani 1996) approaches. For the RR approach, we used the *mixed.solve* function from the “rrBLUP” package (Endelman 2011) in R (R Core Team 2021). For the LASSO approach, we used the *cv.glmnet* function from the ”glmnet” package (Friedman et al. 2010) in R (R Core Team 2021). In both approaches, we fitted a multiple linear regression model as shown below:

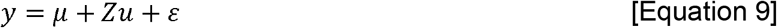

Where the model terms are described as below.

*y* is a vector of phenotypic trait values for *n* varieties.
*μ* is the trait mean.

*Z* is a *n*×*p* matrix of numerical marker genotypes coded as −1, 0 and 1 for homozygous first allele, heterozygous and homozygous second allele, respectively. The number of markers is *p.*

*u* is a vector of marker allele effects. In RR, *u* is estimated from minimizing the loss function of *L_RR_*(*u*) = ||*y* – *μ* – *Zu*||^2^ + *λ*||*u*||^2^ where 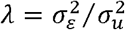 and 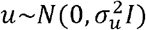 (Endelman 2011). In LASSO, *u* is estimated from minimizing the loss function of *L_LASSO_*(*u*) = ||*y* – *μ* – *Zu*||^2^ + *λ*||*u*|| where *λ* is determined from the default 10-fold cross validations in *cv.glmnet* (Friedman et al. 2010). In addition, the multivariate LASSO model in “glmnet” was used to ensure that the effects for all traits are estimated from the same set of chosen markers.

*ε* is a vector of residual effects that follows a distribution of 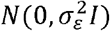.

For each trait *j* and marker *k*, we identified 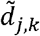 which is the effect direction for the allele that is increasing in frequency over time, as follows: first, we determined 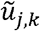 which is the direction of marker allele effect estimated from either RR or LASSO using the *sign* function in R (R Core Team 2021). This resulted in 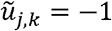 for negative effect, 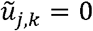 for no effect and 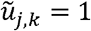 for positive effect. Next, we determined 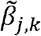 which is the direction of year regression coefficient estimated from RALLY. This resulted in 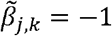 for decreasing allele and 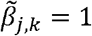 for increasing allele. Because the marker alleles were coded similarly in the RALLY and marker BLUP models, we could calculate 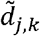 as 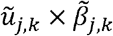 directly. 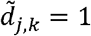 suggests that the increasing allele has a positive effect and 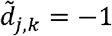 suggests that the increasing allele has a negative effect. 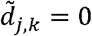 is only possible in LASSO due to variable selection, which simply implies that there is no effect. For any trait, an excess of either 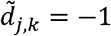 or 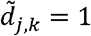 across all markers indicates a possible directional selection.

For a pair of traits *j*1 and *j*2, we calculated 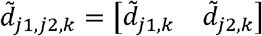 which is the pairwise effect direction for the increasing allele. 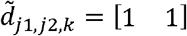 implies that the increasing allele has positive effects on both traits, 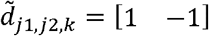 or 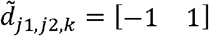 implies that the increasing allele has a positive and a negative effect on either trait, and 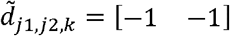 implies that the increasing allele has negative effects on both traits. By forming a contingency table from the counts of all four possible 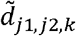 combinations, we tested for selection-related interaction between the pairs of traits using a 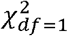 test in the results involving LASSO. We did not test the results involving RR because the marker effects are not independent.

### Estimating multivariate selection parameters

We estimated the multivariate selection parameters in the TG panel using the multivariate breeder’s equation of Δ*Z* = *Gβ_sel_* (Lande and Arnold 1983). We obtained the selection response (Δ*Z*), genetic variance-covariance matrix (*G*) and phenotypic variance-covariance matrix (*P*) from the trait and marker data. Next, we solved the multivariate breeder’s equation for the selection gradient *β_sel_* and the equations of *S* = *Pβ_sel_* and 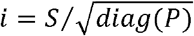 for the selection differential (*S*) and selection intensity (*i*) (Falconer and Mackay 1996). Lastly, we decomposed the multivariate selection parameters into direct and indirect partitions as a method to quantify the direct and indirect historical selection in the TG panel. As a check, we repeated the same process in a simulated example. Complete details on the methods on estimating multivariate selection parameters are provided in the Supplementary Methods.

## Results

### RALLY and GWAS in simulated populations

We tested RALLY’s ability in identifying selection- or drift-induced marker allele frequency changes in simulated populations with (S) and without (U) selection (Figure 1) by varying the degree of parametric control (PC). Briefly, PC combines genomic control (GC) (Devlin and Roeder 1999) and delta control (DC) (Gorroochurn et al. 2006) to correct for inflation in test statistics due to population structure. Details on the PC approach and simulations are described in the Materials and Methods section. Across all tested allele frequency change thresholds (*t*) for null marker set, setting t > 0.11 produced better control of test statistics (significant markers in S < 1.867%, U < 0.109%) than without any correction (significant markers in S = 1.942%, U = 0.089%) (Figure 2A, Table S1). At t = 0.15, we found little significance in the unselected population across all 100 simulations with some inevitable loss of significance in the selected population (significant markers in S = 0.994%, U = 0.012%) (Table S1). This result suggests that PC at this threshold can reasonably separate out the true selection signals from drift in our simulation. To err on the cautious side, we used a higher threshold of t = 0.20 in the simulation, TG and WAGTAIL panels.

**Figure 2.**
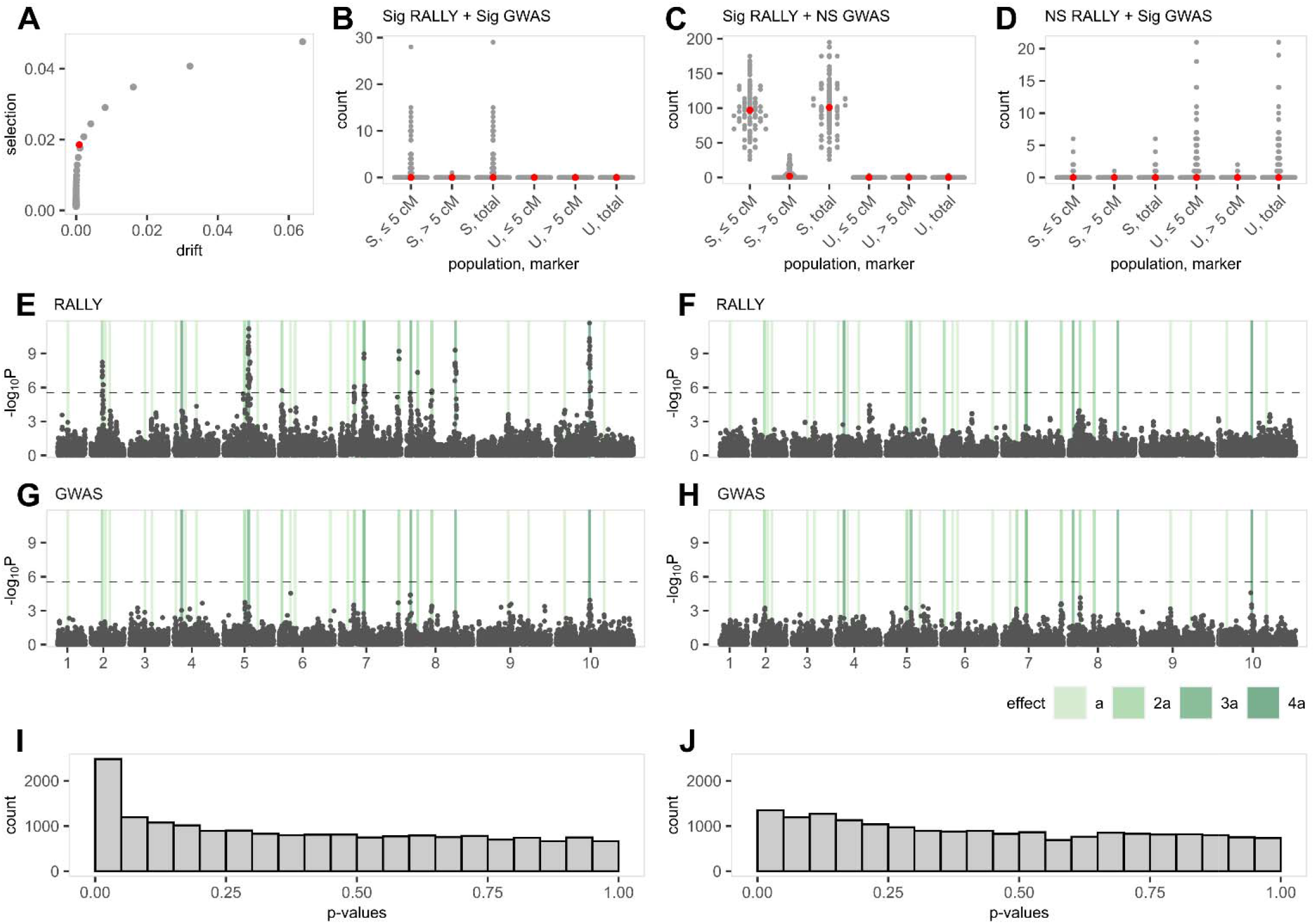
RALLY and GWAS in simulated populations. Selected (S) and unselected (U) populations are simulated for 100 times and mapped for QSLs/QTLs using RALLY and GWAS. [**A**] Significant proportions of total markers identified from RALLY in S and U populations are calculated under various thresholds used in choosing null markers for delta control (DC) and genomic control (GC). The red point is estimated without DC and GC (uncorrected). Under the assumption that significant markers in U are due to drift alone and in S are due to both drift and selection, the X-axis is shown as the proportions in U while the Y-axis is shown as the differences in proportions between S and U. [**B – D**] Counts of significant markers identified from RALLY and GWAS are shown according to their distance from QTLs in both S and U populations. Medians are shown in red points. [**E**] Manhattan plot for RALLY in one simulated S population. QTLs are highlighted in vertical bars according to their effect sizes. [**F**] Manhattan plot for RALLY in one simulated U population. [**G**] Manhattan plot for GWAS in one simulated S population. [**H**] Manhattan plot for GWAS in one simulated U population. [**I**] Histogram of RALLY p-values for the same simulated S population. [**J**] Histogram of RALLY p-values in the same simulated U population.

We evaluated the QSL/QTL mapping performances of RALLY and GWAS in the simulated populations with selection (Figure 1) and found a higher mapping power in RALLY over GWAS (Figure 2). Across the 100 simulations, we found that the individual significant markers are rarely shared between RALLY and GWAS (Figure 2B), and even less likely to be found in GWAS but not RALLY (Figure 2D). Most of the significant markers are found in RALLY but not GWAS (Figure 2C). The low number of significances in GWAS is likely because the simulated QTLs have small effects and low heritabilities, which is common for quantitative traits. The heritabilities for the largest QTLs are approximately 0.030 and the smallest QTLs are approximately 0.002. An additional intention of having low heritabilities is to reduce the fixation rate of QTL due to selection and prevent pre-matured fixation of QTL in the simulated population.

We repeated the RALLY and GWAS analyses in the unselected populations as a control for the same analyses in the selected populations (Figure 2). On average across all 100 simulations, RALLY identifies 0.1 significant markers out of 19,000 total markers in the unselected population compared to 104.5 significant markers in the selected population. This result suggests that less than 0.1% of the significant markers in the selected population are likely caused by drift instead of selection. In the selected population, there are more significant markers (means of 99.4 versus 5.1) that are close to the QTLs (≤ 5 cM) than far (> 5 cM) (Figure 2B-C). Assuming that all 32 QTLs are selected and all markers within 5 cM of the QTLs experience hitch-hiking effect, there should be a maximum of 3,200 significant markers in the selected population. However, the number of significant markers is much lower in reality because: (1) the selection force is proportional to the QTL effects (Figure S3), (2) the hitch-hiking effect depends on the initial marker-QTL haplotype distribution (Figure S1), and (3) the hitch-hiking effect decreases as genetic distance increases. On the other hand, GWAS performance remains similar between the selected and unselected populations (Figure 2).

### Detection limits of RALLY

Following from the previous simulation, we investigated the relationship between QTL under selection and its proximal markers and the results suggested a detection limit of approximately 5 cM (Figure S4). Here, we considered 10 markers that are evenly spaced between 1 to 10 cM away from a QTL and evaluated how these marker allele frequencies change as a result of increasing QTL frequency. Because the markers are linked to the QTL, we expect their frequencies to follow the QTL frequency in an inversely proportional way according to their genetic distances from the QTL. This process is commonly known as hitch-hiking, and it is an important consideration for RALLY because hitch-hiking markers are more likely to be genotyped than the true QTLs. Curiously, our results suggest that the ability of RALLY in identifying significant hitch-hiking markers depends on the QTL-marker haplotypes, QTL initial frequency, and genetic distance between QTL and marker (Figure S4). With all factors considered, RALLY rarely detects significance beyond 5 cM although our previous results showed that some long-range significances may still be present (Figure 2B). A possible explanation for this is when multiple QTLs co-localize into one major QTL haplotype, which may amplify the significances of surrounding markers.

### RALLY in two wheat panels

We mapped 22 significant QSLs (Bonferroni corrected p < 0.05) across 14 chromosomes in the Triticeae Genome (TG) panel using RALLY (Table 1, Figure 3, Figure S5, File S1). Because the distances between significant markers and true QTLs are unknown, we used a linkage disequilibrium (LD) measure of *r*^2^ > 0.2 as a method to identify the genomic boundaries that the significant markers tag. This method resulted in QSL intervals ranging from 1.46 Mb to 774.73 Mb with a mean of 148.74 Mb. Given the large blocks of genomic regions and a previously approximated RALLY detection limit of 5 cM, many of the QSLs are likely to fall within low recombination regions. QSLs in high recombination regions are harder to map due to the lack of markers tagging the causative QTLs. Besides, sustained selection is more likely to be observed on multiple weakly favorable alleles in low than high recombination regions.

**Figure 3.**
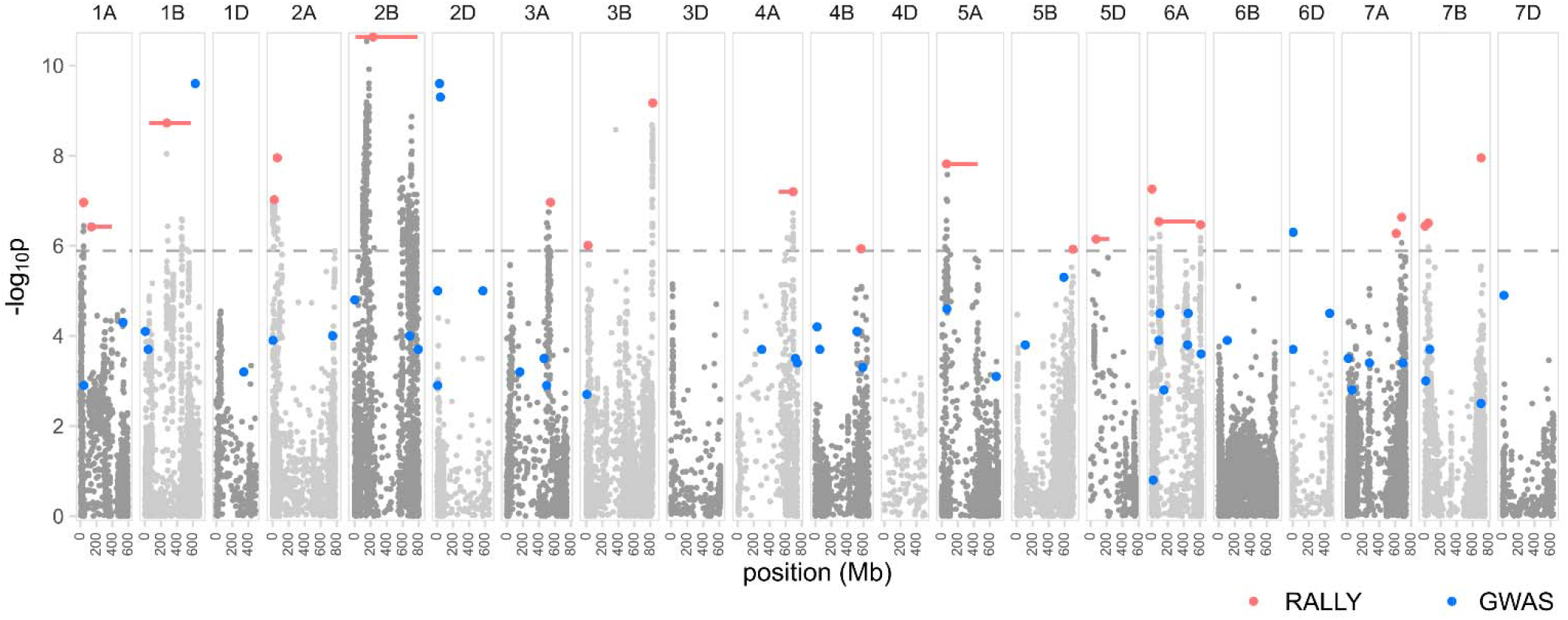
Manhattan plot for RALLY results in the TG panel. RALLY peaks and their extents of LD are shown in red points and horizontal bars, respectively. GWAS peaks from Ladejobi et al. (2019) are shown in blue points. The dashed horizontal line represents the Bonferroni threshold of 0.05.

**Table 1.** Genomic positions of 22 RALLY QSLs in the TG panel. The IWGSC RefSeq v1.0 physical positions of the peaks and LD boundaries are shown, along with the −log_10_P scores associated with the peaks and overlapping QSLs/QTLs from RALLY in WAGTAIL panel and GWAS in TG panel (Ladejobi et al. 2019).

| QSL | Chr | Position (bp) | | | $-\log_{10}P$ | Overlapping QSLs/QTLs | |
| --- | --- | --- | --- | --- | --- | --- | --- |
|  |  | Peak | Start | End |  | WAGTAIL | GWAS |
| 1 | 1A | 40,535,368 | 36,797,100 | 42,151,448 | 6.964 | 2 | 1 |
| 2 | 1A | 138,028,803 | 105,888,613 | 395,485,807 | 6.419 | 2 | 0 |
| 3 | 1B | 274,240,580 | 52,771,404 | 572,972,254 | 8.726 | 3 | 0 |
| 4 | 2A | 19,049,803 | 502,328 | 36,134,675 | 7.022 | 5 | 7 |
| 5 | 2A | 56,159,824 | 56,159,824 | 115,431,692 | 7.951 | 6 | 0 |
| 6 | 2B | 230,348,363 | 10,888,962 | 785,614,875 | 10.634 | 9,10 | 10 |
| 7 | 3A | 544,972,180 | 488,305,885 | 574,586,196 | 6.965 | 13 | 19 |
| 8 | 3B | 20,035,143 | 19,086,765 | 31,366,713 | 6.011 | 0 | 0 |
| 9 | 3B | 829,382,536 | 813,333,316 | 829,954,621 | 9.170 | 0 | 0 |
| 10 | 4A | 690,425,855 | 507,739,498 | 695,893,542 | 7.199 | 14 | 22 |
| 11 | 4B | 570,537,081 | 507,170,910 | 593,797,914 | 5.934 | 0 | 26,27 |
| 12 | 5A | 59,666,472 | 31,088,127 | 449,788,941 | 7.816 | 16 | 28 |
| 13 | 5B | 703,651,326 | 681,349,598 | 703,858,824 | 5.923 | 0 | 0 |
| 14 | 5D | 69,776,655 | 43,408,942 | 233,674,405 | 6.148 | 0 | 0 |
| 15 | 6A | 2,160,664 | 684,328 | 5,113,555 | 7.259 | 0 | 32 |
| 16 | 6A | 89,355,276 | 61,817,777 | 545,399,189 | 6.538 | 18 | 33 - 37 |
| 17 | 6A | 609,106,971 | 596,590,923 | 617,255,792 | 6.468 | 0 | 38 |
| 18 | 7A | 612,599,663 | 610,209,166 | 612,599,663 | 6.272 | 0 | 0 |
| 19 | 7A | 681,696,004 | 669,820,116 | 695,003,193 | 6.632 | 0 | 0 |
| 20 | 7B | 3,693,110 | 3,366,069 | 4,826,131 | 6.438 | 0 | 0 |
| 21 | 7B | 43,221,041 | 40,293,564 | 58,886,832 | 6.503 | 0 | 48 |
| 22 | 7B | 704,838,082 | 698,229,993 | 707,941,517 | 7.952 | 0 | 49 |

Of the 22 QSLs, 12 co-localize with previously mapped QTLs using GWAS (Ladejobi et al. 2019) in the TG panel (Table 1, Figure 3, Figure 4, Table S2). QSLs/QTLs found in both RALLY and GWAS indicate that their effects are likely beneficial and have been selected during the breeding process. QSLs unique to RALLY suggest that their effects might be too small for GWAS to detect or the specific traits have not been analyzed for GWAS. QTLs unique to GWAS suggest that they are still segregating in the population, which could be due to various reasons like recent introduction into the breeding population and linkage drag.

**Figure 4.**
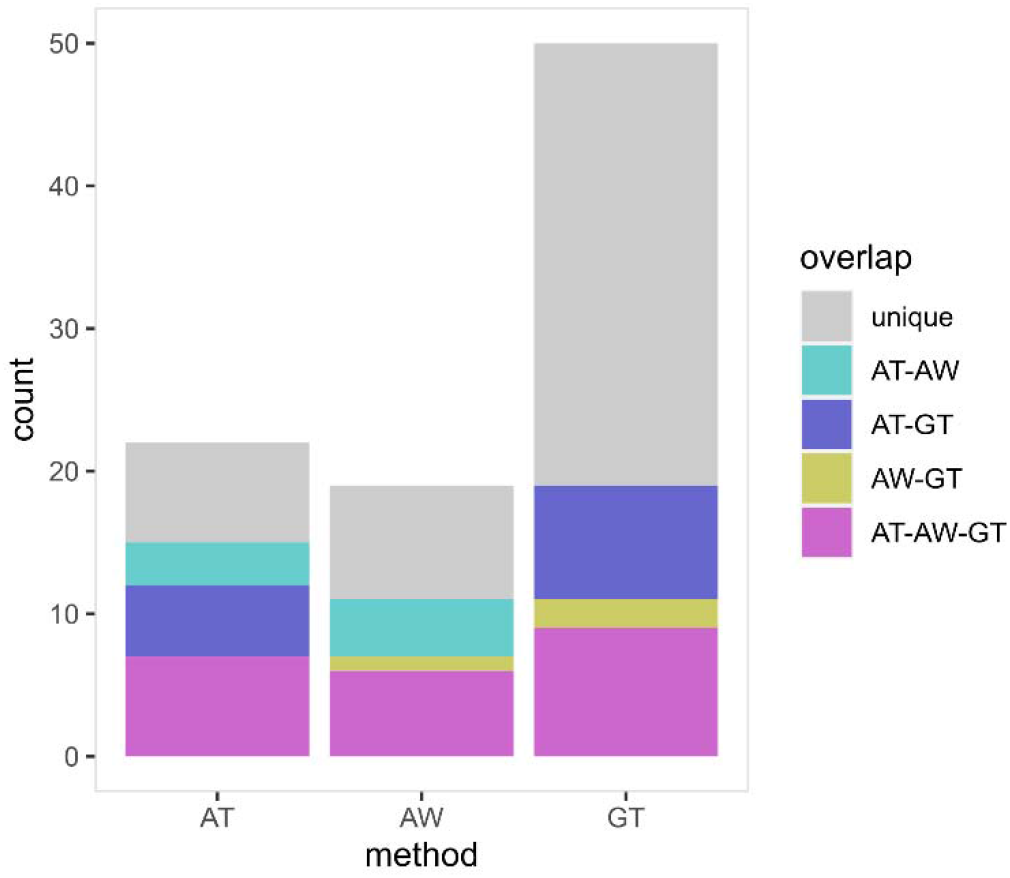
QSL/QTL overlaps across different results. The number of overlapping QTLs among RALLY in TG panel (AT), RALLY in WAGTAIL panel (AW) and GWAS in TG panel (GT) are shown.

A literature search showed that RALLY QSLs occur in both well-characterized and novel genomic regions in winter wheat (Table S3). The most significant RALLY QSL-6 mapped to a large region in chromosome 2B: 11 – 230 Mb, which includes *Ppd-B1* (Mohler et al. 2004) and multiple resistance loci of *Yr5, Yr7* and *YrSP* (Marchal et al. 2018). Another major QSL-16 mapped to a large region in chromosome 6A: 62 – 545 Mb, which contains *TaGW2* (Su et al. 2011) and the GA-responsive dwarfing genes of *Rht24* (Würschum et al. 2017) and *Rht25* (Mo et al. 2018). Interestingly, the durum wheat dwarfing gene *Rht14/16/18* resides in the same genomic region, although it remains to be tested whether it is allelic to *Rht24* (Haque et al. 2011). A recent EnvGWAS in winter wheat by Sharma et al. (2021) also mapped to the same genomic region (6A: 396 Mb) but without mention of any *Rht* candidate gene. On a broader scale, 16 RALLY QSLs co-localize with the recently identified meta-QTLs on yield and yield-related traits in wheat (Yang et al. 2021). 9 RALLY QSLs overlap with the QTLs identified from a Multi-parental Advanced Generation Inter-Cross (MAGIC) population of 16 diverse UK winter wheat varieties (Scott et al. 2021).

In addition, we found 11 RALLY QSLs that overlap with known alien and non-alien introgressions in wheat (Cheng et al. 2019). These include major introgressions like the 2A: 0 – 11 Mb from *Aegilops ventricosa* (Robert et al. 1999, Rhoné et al. 2007) and 2B: 90 – 749 Mb from *Triticum timopheevii* (Tsilo et al. 2008, Martynov et al. 2018). These two introgressions were shown to segregate among the UK winter wheat varieties by Scott et al. (2021). Because alien introgressions tend suppress recombination (Gill et al. 2011), they can be easily mapped using RALLY. Considering all overlaps in results between RALLY and the studies described thus far, we found 19 RALLY QSLs that can be traced to at least one study.

In the WAGTAIL panel, we mapped 19 significant QSLs across 13 chromosomes using RALLY (Table S4, Figure S6, File S1). We used the same approach as we did with the TG panel to identify the boundaries of these significant QSLs. With 99 varieties in common between the TG and WAGTAIL panels, we expect a high number of overlapping QSLs. 10 out of 19 QSLs in the WAGTAIL panel matched with 10 out of 22 QSLs in the TG panel (Figure 4), which is approximately one-half overlap between them. Given that the TG panel was genotyped using GBS (Elshire et al. 2011) while the WAGTAIL panel was genotyped using the 90k array (Wang et al. 2014), the genotyping and mapping quality of these two panels are likely different. This may partially explain why the results from the TG and WAGTAIL panels did not fully overlap. Another possible reason is that the distributions of countries of origins differ in the two panels in which the TG panel is more homogeneous than the WAGTAIL panel.

### Local heritabilities in the RALLY QSLs

We calculated local heritabilities for the 22 RALLY QSLs as a support for possible selection over drift at these QSLs (Table 2, Figure 5). We found that all 22 QSLs have non-zero local heritabilities for at least one trait. We tested for non-zero in the local heritabilities using a likelihood ratio test to compare between the mixed models with and without QSLs (Santantonio et al. 2019). However, most of the tests were non-significant due to low power (Table S5). The tests for QSLs collectively showed significance in 5 out of 12 traits, which comes at a cost of losing the test on individual QSL in exchange for a slightly higher power. In an extreme example with a total heritability of 0.379, QSL-16 at 6A: 89,355,276 is associated with 8 traits and found to co-localized with all other previously mentioned results. While it is possible that the underlying candidate genes *TaGW2* (Su et al. 2011), *Rht24* (Würschum et al. 2017) and *Rht25* (Mo et al. 2018) have pleiotropic effects that are beneficial for wheat breeding, we cannot exclude the possibility of additional genes that provide breeding advantages in the same haplotype block. Nonetheless, given that QSL-16 has already played a major role in wheat breeding, it is unlikely to be useful for future breeding. The genomic region with the next largest total heritability of 0.226 is located in QSL-2 at 1A: 138,028,803. While no known gene has been mapped around QSL-2, results from our analysis and others (Cadalen et al. 1998, Griffiths et al. 2012, Tiwari et al. 2016) suggest that it may contain loci responsible for plant height and grain protein content.

**Figure 5.**
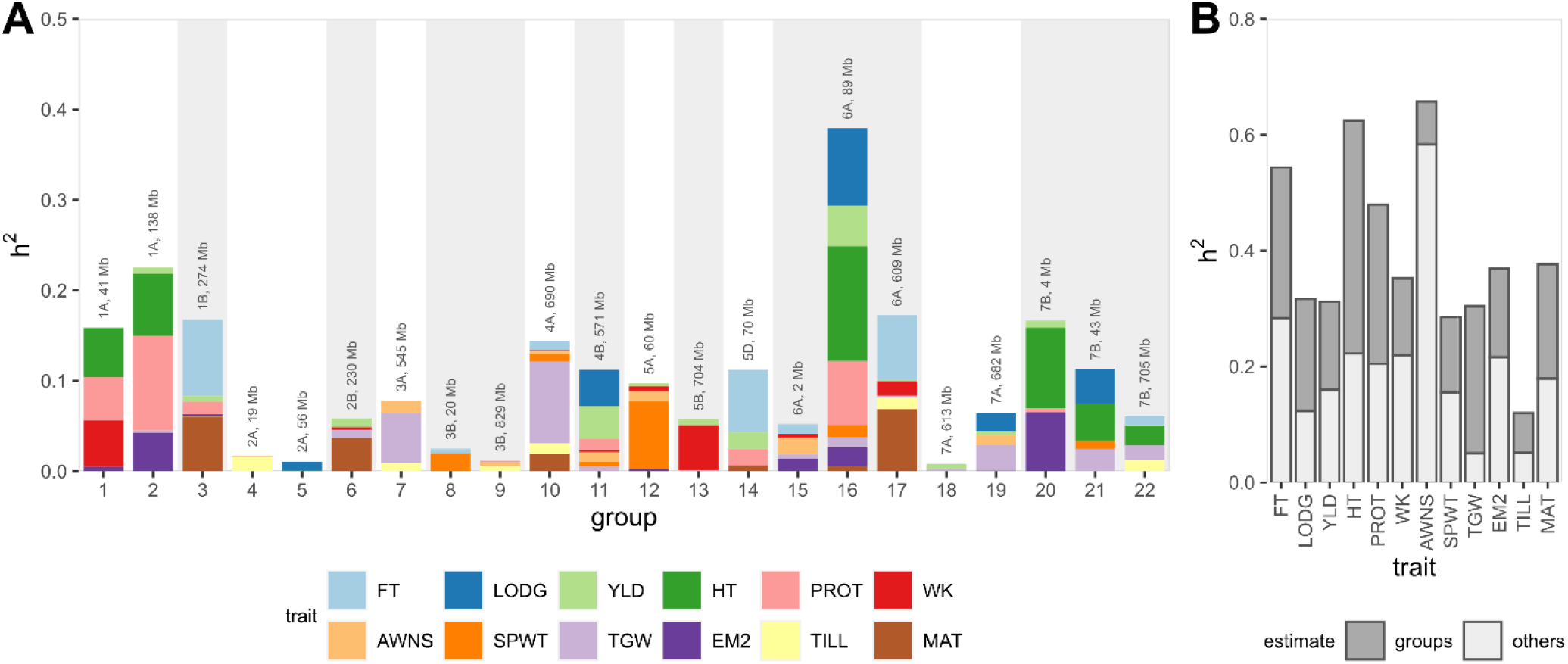
Local heritabilities in RALLY QSL groups. [**A**] Local heritabilities for all 12 traits are shown as stacked bars for each RALLY QSL (defined in Table 1). [**B**] Local heritabilities from 22 RALLY groups are summed and compared against the heritabilities from other genomic markers.

**Table 2.**
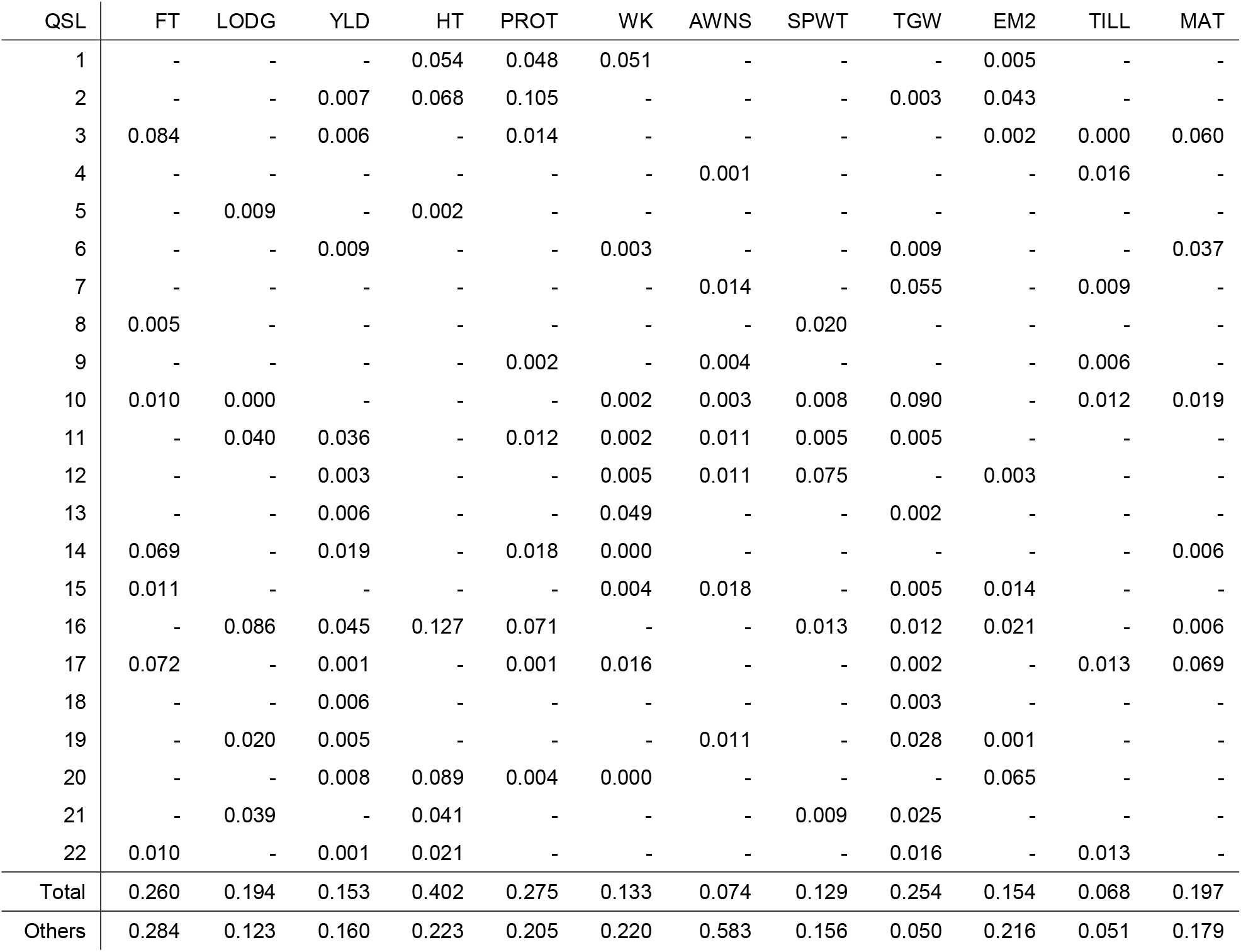
Local heritabilities associated with 22 RALLY QSLs. Local heritabilities less than 0.001 are not shown.

| QSL | FT | LODG | YLD | HT | PROT | WK | AWNS | SPWT | TGW | EM2 | TILL | MAT |
| --- | --- | --- | --- | --- | --- | --- | --- | --- | --- | --- | --- | --- |
| 1 | - | - | - | 0.054 | 0.048 | 0.051 | - | - | - | 0.005 | - | - |
| 2 | - | - | 0.007 | 0.068 | 0.105 | - | - | - | 0.003 | 0.043 | - | - |
| 3 | 0.084 | - | 0.006 | - | 0.014 | - | - | - | - | 0.002 | 0.000 | 0.060 |
| 4 | - | - | - | - | - | - | 0.001 | - | - | - | 0.016 | - |
| 5 | - | 0.009 | - | 0.002 | - | - | - | - | - | - | - | - |
| 6 | - | - | 0.009 | - | - | 0.003 | - | - | 0.009 | - | - | 0.037 |
| 7 | - | - | - | - | - | - | 0.014 | - | 0.055 | - | 0.009 | - |
| 8 | 0.005 | - | - | - | - | - | - | 0.020 | - | - | - | - |
| 9 | - | - | - | - | 0.002 | - | 0.004 | - | - | - | 0.006 | - |
| 10 | 0.010 | 0.000 | - | - | - | 0.002 | 0.003 | 0.008 | 0.090 | - | 0.012 | 0.019 |
| 11 | - | 0.040 | 0.036 | - | 0.012 | 0.002 | 0.011 | 0.005 | 0.005 | - | - | - |
| 12 | - | - | 0.003 | - | - | 0.005 | 0.011 | 0.075 | - | 0.003 | - | - |
| 13 | - | - | 0.006 | - | - | 0.049 | - | - | 0.002 | - | - | - |
| 14 | 0.069 | - | 0.019 | - | 0.018 | 0.000 | - | - | - | - | - | 0.006 |
| 15 | 0.011 | - | - | - | - | 0.004 | 0.018 | - | 0.005 | 0.014 | - | - |
| 16 | - | 0.086 | 0.045 | 0.127 | 0.071 | - | - | 0.013 | 0.012 | 0.021 | - | 0.006 |
| 17 | 0.072 | - | 0.001 | - | 0.001 | 0.016 | - | - | 0.002 | - | 0.013 | 0.069 |
| 18 | - | - | 0.006 | - | - | - | - | - | 0.003 | - | - | - |
| 19 | - | 0.020 | 0.005 | - | - | - | 0.011 | - | 0.028 | 0.001 | - | - |
| 20 | - | - | 0.008 | 0.089 | 0.004 | 0.000 | - | - | - | 0.065 | - | - |
| 21 | - | 0.039 | - | 0.041 | - | - | - | 0.009 | 0.025 | - | - | - |
| 22 | 0.010 | - | 0.001 | 0.021 | - | - | - | - | 0.016 | - | 0.013 | - |
| Total | 0.260 | 0.194 | 0.153 | 0.402 | 0.275 | 0.133 | 0.074 | 0.129 | 0.254 | 0.154 | 0.068 | 0.197 |
| Others | 0.284 | 0.123 | 0.160 | 0.223 | 0.205 | 0.220 | 0.583 | 0.156 | 0.050 | 0.216 | 0.051 | 0.179 |

Between the cumulative heritabilities explained by these 22 QSLs and the remaining genomic regions, HT and TGW are higher in the QSLs, AWNS is lower in the QSLs and the other 9 traits are about equal (Table 2, Figure 5). This result highlights the narrow genetic diversity that is often seen in modern varieties (Reif et al. 2005) due to the repeated use of identical favorable haplotypes in wheat breeding. Fortunately, the remaining “unselected” genomic regions for important traits like yield, grain protein content and plant height are not fully devoid of heritabilities. There is still room for varietal improvement without the introduction of favorable exotic alleles in the short term, which suggests that it might be better to devote some of the resources in pre-breeding on these genomic regions instead. For traits like TGW and TILL, breeders may need to look for alternative genetic resources to compensate for the lack of diversity.

### Marker effects of alleles that are increasing over time

We evaluated the marker allele effects using the prediction models from Ridge Regression (RR) and Least Absolute Shrinkage and Selection Operator (LASSO). Across all 12 traits, RR resulted in higher prediction accuracy than LASSO although the differences were comparable in some traits (Figure S7 and S8). Despite that, we retained the results from both approaches because the variable selection step in LASSO is important for a follow-up test involving trait pairs.

We examined the marker allele effect directions for increasing alleles and found excesses in one over another direction across each of the 12 traits (Figure 6, Figure S9). We first partitioned the markers based on their RALLY significance into three groups: (1) markers with p-values lower than the Bonferroni corrected threshold of 0.05, (2) markers with p-values between 0.05 and the Bonferroni corrected threshold of 0.05, and (3) markers with p-values higher than 0.05. The results from using either RR (Figure 6) or LASSO (Figure S9) are similar although the differences across the significance groups in LASSO are less pronounced, i.e. there are more differences between group 1 and 2 in RR than LASSO results. This might be due to LASSO selected markers having weak but small, non-significant changes in allele frequencies over time. Within the RR results, the excesses in effect directions are strongest in the significance group 1 and weakest in the significant group 3, which suggest that the excesses can be related to the favored direction of selection. The lack of excesses in significance group 3 implies that favorable and unfavorable alleles are still segregating about equally in the unselected genomic regions.

**Figure 6.**
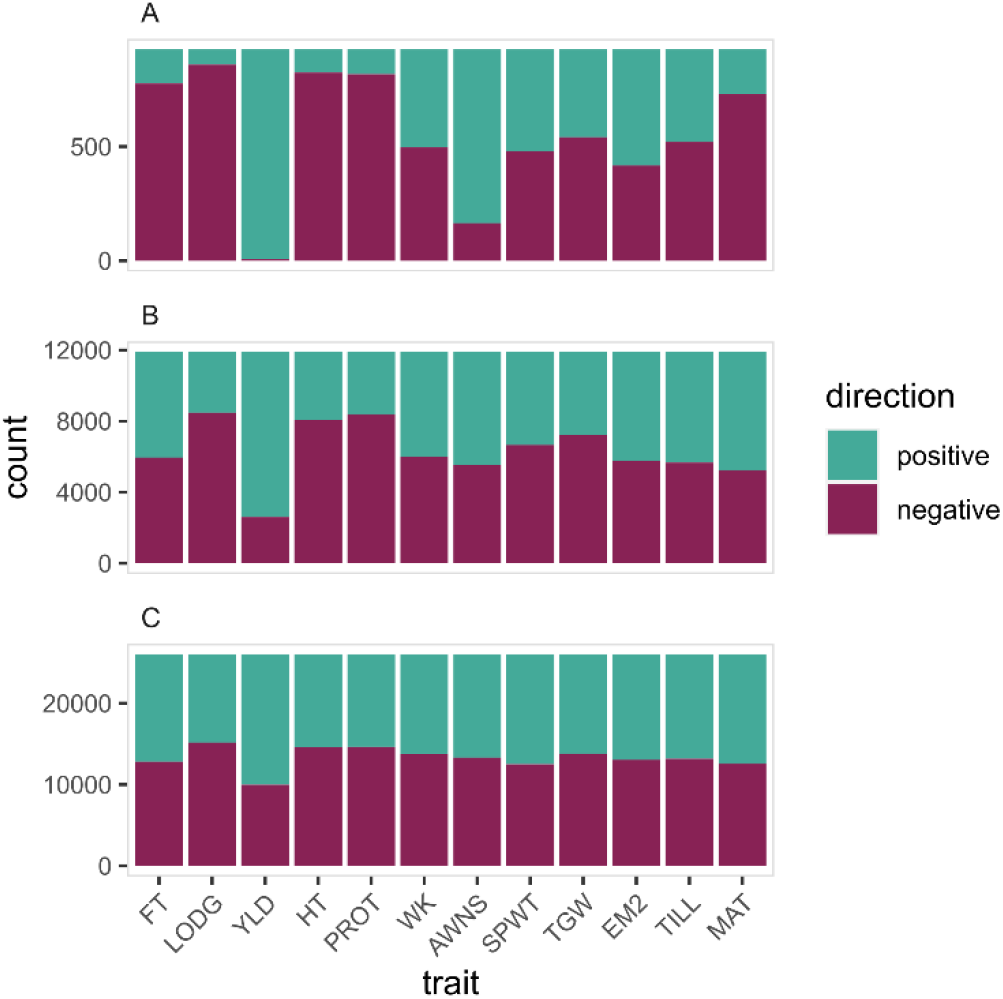
Distributions of positive and negative RR effects in the increasing alleles. [A] Markers with RALLY P-values of lower than the Bonferroni corrected threshold. [B] Markers with RALLY P-values between 0.05 and the Bonferroni corrected threshold. [C] Markers with RALLY P-values of higher than 0.05.

Across all 12 traits, the excesses agree with our expectation of traits that are important in wheat breeding. The most extreme example is yield (YLD) where both the RR and LASSO results show a near complete excess of positive effects in the increasing alleles in significance group 1. As shown previously by Mackay et al. (2011), the genetic gain in the UK winter wheat yield has been rising steadily over time. The next four traits with strong excesses are flowering time (FT), lodging score (LODG), plant height (HT) and grain protein content (PROT). FT, LODG and HT are favored for lower trait values, and thus the increasing alleles have excesses in negative effects. On the contrary, higher PROT is valuable for bread making quality, which is unfortunately going in the opposite direction due to a strong negative genetic correlation with yield (Scott et al. 2021). This result suggests that the selection for higher yield is a lot stronger than the selection for higher grain protein content. In the remaining traits, the excesses are smaller and less obvious given the variations seen from RR and LASSO results, which suggests that directional selection is likely weak for these traits.

By comparing the effect directions for increasing alleles in pairs of traits, we identified the priorities of traits under selection (Table 3, Table S6 and S7, Figure 7). Taking YLD and PROT for example, there is a strong excess for alleles with positive YLD but negative PROT. This result reiterates the priority of YLD over PROT in wheat breeding. Between TGW and EM2, there is an excess for alleles with positive EM2 and negative TGW which suggests that more ears with lighter grains are preferred over fewer ears with heavier grains. In a different perspective, the results here also highlight the constraints imposed by genetic correlations across traits. For example, there is a small proportion of alleles with the same effect directions for YLD and PROT. These alleles could be used in breeding high YLD and PROT varieties, although it is still important to consider the possibility that these alleles could be unfavorably associated with other traits.

**Figure 7.**
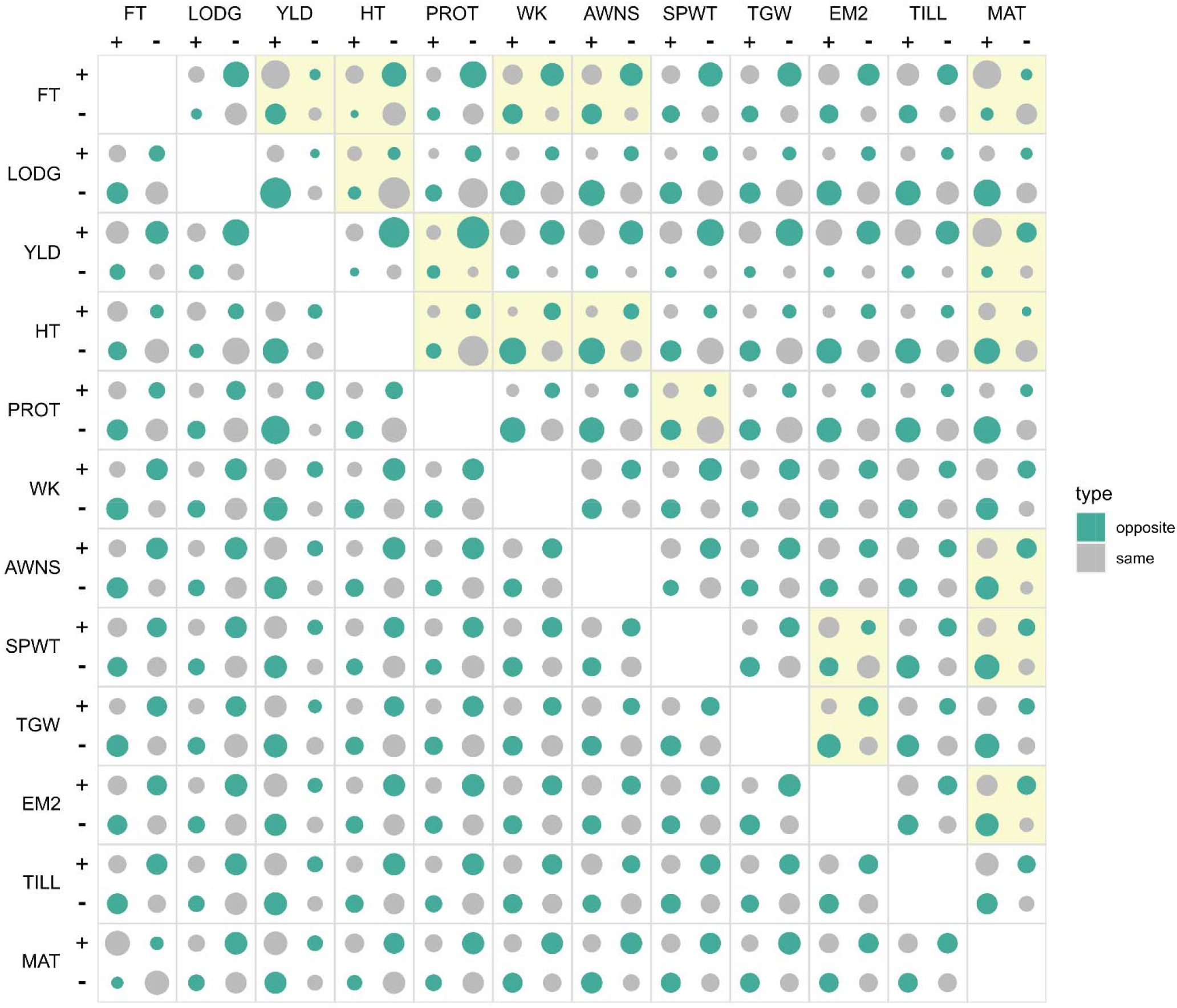
Distribution of pairwise effects for alleles with increasing frequency over time. For each allele with increasing frequency over time, it is classified into pairs of traits for which the allele has an increasing effect on both traits (+/+), a decreasing effect on both traits (-/-) or antagonistic/opposite effects (+/− and -/+). The circle areas are scaled according to the marker counts. The bottom left triangle represents the RR effects and the top right triangle represents the LASSO effects. The distributions of the effect classes in each trait pair are tested using in the LASSO effects and significant results (Bonferroni-corrected threshold of p = 0.05) are highlighted in yellow. No test is performed in the RR effects because the marker effects in RR are not independent.

**Table 3.**
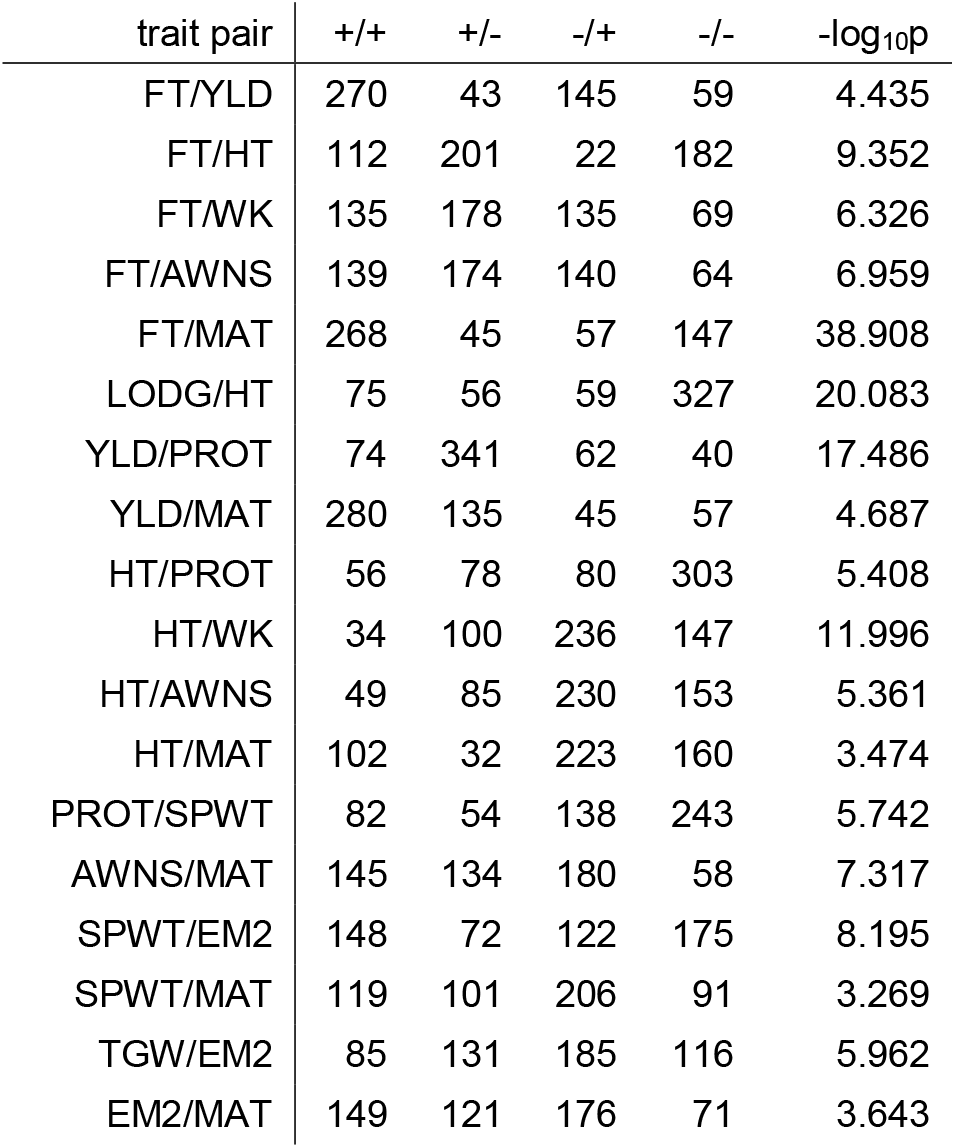
Counts of pairwise LASSO effects for alleles that are increasing over time. For each allele with increasing frequency over time, it is classified into pairs of traits for which the allele has an increasing effect on both traits (+/+), a decreasing effect on both traits (-/-) or antagonistic effects (+/− and -/+). The distribution for each pair of traits is tested with a 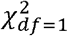 contingency table where the significant threshold is set to −log_10_p = 3.121 (equivalent to a Bonferroni-corrected threshold of p = 0.05 for 66 possible pairs of traits). Only the significant trait pairs are shown here, and the full results are available in Table S7.

| trait pair | +/+ | +/- | -/+ | -/- | $-\log_{10}p$ |
| --- | --- | --- | --- | --- | --- |
| FT/YLD | 270 | 43 | 145 | 59 | 4.435 |
| FT/HT | 112 | 201 | 22 | 182 | 9.352 |
| FT/WK | 135 | 178 | 135 | 69 | 6.326 |
| FT/AWNS | 139 | 174 | 140 | 64 | 6.959 |
| FT/MAT | 268 | 45 | 57 | 147 | 38.908 |
| LODG/HT | 75 | 56 | 59 | 327 | 20.083 |
| YLD/PROT | 74 | 341 | 62 | 40 | 17.486 |
| YLD/MAT | 280 | 135 | 45 | 57 | 4.687 |
| HT/PROT | 56 | 78 | 80 | 303 | 5.408 |
| HT/WK | 34 | 100 | 236 | 147 | 11.996 |
| HT/AWNS | 49 | 85 | 230 | 153 | 5.361 |
| HT/MAT | 102 | 32 | 223 | 160 | 3.474 |
| PROT/SPWT | 82 | 54 | 138 | 243 | 5.742 |
| AWNS/MAT | 145 | 134 | 180 | 58 | 7.317 |
| SPWT/EM2 | 148 | 72 | 122 | 175 | 8.195 |
| SPWT/MAT | 119 | 101 | 206 | 91 | 3.269 |
| TGW/EM2 | 85 | 131 | 185 | 116 | 5.962 |
| EM2/MAT | 149 | 121 | 176 | 71 | 3.643 |

### Multivariate selection parameters

In contrast to a genomic-centric approach that has been described thus far, the multivariate selection parameters may provide an alternative, trait-focused perspective on the historical selection of winter wheat represented by the TG panel. We found a strong misalignment between the selection response (ΔZ) and gradient (β_sel_) where the directions of the vectors’ elements are the opposite in 5 out of 12 traits (Table S8). If the selection parameters are estimated accurately, such divergence may imply an inefficient selection process. In addition, we partitioned the selection response (ΔZ), differential (S) and intensity (i) into direct and indirect components to quantify the amount of each selection parameter that is directly due to the available variation within a trait or indirectly due to the covariation with other traits. In an example with HT, we found positive direct effects in ΔZ, S and i, which contradicts the known selection on dwarfing genes like *Rht1, Rht2* and *Rht24* (Pearce et al. 2011, Würschum et al. 2017). Given the uncertainties in the multivariate selection parameters, we have provided the full results in the Supplementary Results and we advise to treat these estimates with caution.

Following the results, we investigated the possible causes of issues in estimating multivariate selection parameters using a simulated example with a single generation of selection involving three genetically correlated traits. First, we found that the genetic variances and covariances (G) estimated from mixed linear model were close to the true simulated values but with low precision (Table S9, Figure S10). Next, we computed the selection parameters (ΔZ, β_sel_, S, i) from the simulation directly, true G and estimated G, which are referred to as true, realized and estimated values, respectively. Given the imprecise estimates of G, we observed lower correlations between the estimated and true values than between the realized and true values (Table S9, Figure S11-S16). Despite using the true G, the realized values still failed to match the true values perfectly, which indicates that the deviations in realized ΔZ are carried over into the other selection parameters that are estimated downstream.

## Discussion

### Advantages and disadvantages of RALLY

RALLY has a major feature of being a trait-free method for mapping QSLs; however, this feature is a double-edged sword. For any population, RALLY involves only a single, relatively simple logistic regression analysis. In contrast, GWAS requires either multiple, simple mixed model analyses for each trait or a single, yet computationally intensive multi-trait analysis. Unlike any other trait-based mapping methods, the QSLs identified through RALLY are not restricted to only traits that are scored. While this makes RALLY a convenient method, the results do not inform us which traits the QSLs are associated with. In this regard, we will need to rely on other trait-based analyses like GWAS or genomic variance partitioning (Schork 2001, Visscher et al. 2007) to relate QSLs to traits. This additional step is not restricted to the same population as the QSL-trait information can be drawn from other studies such as GWAS on 47 traits in the wheat MAGIC diverse population (Scott et al. 2021). Therefore, RALLY can function as a replication of results from other studies.

As a kinship-free method, RALLY avoids any potential issues that may arise from the use of genomic relationship matrix (GRM) in mixed linear models. Recently, kinship estimates have been shown to be biased under complex population structure (Ochoa and Storey 2021), which can arise due to selection and migration of materials across breeders and countries. Besides, kinship estimates depend on the assumption that the alleles frequencies observed in the study population are representative of the reference or base population. For a population that has only experienced weak to no selection, the mean of genome-wide marker variance might be a reasonable approximation to the reference population. But, in populations under strong selection like modern crop varieties, the deviation between observed and true (reference) distribution of allele frequencies may not be trivial. Jiang et al. (2021) showed that the kinship estimates are biased when the observed distribution of allele frequencies fails to match the true distribution. In addition, a similar study on populations of modern wheat and barley varieties suggested that their kinship estimates may be biased due to long period of intensive selection (Sharma et al. 2021). However, the bias impacts on mapping power in GWAS and accuracy of variance component estimates remain to be evaluated.

Given that RALLY is designed specifically for mapping QSLs that have been selected over a time period, there may be limited utilities outside of its target scope. Our RALLY analyses model the change in allele frequency under a logistic distribution, which requires both genomic marker and year of variety release information. So, RALLY cannot be immediately applied to typical artificial mapping populations like bi-parental, nested association mapping (NAM) or MAGIC populations. However, we can extend the use of RALLY by conceptualizing it in its simplest form, which is a regression of marker allele on a variable of interest. For example, we can regress the marker allele on a continuous geographical origin variable such as latitudes and altitudes. The outcomes would directly define alleles that are relevant to local adaptation. Furthermore, the hybrid approach of parametric control (PC) is independent of RALLY and can be used in any genome-wide mapping analyses as a replacement for GRM and mixed linear model.

### Selection history and future direction in winter wheat breeding

Given the largely incomplete overlap between RALLY and GWAS QSLs/QTLs in the TG panel, GWAS-specific QTLs may not have been directly useful in breeding. Several possible reasons include linkage drag between the QTLs, recent introduction of QTL alleles into the breeding pool, and ineffective selection at those QTLs. In the absence of genome editing to remove unfavorable alleles (Johnsson et al. 2019), linkage drag is unavoidable due the low probability of creating favorable recombinant haplotypes. New QTL alleles are hard to map under RALLY due to low power issue, but it can be improved by including more recent varieties. Ineffective selection is a direct consequence of the selection tendency towards low-hanging fruits. In an extreme example involving a cross between an elite variety and an exotic wild relative, selection is bound to reconstitute the elite genome because of the higher probabilities of favorable alleles in the elite over exotic genomes (Gorjanc et al. 2016). This phenomenon is observed in a large-scale crossing program involving groups of one exotic and two elite parents, in which the resulting lines lost approximately two-thirds of the expected exotic genome (Singh et al. 2021). In this regard, the approach of Origin Specific Genomic Selection (OSGS) (Yang et al. 2020) can be used to specifically target genomic regions outside of RALLY QSLs for selection.

The association between directions of allele frequency change and predicted marker effects provides us with an overview of selection priorities (Figure 6 and 7). High yield, short plants, early flowering, reduced lodging and reduced grain protein content are clearly preferred under directional selection. However, there is no obvious directional selection on spikes and grain related traits, which suggests that there is no specific morphology that provides advantage in the breeding practice. The pairwise analysis further demonstrates the selection priorities and genetic correlations between traits. The results can be used to formulate a future breeding direction, for example, breeding for varieties with high yield and grain protein content by focusing on increasing the frequencies of the favorable alleles on both traits. In line with the global interest in shifting towards more sustainable agricultural practice (Hoad 2010), this approach can be extended to include traits relevant to sustainability and climate resilience to better guide the breeding direction.

### Limited practical use of multivariate breeder’s equation

As shown in the results involving the TG panel, the multivariate breeder’s equation has limited practical use in estimating selection parameters (Table S8). An important component of the equation is the genetic variance-covariance matrix (G). The assumption that G is constant is likely violated because G should have been calculated from the base population (Walsh and Lynch 2018) rather than a population under selection over a time period. While this violation likely contributes to the poor estimates of the selection parameters, it is not the only source of issue. Variations across two tested genotyping methods (GBS and DArT) resulted in severely different selection parameters (Table S8) even when the G were similar across the two methods (Table S10).

Despite fulfilling the assumption of constant G and eliminating the genotyping discrepancy in our simulated example, additional issues remain in estimating selection parameters from the multivariate breeder’s equation. We found that the poor estimation of multivariate selection parameters is caused by imprecise G estimated from mixed linear model. However, the estimation of multivariate selection parameters cannot be completely recovered even when the true G is used. This is probably because the multivariate breeder’s equation can only capture the means but not the variances of the selection parameters (ΔZ, β_sel_, S, i). Since the selection parameters are derived sequentially, repeated deviations from the means result in poor estimates of the selection parameters. This issue can be remedied by increasing the sample size, although there is a limit to the sample size due to practicality in breeding practice. Furthermore, the deviation is amplified across multiple generations of selection. Given the multi-layered issues with estimating selection parameters using the multivariate breeder’s equation, it is best to limit its use to predict forward for a single generation as a rough guide to selection experiments involving crop varieties.

## Acknowledgments

We thank Rajiv Sharma, Ian Dawson and David Marshall for helpful discussion on the work.

## Author Contributions

CJY and IM conceived the work, performed the analyses and wrote the manuscript. OL provided data for the Triticeae Genome (TG) panel. RM and WP provided critical comments. All authors read, revised and approved the manuscript. No external funding was received for the work in this manuscript.

## Data Availability

The GBS and phenotypic trait data for the TG panel (Ladejobi et al. 2019) were downloaded from doi.org/10.6084/m9.figshare.7350284. The TG panel DArT data (Bentley et al. 2014) and WAGTAIL panel data (Fradgley et al. 2019) were downloaded from https://www.niab.com/research/agricultural-crop-research/resources. The IWGSC RefSeq v1.0 annotation file containing the physical map positions for the 90k wheat array was downloaded from https://urgi.versailles.inra.fr/download/iwgsc/IWGSC_RefSeq_Annotations/v1.0/iwgsc_refseqv1.0_Marker_mapping_summary_2017Mar13.zip. Computational analyses were performed using R version 4.1.0. All R scripts used in the analyses can be found at https://cjyang-sruc.github.io/RALLY.

