## Supplementary Material for "Analysis of historical selection in winter wheat"

Chin Jian Yang*

Olufunmilayo Ladejobi†

Richard Mott†

Wayne Powell*

Ian Mackay*,‡

*Scotland's Rural College (SRUC), Kings Buildings, West Mains Road, Edinburgh, EH9 3JG, UK.

†Department of Genetics, Evolution & Environment, University College London, London, WC1E 6BT

‡IMplant Consultancy Ltd., Chelmsford, UK.

### Supplementary Methods

#### Estimating multivariate selection parameters

We calculated the genetic variance-covariance ($G$) matrix for all 12 traits in the TG panel GBS marker data (Ladejobi et al. 2019). The variances were estimated from a univariate model (Equation 8) while the covariances were estimated from a bivariate model (Equation 9). All models were fitted using the *mmer* function from "sommer" (Covarrubias-Pazaran 2016) in R (R Core Team 2021). The model terms are similar to Equation 7 except that $g$ follows a normal distribution of $N(0, \sigma_g^2 K)$ while $g_1$ and $g_2$ follow a multivariate normal distribution of $MVN(0, \begin{bmatrix} \sigma_{g1}^2 & \sigma_{g1,g2} \\ \sigma_{g1,g2} & \sigma_{g2}^2 \end{bmatrix} \otimes K)$. The diagonal terms $\sigma_{g1}^2$ and $\sigma_{g2}^2$ are the genetic variances for trait 1 and 2, and the off-diagonal term $\sigma_{g1,g2}$ is the genetic covariance between trait 1 and 2. The residual effect $\varepsilon_1$ and $\varepsilon_2$ follow a multivariate normal distribution of $MVN(0, \begin{bmatrix} \sigma_{\varepsilon 1}^2 & \sigma_{\varepsilon 1,\varepsilon 2} \\ \sigma_{\varepsilon 1,\varepsilon 2} & \sigma_{\varepsilon 2}^2 \end{bmatrix} \otimes I)$.

$$y = \mu + X\beta + g + \varepsilon \qquad \text{[Equation 10]}$$

$$\begin{cases} y_1 = \mu_1 + X\beta_1 + g_1 + \varepsilon_1 \\ y_2 = \mu_2 + X\beta_2 + g_2 + \varepsilon_2 \end{cases} \qquad \text{[Equation 11]}$$

Similar to above, we calculated the $G$-matrices for each country of origin ($G_{DE}$, $G_{FR}$, $G_{UK}$) and year group ($G_{y1}$, $G_{y2}$, $G_{y3}$) to test if $G$ is stable across geographical and temporal variations in the TG panel. We divided the panel into three approximately equal year groups by choosing varieties in 1990 or earlier as group 1, after 1990 but before 2002 as group 2, and 2002 or later as group 3. We used Random Skewers (RS) (Cheverud and Marroig 2007) as provided by the *skewers* function from the "phytools" package (Revell 2012) in R (R Core Team 2021) to test if any two $G$'s are different. Specifically, RS measures the correlations in their selection responses $\Delta Z$ when two $G$'s are multiplied by random selection gradients $\beta_{sel}$. We tested if $G_{DE}$, $G_{FR}$ and $G_{UK}$ are different from each other and also if they are different from $G$. Similar comparisons were applied to $G_{y1}$, $G_{y2}$, and $G_{y3}$. We also repeated all of the comparisons using the Mantel test (Mantel 1967) as provided by the *mantel.test* function from the ape" package (Paradis et al. 2004) in R (R Core Team 2021).

In addition, we calculated the genetic correlation ($g$) matrices by scaling all $G$ matrices by their diagonals (variances). We repeated all of the comparisons using RS and Mantel test in the $g$ matrices. The use of $g$ matrices eliminates any variation due to the differences in trait scales.

Next, we estimated the $G_{DArT}$ and $g_{DArT}$ from the TG panel DArT marker data (Bentley et al. 2014) as a check if $G$ and other selection parameters are sensitive to the choice of marker data. To do so, we applied the same RS and Mantel test to check between $G$ and $G_{DArT}$ as well as between $g$ and $g_{DArT}$.

As elucidated in the RS test, the multivariate breeder's equation of $\Delta Z = G\beta_{sel}$ (Lande and Arnold 1983) can be applied to estimate the selection gradient in the TG panel. Using a simple linear regression of trait $j$ as the dependent variable and the variety year of release and country of origin as independent variables, we obtained the year regression coefficient as an estimate for the annual selection response $\Delta Z$. Given $\Delta Z$ and $G$, we solved the multivariate breeder's equation and obtained the annual selection gradient $\beta_{sel}$. We also derived the phenotypic variance-covariance matrix ($P$) as a sum of genetic ($G$) and residual ($E$) variance-covariance matrix. This allowed us to solve the equation of $S = P\beta_{sel}$ and $i =$

$S/\sqrt{diag(P)}$ for the annual selection differentials $S$ (Falconer and Mackay 1996) and annual selection intensity $i$.

Additionally, we decomposed the selection responses $\Delta Z$ and differentials $S$ into direct and indirect components. This calculation is explained in the following equation for $\Delta Z$, which can be easily translated for $S$ and $i$.

$$\Delta Z = \begin{bmatrix} \Delta Z_1 \\ \Delta Z_2 \\ \vdots \\ \Delta Z_{12} \end{bmatrix} = G\beta_{sel} = \begin{bmatrix} G_{1,1} & G_{1,2} & \cdots & G_{1,12} \\ G_{2,1} & G_{2,2} & \vdots & G_{2,12} \\ \vdots & \cdots & \ddots & \vdots \\ G_{12,1} & G_{12,2} & \cdots & G_{12,12} \end{bmatrix} \begin{bmatrix} \beta_{sel,1} \\ \beta_{sel,2} \\ \vdots \\ \beta_{sel,12} \end{bmatrix} \quad \text{[Equation 12]}$$

Following Equation 9, trait 1 selection response can be written as $\Delta Z_1 = G_{1,1}\beta_{sel,1} + G_{1,2}\beta_{sel,2} + \cdots + G_{1,12}\beta_{sel,12}$. The direct response is $G_{1,1}\beta_{sel,1}$ due to selection gradient $\beta_{sel,1}$ acting on the genetic variance $G_{1,1}$, and the indirect response is $G_{1,2}\beta_{sel,2} + \cdots + G_{1,12}\beta_{sel,12}$ due to selection gradients on other traits acting on the genetic covariances. Therefore, the indirect response of trait $j$ is caused by genetic correlations between trait $j$ and other traits that are under selection. This explanation here can be extended to other traits, as well as the direct and indirect selection differentials.

**Multivariate selection parameters in simulated example**

We simulated a single generation of selection to investigate if it is possible to obtain correct selection parameters from the multivariate breeder's equation. All simulation was done using AlphaSimR (Gaynor et al. 2021). The simulation used the same fictitious species with 10 chromosomes as described previously. We created 10 unrelated founders and made 300 random crosses and retained 1 $F_1$ individual from each cross. We then created one doubled-haploid (DH) individual from each of the $F_1$ individual. These 300 DH individuals became the pre-selected population. We simulated three traits and 15 QTLs for each trait, where there are 9 unique QTLs for each trait, two pleiotropic QTLs that are shared by any two traits, and 2 pleiotropic QTLs that are shared by all three traits. This ensured that the traits have some degree of correlation. We set the QTL effects ($\mathrm{a}$) based on the founder allele frequencies: one $4\mathrm{a}$-QTL at 0.1 frequency, two $3\mathrm{a}$-QTL at 0.2 frequency, four $2\mathrm{a}$-QTL at 0.3 frequency, and eight $\mathrm{a}$-QTL at 0.4 to 0.5 frequency. We scaled all QTL effects such that the additive genetic variance is 1, and we added residual effect with a variance of 1 so that all traits have a heritability of 0.5.

Next, we applied three different selection schemes: select for individuals with top 10% of trait 1 (Sel1), select for individuals with either top 10% of trait 1 or 2 (Sel2), and select for individuals with top 10% of trait 1, 2 or 3 (Sel3). The differences between mean of selected individuals and mean of pre-selected population were calculated as the true selection differentials ($\mathrm{S}$). Similar to previous crosses with the founders, we made 300 random crosses among the selected individuals and created a DH individual from each of the resulting $F_1$ individual. These 300 individuals became the post-selected population. The differences between mean of post- and pre-selected populations were calculated as the realized selection responses ($\Delta\mathrm{Z}$). We calculated the true genetic ($\mathrm{G}$) and phenotypic ($\mathrm{P}$) variance-covariance matrices directly from the genetic values in the pre-selected population, and we also estimated $\mathrm{G}$ and $\mathrm{P}$ from the pre-selected population using the mixed linear model approach as shown in Equation 10 and 11.

For Sel1, Sel2 and Sel3, we determined the true, realized and estimated multivariate selection parameters. We obtained the true $\beta_{sel}$, $Z$ and $i$ from solving $S = P\beta_{sel}$, $\Delta Z = G\beta_{sel}$ and $i = S/\sqrt{diag(P)}$ with true $S$, $G$ and $P$. We calculated the realized $\beta_{sel}$, $S$ and $i$ from solving $\Delta Z = G\beta_{sel}$, $S = P\beta_{sel}$, and $i =$

$S/\sqrt{diag(P)}$ with realized $\Delta Z$ and true $G$ and $P$. Lastly, we computed the realized $\beta_{sel}$, $S$ and $i$ from solving $\Delta Z = G\beta_{sel}$, $S = P\beta_{sel}$, and $i = S/\sqrt{diag(P)}$ with realized $\Delta Z$ and estimated $G$ and $P$. We evaluated the accuracies of multivariate selection parameters by correlating the true values against either realized or estimated values.

### Supplementary Results

#### Multivariate selection parameters in the TG panel

We calculated the genetic variance-covariance ($\mathrm{G}$) matrix for all 12 traits using the TG panel GBS marker data (Table S11) as a pre-requisite to the multivariate selection analysis. To verify that $\mathrm{G}$ is representative across years of release and countries of origin, we compared the overall $\mathrm{G}$ with $\mathrm{G}$'s calculated from subsets of the TG panel for three year-groups of roughly equal number of varieties ($\mathrm{G}_{y1}$: 1948 to 1990, $\mathrm{G}_{y2}$: 1991 to 2001, $\mathrm{G}_{y3}$: 2002 to 2007) and three countries of origin ($\mathrm{G}_{DE}$: Germany, $\mathrm{G}_{FR}$: France, $\mathrm{G}_{UK}$: United Kingdom). The comparisons suggested that the $\mathrm{G}$'s are similar when tested using Random Skewers (Cheverud and Marroig 2007) but less so when tested using Mantel test (Mantel 1967) (Table S10). Because the $\mathrm{G}$'s may be influenced by the trait magnitudes, we also compared the genetic correlation ($\mathrm{g}$) matrix similarly. In contrast to the previous comparisons with $\mathrm{G}$'s, the results for $\mathrm{g}$'s indicated that the matrices were more dissimilar when tested using Randoms Skewers but more similar when tested using Mantel test (Table S10). However, we found higher similarities between $\mathrm{G}$ and $\mathrm{G}_{y1,y2,y3}$ or between $\mathrm{G}$ and $\mathrm{G}_{DE,FR,UK}$ than among $\mathrm{G}_{y1,y2,y3}$ or $\mathrm{G}_{DE,FR,UK}$. This result indicates that the overall $\mathrm{G}$ is a reasonable average estimate of the variations that exist due to years of release and countries of origin.

Using the selection response ($\Delta\mathrm{Z}$) estimated from linear regression of each trait over years of release and $\mathrm{G}$, we computed the selection gradient ($\beta_{sel}$) from the multivariate breeder's equation of $\Delta Z = G\beta_{sel}$ (Table S8). Between $\Delta\mathrm{Z}$ and $\beta_{sel}$, the directions of the vectors' elements are only the same for 7 out of 12 traits. This suggests that the selection response is divergent from the selection gradient, which is the relationship between trait value and individual fitness (Lande and Arnold 1983). The divergence is almost perpendicular at angle of 85.8° between the two vectors, therefore implies that a strong selection gradient is required to achieve the desired selection response. In addition, we converted $\beta_{sel}$ into an easier-to-understand measure called the selection differential ($\mathrm{S}$) (Table S8) that is computed from $S = P\beta_{sel}$ where $\mathrm{P}$ is the phenotypic variance-covariance matrix. Similar to its univariate counterpart, $\mathrm{S}$ is the difference between the mean of selected subset from the mean of population. Like our previous results on $\Delta\mathrm{Z}$ and $\beta_{sel}$, the directions of $\Delta\mathrm{Z}$ and $\mathrm{S}$ elements are the same for only 5 out of 12 traits with an angle of 79.1° between the two vectors. We also computed the selection intensity ($\mathrm{i}$) which is a standardized measure of $\mathrm{S}$ (Table S8). Overall, the selection force, as measured by $\beta_{sel}$, $\mathrm{S}$ and $\mathrm{i}$, is not well aligned with the selection response, $\Delta\mathrm{Z}$.

Subsequently, we showed the partitioning of $\Delta\mathrm{Z}$, $\mathrm{S}$ and $\mathrm{i}$ into the direct and indirect components by isolating the products of $\beta_{sel}$ with the matrix diagonals (direct) and matrix off-diagonals (Table S8). This method allows us to quantify the amount of each selection measure that is directly due to the available variation within a trait or indirectly due to the covariation with other traits. Using YLD as an example, we observe a large proportion of direct effect in $\Delta\mathrm{Z}$ but a slightly higher proportion of indirect over direct effect in $\mathrm{S}$ and $\mathrm{i}$. This result suggests that the selection force on YLD receives a higher contribution from selection on other traits more than selection on the trait itself. However, the selection response in YLD receives little to no selection response that can be attributed due to covariation with other traits. In another example with HT, we find positive direct effects in $\Delta\mathrm{Z}$, $\mathrm{S}$ and $\mathrm{i}$, which contradicts the known selection on dwarfing genes like *Rht1*, *Rht2* and *Rht24* (Pearce et al. 2011, Würschum et al.

2017). Given the uncertainties in the multivariate selection parameters, it is best to treat these estimates with caution as we explore the cause of issues later using simulation.

To test the consistency in multivariate selection parameters across genotyping platforms, we applied the same approach to the TG panel that was genotyped using the Diversity Array Technology (DArT) (Bentley et al. 2014). First, the comparison of $\mathrm{G}$'s and $\mathrm{g}$'s between GBS and DArT showed highly significant similarity in both Random Skewers (Cheverud and Marroig 2007) and Mantel tests (Mantel 1976) (Table S10). For consistency, we matched the phenotypic data from Bentley et al. (2014) to the DArT marker data and phenotypic data from Ladejobi et al. (2019) to the GBS marker data. The resulting $\Delta\mathrm{Z}$ are the same between the two except for YLD, which has only a small difference. Given that the $\mathrm{G}$ and $\Delta\mathrm{Z}$ are highly similar, we expected the $\beta_{\mathrm{sel}}$ to be similar. In contrast, the $\beta_{\mathrm{sel}}$ shared only 4 traits in the same directions and are separated by an angle of 104.6°. As a result, the estimates of $\mathrm{S}$ and $\mathrm{i}$, as well as direct and indirect effects, are vastly different (Table S8).

### Multivariate selection parameters in a simulated example

To better understand the issues with multivariate selection parameters, we used a simple simulation with a single generation of selection involving three genetically correlated traits. The details of the simulation can be found in the Materials and Methods section. First, we compared $\mathrm{G}$ and $\mathrm{E}$ between the version estimated from mixed model and the version derived from true simulated values. $\mathrm{E}$ is the residual variance-covariance matrix where $\mathrm{P} = \mathrm{G} + \mathrm{E}$. While the means of estimated variances were close to true simulated variance of 1 (Figure S10), the precisions of the estimates were poor (Table S9). Next, we simulated three different types of selection: (1) Sel1: select on individuals with top 10% in trait 1, (2) Sel2: select on individuals with top 10% in trait 1 or 2, (3) Sel3: select on individuals with top 10% in trait 1, 2 or 3.

For each type of selection, we compared the estimated or realized values of $\Delta\mathrm{Z}$, $\beta_{\mathrm{sel}}$, $\mathrm{S}$ and $\mathrm{i}$ to the true or expected values. First, we calculated the true values of $\Delta\mathrm{Z}$, $\beta_{\mathrm{sel}}$ and $\mathrm{i}$ using the true values for $\mathrm{S}$, $\mathrm{G}$ and $\mathrm{P}$. Next, we obtained the realized values of $\beta_{\mathrm{sel}}$, $\mathrm{S}$ and $\mathrm{i}$ using the realized $\Delta\mathrm{Z}$ and true $\mathrm{G}$ and $\mathrm{P}$. We also obtained the estimated values of $\beta_{\mathrm{sel}}$, $\mathrm{S}$ and $\mathrm{i}$ using the realized $\Delta\mathrm{Z}$ and estimated $\mathrm{G}$ and $\mathrm{P}$. Across Sel1, Sel2 and Sel3, we found that the correlations were higher between realized and true values than between estimated and true values (Table S9, Figure S11-S16). This result further highlights the issues due to poor estimates of $\mathrm{G}$ and $\mathrm{P}$. However, the correlations between the realized and expected values are far from perfect, which suggest that the discrepancies between observed and expected $\Delta\mathrm{Z}$ are sufficient to distort the realized values of $\beta_{\mathrm{sel}}$, $\mathrm{S}$ and $\mathrm{i}$. Lastly, we observed that the correlations were generally higher in Sel1, followed by Sel2 and Sel3 (Table S9). This may suggest that the multivariate selection parameters get poorer as more traits are directly selected.

### Supplementary Tables

#### Table S1. Parametric control estimates and percent significant markers.

Maximum likelihood estimates (MLE) of the mean (δ) and standard deviation (σ) for model correction using Parametric Control (PC) at various allele frequency change thresholds (t = 0.05 to 0.50). The values are shown as the mean and range (in parentheses) across n-simulations with converged MLE (conv.). The first row with t = 0.00 is from the uncorrected model. S: selected, U: unselected population.

| | δ | | σ | | Significant markers (%) | | |
|---|---|---|---|---|---|---|---|
| t | S | U | S | U | S | U | conv. |
| 0.00 | NA | NA | NA | NA | 1.942 (0.858, 3.535) | 0.089 (0, 0.391) | NA |
| 0.05 | 0.0002 (-0.0143, 0.0162) | 0.0006 (-0.0157, 0.0143) | 0.5994 (0.5470, 0.6529) | 0.5814 (0.5498, 0.6079) | 11.152 (8.013, 14.393) | 6.391 (4.042, 9.988) | 100 |
| 0.06 | -0.0004 (-0.0192, 0.0178) | 0.0007 (-0.0199, 0.0194) | 0.6875 (0.6502, 0.7467) | 0.6656 (0.6292, 0.7020) | 7.284 (4.418, 9.943) | 3.212 (1.795, 5.012) | 100 |
| 0.07 | -0.0001 (-0.0219, 0.0220) | 0.0012 (-0.0209, 0.0204) | 0.7633 (0.7182, 0.8116) | 0.7398 (0.6955, 0.7812) | 5.093 (2.849, 7.578) | 1.613 (0.707, 2.706) | 100 |
| 0.08 | 0 (-0.0205, 0.0229) | 0.0007 (-0.0210, 0.0188) | 0.8341 (0.7941, 0.8825) | 0.8086 (0.7710, 0.8498) | 3.723 (1.945, 5.833) | 0.819 (0.281, 1.385) | 100 |
| 0.09 | -0.0008 (-0.0188, 0.0231) | 0.0003 (-0.0215, 0.0196) | 0.8990 (0.8556, 0.9518) | 0.8710 (0.8319, 0.9096) | 2.857 (1.476, 4.591) | 0.414 (0.060, 1.039) | 98 |
| 0.10 | -0.0007 (-0.0248, 0.0220) | 0.0000 (-0.0202, 0.0223) | 0.9578 (0.9105, 1.0068) | 0.9270 (0.8917, 0.9713) | 2.297 (1.117, 3.798) | 0.218 (0.011, 0.557) | 96 |
| 0.11 | 0.0002 (-0.0229, 0.0217) | 0.0001 (-0.0217, 0.0222) | 1.0128 (0.9622, 1.0589) | 0.9791 (0.9411, 1.0207) | 1.867 (0.961, 3.097) | 0.109 (0, 0.440) | 92 |
| 0.12 | -0.0003 (-0.0218, 0.0196) | -0.0003 (-0.0224, 0.0205) | 1.0645 (1.0096, 1.1169) | 1.0270 (0.9899, 1.0776) | 1.554 (0.750, 2.692) | 0.058 (0, 0.272) | 92 |
| 0.13 | 0.0001 (-0.0238, 0.0186) | 0.0001 (-0.0189, 0.0207) | 1.1127 (1.0504, 1.1702) | 1.0724 (1.0306, 1.1236) | 1.313 (0.618, 2.506) | 0.029 (0, 0.239) | 93 |
| 0.14 | -0.0002 (-0.0200, 0.0217) | 0.0007 (-0.0231, 0.0213) | 1.1578 (1.0935, 1.2130) | 1.1140 (1.0709, 1.1787) | 1.142 (0.476, 2.177) | 0.017 (0, 0.147) | 94 |
| 0.15 | 0.0002 (-0.0212, 0.0213) | 0.0007 (-0.0223, 0.0228) | 1.2002 (1.1327, 1.2609) | 1.1519 (1.1042, 1.2186) | 0.994 (0.391, 2.007) | 0.012 (0, 0.092) | 93 |
| 0.16 | 0.0003 (-0.0237, 0.0224) | 0.0007 (-0.0212, 0.0208) | 1.2393 (1.1718, 1.3012) | 1.1888 (1.1337, 1.2567) | 0.882 (0.332, 1.848) | 0.007 (0, 0.065) | 95 |
| 0.17 | 0.0001 (-0.0248, 0.0218) | 0.0011 (-0.0196, 0.0221) | 1.2780 (1.2161, 1.3454) | 1.2228 (1.1612, 1.2916) | 0.791 (0.230, 1.607) | 0.003 (0, 0.054) | 95 |
| 0.18 | 0.0002 (-0.0233, 0.0198) | 0.0007 (-0.0231, 0.0233) | 1.3133 (1.2562, 1.3828) | 1.2536 (1.1898, 1.3298) | 0.709 (0.197, 1.431) | 0.002 (0, 0.033) | 96 |
| 0.19 | 0.0002 (-0.0216, 0.0200) | 0.0012 (-0.0200, 0.0237) | 1.3457 (1.2894, 1.4203) | 1.2818 (1.2121, 1.3634) | 0.634 (0.170, 1.146) | 0.001 (0, 0.027) | 96 |
| 0.20 | -0.0004 (-0.0231, 0.0204) | 0.0008 (-0.0185, 0.0256) | 1.3758 (1.3185, 1.4528) | 1.3074 (1.2363, 1.3972) | 0.569 (0.142, 1.066) | 0 (0, 0.016) | 96 |
| 0.21 | -0.0008 (-0.0245, 0.0223) | 0.0007 (-0.0215, 0.0252) | 1.4049 (1.3440, 1.4807) | 1.3303 (1.2555, 1.4247) | 0.521 (0.137, 1.050) | 0 (0, 0.016) | 97 |
| 0.22 | -0.0010 (-0.0261, 0.0215) | 0.0009 (-0.0226, 0.0263) | 1.4325 (1.3667, 1.5127) | 1.3524 (1.2679, 1.4496) | 0.470 (0.109, 0.957) | 0 (0, 0.005) | 97 |
| 0.23 | -0.0007 (-0.0250, 0.0257) | 0.0011 (-0.0258, 0.0263) | 1.4576 (1.3884, 1.5392) | 1.3712 (1.2799, 1.4770) | 0.430 (0.092, 0.886) | 0 (0, 0.005) | 98 |
| 0.24 | -0.0007 (-0.0252, 0.0251) | 0.0009 (-0.0275, 0.0269) | 1.4817 (1.4103, 1.5636) | 1.3882 (1.2889, 1.4984) | 0.392 (0.066, 0.760) | 0 (0, 0) | 98 |

Table S1 continues to the next page.

Table S1 continues from the previous page.

| t | δ | | σ | | Significant markers (%) | | conv. |
|---|---|---|---|---|---|---|---|
| | S | U | S | U | S | U | |
| 0.25 | -0.0007<br>(-0.0260, 0.0268) | 0.0008<br>(-0.0269, 0.0256) | 1.5045<br>(1.4262, 1.5918) | 1.4028<br>(1.2990, 1.5136) | 0.357<br>(0.044, 0.735) | 0<br>(0, 0) | 98 |
| 0.26 | -0.0013<br>(-0.0270, 0.0257) | 0.0007<br>(-0.0303, 0.0241) | 1.5251<br>(1.4464, 1.6161) | 1.4162<br>(1.3064, 1.5332) | 0.330<br>(0.027, 0.697) | 0<br>(0, 0) | 97 |
| 0.27 | -0.0013<br>(-0.0288, 0.0243) | 0.0005<br>(-0.0293, 0.0242) | 1.5450<br>(1.4696, 1.6380) | 1.4280<br>(1.3107, 1.5512) | 0.303<br>(0.005, 0.659) | 0<br>(0, 0) | 98 |
| 0.28 | -0.0011<br>(-0.0294, 0.0269) | 0.0004<br>(-0.0296, 0.0252) | 1.5625<br>(1.4897, 1.6603) | 1.4386<br>(1.3163, 1.5714) | 0.284<br>(0.005, 0.648) | 0<br>(0, 0) | 98 |
| 0.29 | -0.0014<br>(-0.0295, 0.0253) | 0.0004<br>(-0.0290, 0.0228) | 1.5790<br>(1.5032, 1.6778) | 1.4477<br>(1.3199, 1.5832) | 0.265<br>(0.005, 0.611) | 0<br>(0, 0) | 98 |
| 0.30 | -0.0013<br>(-0.0293, 0.0248) | 0.0003<br>(-0.0287, 0.0239) | 1.5943<br>(1.5109, 1.6939) | 1.4549<br>(1.3229, 1.5957) | 0.251<br>(0.005, 0.606) | 0<br>(0, 0) | 98 |
| 0.31 | -0.0012<br>(-0.0310, 0.0267) | 0.0004<br>(-0.0267, 0.0237) | 1.6088<br>(1.5185, 1.7096) | 1.4616<br>(1.3243, 1.6066) | 0.235<br>(0.005, 0.578) | 0<br>(0, 0) | 98 |
| 0.32 | -0.0013<br>(-0.0308, 0.0237) | 0.0001<br>(-0.0271, 0.0239) | 1.6224<br>(1.5267, 1.7176) | 1.4672<br>(1.3295, 1.6194) | 0.221<br>(0, 0.567) | 0<br>(0, 0) | 98 |
| 0.33 | -0.0008<br>(-0.0325, 0.0333) | 0.0001<br>(-0.0282, 0.0236) | 1.6344<br>(1.5421, 1.7337) | 1.4720<br>(1.3303, 1.6227) | 0.212<br>(0, 0.540) | 0<br>(0, 0) | 99 |
| 0.34 | -0.0008<br>(-0.0324, 0.0320) | 0.0001<br>(-0.0279, 0.0253) | 1.6458<br>(1.5479, 1.7446) | 1.4756<br>(1.3327, 1.6260) | 0.199<br>(0, 0.535) | 0<br>(0, 0) | 99 |
| 0.35 | -0.0010<br>(-0.0322, 0.0303) | 0.0001<br>(-0.0279, 0.0249) | 1.6567<br>(1.5580, 1.7542) | 1.4787<br>(1.3327, 1.6308) | 0.191<br>(0, 0.524) | 0<br>(0, 0) | 99 |
| 0.36 | -0.0011<br>(-0.0348, 0.0305) | 0.0001<br>(-0.0286, 0.0252) | 1.6671<br>(1.5693, 1.7683) | 1.4810<br>(1.3327, 1.6329) | 0.182<br>(0, 0.518) | 0<br>(0, 0) | 99 |
| 0.37 | -0.0010<br>(-0.0336, 0.0297) | 0.0002<br>(-0.0286, 0.0252) | 1.6768<br>(1.5755, 1.7795) | 1.4831<br>(1.3327, 1.6366) | 0.174<br>(0, 0.513) | 0<br>(0, 0) | 99 |
| 0.38 | -0.0010<br>(-0.0338, 0.0296) | 0.0002<br>(-0.0291, 0.0257) | 1.6857<br>(1.5807, 1.7894) | 1.4848<br>(1.3342, 1.6374) | 0.166<br>(0, 0.502) | 0<br>(0, 0) | 99 |
| 0.39 | -0.0011<br>(-0.0356, 0.0298) | 0.0002<br>(-0.0286, 0.0257) | 1.6940<br>(1.5849, 1.8016) | 1.4860<br>(1.3342, 1.6383) | 0.159<br>(0, 0.497) | 0<br>(0, 0) | 99 |
| 0.40 | -0.0012<br>(-0.0402, 0.0279) | 0.0001<br>(-0.0286, 0.0259) | 1.7024<br>(1.5867, 1.8220) | 1.4872<br>(1.3342, 1.6402) | 0.151<br>(0, 0.497) | 0<br>(0, 0) | 99 |
| 0.41 | -0.0012<br>(-0.0409, 0.0275) | 0.0000<br>(-0.0286, 0.0259) | 1.7098<br>(1.5885, 1.8439) | 1.4879<br>(1.3342, 1.6409) | 0.146<br>(0, 0.480) | 0<br>(0, 0) | 99 |
| 0.42 | -0.0010<br>(-0.0392, 0.0265) | 0.0001<br>(-0.0286, 0.0259) | 1.7174<br>(1.5984, 1.8578) | 1.4887<br>(1.3342, 1.6420) | 0.14<br>(0, 0.475) | 0<br>(0, 0) | 99 |
| 0.43 | -0.0009<br>(-0.0386, 0.0270) | 0.0001<br>(-0.0286, 0.0259) | 1.7242<br>(1.6004, 1.8697) | 1.4891<br>(1.3342, 1.6420) | 0.135<br>(0, 0.469) | 0<br>(0, 0) | 99 |
| 0.44 | -0.0010<br>(-0.0371, 0.0260) | 0.0000<br>(-0.0286, 0.0259) | 1.7304<br>(1.6026, 1.8788) | 1.4894<br>(1.3342, 1.6425) | 0.131<br>(0, 0.453) | 0<br>(0, 0) | 99 |
| 0.45 | -0.0010<br>(-0.0371, 0.0292) | 0.0000<br>(-0.0286, 0.0259) | 1.7363<br>(1.6113, 1.8939) | 1.4896<br>(1.3342, 1.6425) | 0.127<br>(0, 0.447) | 0<br>(0, 0) | 99 |
| 0.46 | -0.0009<br>(-0.0358, 0.0301) | 0.0000<br>(-0.0286, 0.0259) | 1.7420<br>(1.6128, 1.9033) | 1.4898<br>(1.3342, 1.6425) | 0.123<br>(0, 0.442) | 0<br>(0, 0) | 99 |
| 0.47 | -0.0008<br>(-0.0374, 0.0294) | 0.0000<br>(-0.0286, 0.0259) | 1.7480<br>(1.6199, 1.9128) | 1.4900<br>(1.3342, 1.6425) | 0.120<br>(0, 0.440) | 0<br>(0, 0) | 99 |
| 0.48 | -0.0009<br>(-0.0384, 0.0288) | 0.0001<br>(-0.0286, 0.0259) | 1.7535<br>(1.6219, 1.9196) | 1.4902<br>(1.3342, 1.6425) | 0.117<br>(0, 0.440) | 0<br>(0, 0) | 99 |
| 0.49 | -0.0011<br>(-0.0386, 0.0304) | 0.0001<br>(-0.0286, 0.0259) | 1.7590<br>(1.6279, 1.9325) | 1.4903<br>(1.3342, 1.6425) | 0.114<br>(0, 0.440) | 0<br>(0, 0) | 99 |
| 0.50 | -0.0011<br>(-0.0382, 0.0307) | 0.0000<br>(-0.0286, 0.0259) | 1.7647<br>(1.6309, 1.9376) | 1.4904<br>(1.3342, 1.6425) | 0.110<br>(0, 0.440) | 0<br>(0, 0) | 99 |

**Table S2. Genomic positions of 50 GWAS QTLs in the TG panel.**

The IWGSC RefSeq v1.0 physical positions of the peaks and boundaries (Ladejobi et al. 2019) are clustered into groups according to their positions. The traits associated with each group are listed along with the percent variance explained (PVE). The overlapping QSLs from RALLY in TG and WAGTAIL panels are also shown.

| QTL | Chr | Position (bp) | | | Trait (PVE) | Overlapping QSL/QTL | |
|---|---|---|---|---|---|---|---|
| | | Start | End | Peak | | TG | WAGTAIL |
| 1 | 1A | - | - | 42,151,448 | PROT (2.9) | 1 | 0 |
| 2 | 1A | 532,433,018 | 532,529,494 | 532,529,494 | AWNS (4.3) | 0 | 0 |
| 3 | 1B | - | - | 4,341,220 | FT (4.1) | 0 | 0 |
| 4 | 1B | 40,429,328 | 40,806,011 | 40,806,011 | FT (3.7) | 0 | 0 |
| 5 | 1B | 628,112,579 | 636,796,017 | 628,112,579 | AWNS (9.6) | 0 | 0 |
| 6 | 1D | - | - | 343,103,677 | AWNS (3.2) | 0 | 0 |
| 7 | 2A | - | - | 3,634,548 | LODG (3.9) | 4 | 5 |
| 8 | 2A | - | - | 748,997,740 | YLD (4.0) | 0 | 0 |
| 9 | 2B | - | - | 1,467,644 | LODG (4.8) | 0 | 0 |
| 10 | 2B | 688,433,192 | 688,433,222 | 688,433,192 | FT (4.0) | 6 | 9 |
| 11 | 2B | 790,828,845 | 800,780,364 | 794,977,501 | AWNS (3.7) | 0 | 0 |
| 12 | 2D | - | - | 7,738,369 | AWNS (2.9) | 0 | 0 |
| 13 | 2D | - | - | 8,787,004 | LODG (5.0) | 0 | 0 |
| 14 | 2D | - | - | 31,468,893 | FT (9.6), HT (3.5), MAT (6.1) | 0 | 0 |
| 15 | 2D | - | - | 42,097,013 | FT (9.3), HT (3.9), MAT (7.2) | 0 | 0 |
| 16 | 2D | 572,734,652 | 608,800,104 | 572,918,169 | LODG (5.0) | 0 | 0 |
| 17 | 3A | - | - | 161,444,257 | AWNS (3.2) | 0 | 0 |
| 18 | 3A | - | - | 463,564,378 | LODG (3.5) | 0 | 0 |
| 19 | 3A | - | - | 495,524,635 | HT (2.9) | 7 | 13 |
| 20 | 3B | - | - | 5,601,689 | PROT (2.7) | 0 | 0 |
| 21 | 4A | 222,595,975 | 298,655,753 | 298,655,753 | LODG (3.7) | 0 | 14 |
| 22 | 4A | 639,058,378 | 729,741,651 | 639,058,378 | PROT (3.1) | 10 | 0 |
| 22 | 4A | 719,113,347 | 729,872,889 | 719,113,347 | AWNS (3.5) | 0 | 0 |
| 23 | 4A | 743,082,595 | 744,433,737 | 743,226,828 | HT (3.4) | 0 | 0 |
| 24 | 4B | 21,378,087 | 21,379,808 | 21,378,087 | HT (4.2) | 0 | 0 |
| 25 | 4B | - | - | 55,242,733 | WK (3.7) | 0 | 0 |
| 26 | 4B | - | - | 520,086,297 | LODG (4.1) | 11 | 0 |
| 27 | 4B | - | - | 591,282,163 | AWNS (3.3) | 11 | 0 |
| 28 | 5A | 59,536,488 | 67,301,668 | 63,500,759 | WK (4.6) | 12 | 16 |
| 29 | 5A | 665,971,659 | 685,792,099 | 679,803,244 | HT (3.1) | 0 | 0 |
| 29 | 5A | 680,532,103 | 709,755,448 | 708,820,599 | FT (3.9), AWNS (28.9) | 0 | 0 |
| 30 | 5B | - | - | 107,367,824 | PROT (3.8) | 0 | 0 |
| 31 | 5B | 580,426,360 | 588,757,384 | 588,757,384 | AWNS (5.3) | 0 | 0 |
| 32 | 6A | 3,758,004 | 21,719,436 | 16,462,141 | AWNS (10.8) | 15 | 0 |
| 33 | 6A | 85,580,220 | 91,681,786 | 88,260,960 | YLD (3.9) | 16 | 18 |
| 34 | 6A | 101,430,018 | 101,851,274 | 101,430,018 | HT (4.5), LODG (4.4) | 16 | 18 |
| 34 | 6A | 85,263,886 | 115,457,176 | 112,688,966 | PROT (5.5) | 16 | 18 |
| 35 | 6A | - | - | 151,492,080 | PROT (2.8) | 16 | 18 |
| 36 | 6A | 373,461,190 | 452,372,111 | 422,756,419 | HT (5.0) | 16 | 18 |
| 36 | 6A | 416,675,591 | 448,706,137 | 448,010,707 | PROT (3.8) | 16 | 18 |

| | | | | | | | |
|---|---|---|---|---|---|---|---|
| 36 | 6A | 373,461,190 | 450,106,742 | 450,106,742 | LODG (6.1) | 16 | 18 |
| 37 | 6A | 453,334,056 | 461,434,500 | 454,675,620 | HT (4.5) | 16 | 18 |
| 37 | 6A | 454,675,620 | 465,641,292 | 465,641,261 | LODG (4.7) | 16 | 18 |
| 38 | 6A | 614,089,848 | 615,386,089 | 614,663,042 | FT (3.6) | 17 | 0 |
| 39 | 6B | - | - | 106,381,266 | PROT (3.9) | 0 | 0 |
| 40 | 6D | - | - | 2,449,348 | LODG (3.7) | 0 | 0 |
| 41 | 6D | - | - | 5,155,743 | AWNS (6.3) | 0 | 0 |
| 42 | 6D | 461,923,517 | 462,303,873 | 461,923,517 | FT (4.5) | 0 | 0 |
| 43 | 7A | 14,418,771 | 15,750,691 | 15,031,731 | HT (3.5) | 0 | 19 |
| 43 | 7A | 14,418,771 | 22,872,792 | 15,639,304 | FT (6.6) | 0 | 19 |
| 44 | 7A | - | - | 62,185,902 | PROT (2.8) | 0 | 0 |
| 45 | 7A | - | - | 278,811,242 | LODG (3.4) | 0 | 0 |
| 46 | 7A | - | - | 695,324,003 | YLD (3.4) | 0 | 0 |
| 47 | 7B | - | - | 11,703,887 | HT (3.0) | 0 | 0 |
| 48 | 7B | 45,162,103 | 65,376,987 | 45,162,103 | LODG (4.3) | 21 | 0 |
| 48 | 7B | 50,826,708 | 63,170,244 | 63,134,685 | YLD (3.7), PROT (3.7) | 21 | 0 |
| 49 | 7B | 678,635,628 | 702,820,571 | 678,635,628 | LODG (3.7) | 22 | 0 |
| 49 | 7B | - | - | 702,820,571 | HT (2.5) | 22 | 0 |
| 50 | 7D | - | - | 16,704,573 | FT (4.9) | 0 | 0 |

**Table S3. Co-localization between RALLY QSLs and others.**

QTLs identified from three recent studies in wheat are compared against the RALLY QSLs. All physical positions are listed according to the IWGSC RefSeq v1.0, except for the meta QTLs by Yang et al. (2021) which have not been explicitly stated.

| QSL | Position (chromosome + interval in Mb) | QTL co-localization (chromosome + interval in Mb) | | |
|---|---|---|---|---|
| | | MAGIC diverse GWAS (Scott et al. 2021) | Meta QTL (Yang et al. 2021) | Introgression (Cheng et al. 2019) |
| 1 | 1A (36.8 - 42.2) | - | - | - |
| 2 | 1A (105.9 - 395.5) | 1A (389.0 - 468.4) | - | 1A (210.3 - 233.7) |
| 3 | 1B (52.8 - 573.0) | 1B (95.6 - 566.7) | 1B-4 (542.9 - 563.1) | 1B (560.0 - 560.1) |
| 4 | 2A (0.5 - 36.1) | Yr17 | 2A-2 (4.0 - 10.0), 2A-3 (28.0 - 30.5), 2A-4 (33.3 - 34.5) | 2A (0.0 - 11.0) |
| 5 | 2A (56.2 - 115.4) | - | 2A-1 (59.4 - 71.6) | - |
| 6 | 2B (10.9 - 785.6) | Ppd-B1, Yr7/Yr5/YrSP | 2B-3 (50.7 - 80.1), 2B-4 (104.1 - 730.2) | 2B (89.6 - 748.7) |
| 7 | 3A (488.3 - 574.6) | - | 3A-3 (471.4 - 533.6) | 3A (508.3 - 509.4) |
| 8 | 3B (19.1 - 31.4) | - | 3B-3 (12.3 - 26.8) | - |
| 9 | 3B (813.3 - 830.0) | - | - | 3B (822.4 - 829.7) |
| 10 | 4A (507.7 - 695.9) | 4A (599.1 - 610.1) | 4A-2 (606.7 - 611.1), 4A-3 (591.0 - 597.1), 4A-4 (618.3 - 621.7), 4A-5 (616.6 - 626.8), 4A-6 (654.5 - 668.8) | 4A (372.4 - 679.2) |
| 11 | 4B (507.2 - 593.8) | - | - | - |
| 12 | 5A (31.1 - 449.8) | 5A (13.7 - 34.2) | - | - |
| 13 | 5B (681.3 - 703.9) | - | 5B-6 (679.4 - 692.6) | - |
| 14 | 5D (43.4 - 233.7) | - | 5D-3 (44.6 - 54.7), 5D-5 (48.7 - 80.3) | 5D (188.3 - 188.8) |
| 15 | 6A (0.7 - 5.1) | - | 6A-1 (0.3 - 6.7) | - |
| 16 | 6A (61.8 - 545.4) | TaGW2, Rht24 | 6A-3 (58.2 - 85.8), 6A-4 (518.8 - 538.4) | 6A (281.5 - 282.0) |
| 17 | 6A (596.6 - 617.3) | 6A (609.3 - 618.3) | 6A-7 (595.0 - 602.9), 6A-8 (606.6 - 615.1) | 6A (596.2 - 598.6) |
| 18 | 7A (610.2 - 612.6) | - | 7A-5 (108.8 - 616.6) | - |
| 19 | 7A (669.8 - 695.0) | WAPO-A1 | 7A-6 (675.3 - 688.7), 7A-7 (694.4 - 703.6) | 7A (669.1 - 698.6) |
| 20 | 7B (3.4 - 4.8) | - | 7B-1 (1.0 - 6.2) | - |
| 21 | 7B (40.3 - 58.9) | - | - | - |
| 22 | 7B (698.2 - 707.9) | - | 7B-6 (697.5 - 707.7) | - |

**Table S4. Genomic positions of 19 RALLY QSLs in the WAGTAIL panel.**

The IWGSC RefSeq v1.0 physical positions of the peaks and LD boundaries are shown, along with the P-values associated with the peaks and overlapping QSLs/QTLs from RALLY in TG panel and GWAS in TG panel (Ladejobi et al. 2019).

| QSL | Chr | Position (bp) | | | $-\log_{10}P$ | Overlapping QSL/QTL | |
|---|---|---|---|---|---|---|---|
| | | Start | End | Peak | | TG | GWAS |
| 1 | 1A | 11,950,553 | 1,336,994 | 14,253,976 | 7.173 | 0 | 0 |
| 2 | 1A | 462,001,757 | 38,729,508 | 490,087,895 | 11.148 | 1,2 | 0 |
| 3 | 1B | 303,680,331 | 303,680,331 | 571,707,456 | 5.665 | 3 | 0 |
| 4 | 1D | 56,751,172 | 34,621,416 | 56,753,155 | 8.717 | 0 | 0 |
| 5 | 2A | 24,278,890 | 2,646,023 | 32,139,650 | 7.634 | 4 | 7 |
| 6 | 2A | 59,558,265 | 58,394,953 | 77,163,127 | 5.429 | 5 | 0 |
| 7 | 2A | 639,988,472 | 605,184,949 | 676,977,112 | 9.025 | 0 | 0 |
| 8 | 2A | 762,503,117 | 762,292,207 | 771,236,862 | 6.997 | 0 | 0 |
| 9 | 2B | 146,634,773 | 44,922,004 | 773,349,577 | 9.882 | 6 | 10 |
| 10 | 2B | 784,551,365 | 784,551,365 | 790,751,851 | 5.198 | 6 | 0 |
| 11 | 2D | 15,967,398 | 14,778,701 | 18,234,286 | 5.506 | 0 | 0 |
| 12 | 3A | 106,435,192 | 86,369,509 | 107,738,831 | 6.884 | 0 | 0 |
| 13 | 3A | 497,649,046 | 495,073,247 | 573,985,537 | 7.685 | 7 | 19 |
| 14 | 4A | 590,123,186 | 68,470,282 | 599,846,393 | 8.838 | 10 | 21 |
| 15 | 4B | 645,298,612 | 634,340,134 | 652,775,661 | 5.135 | 0 | 0 |
| 16 | 5A | 118,667,316 | 32,880,888 | 450,555,721 | 7.028 | 12 | 28 |
| 17 | 5B | 678,337,596 | 663,522,311 | 679,687,636 | 6.331 | 0 | 0 |
| 18 | 6A | 409,092,614 | 61,813,444 | 582,253,761 | 9.596 | 16 | 33,34,35,36,37 |
| 19 | 7A | 35,770,406 | 20,294,545 | 47,283,220 | 5.260 | 0 | 43 |
| 20 | 1A | 11,950,553 | 1,336,994 | 14,253,976 | 7.173 | 0 | 0 |
| 21 | 1A | 462,001,757 | 38,729,508 | 490,087,895 | 11.148 | 1,2 | 0 |
| 22 | 1B | 303,680,331 | 303,680,331 | 571,707,456 | 5.665 | 3 | 0 |

**Table S5. Likelihood ratio tests for local heritabilities**

Mixed linear models with and without the QSLs were compared using a likelihood ratio test (LRT) similar to the approach of Santantonio et al. (2019). The test statistic is $\chi^2_{df=1} = 2 \cdot (LL_1 - LL_0)$ where $LL_1$ is the log-likelihood of the model with QSL and $LL_0$ is the log-likelihood of the model without QSL. P-values are shown here.

| QSL | FT | LODG | YLD | HT | PROT | WK | AWNS | SPWT | TGW | EM2 | TILL | MAT |
|---|---|---|---|---|---|---|---|---|---|---|---|---|
| 1 | 1.000 | 0.998 | 0.936 | 0.087 | 0.159 | 0.171 | 0.999 | 1.000 | 1.000 | 0.324 | 0.999 | 1.000 |
| 2 | 1.000 | 0.450 | 0.140 | 0.047 | 0.552 | 0.999 | 1.000 | 0.824 | 0.235 | 0.272 | 1.000 | 1.000 |
| 3 | 1.000 | 0.994 | 0.615 | 0.993 | 0.482 | 0.998 | 0.867 | 0.868 | 1.000 | 0.741 | 0.763 | 0.462 |
| 4 | 1.000 | 0.995 | 0.997 | 0.994 | 0.998 | 0.998 | 0.570 | 0.999 | 1.000 | 1.000 | 0.031 | 1.000 |
| 5 | 1.000 | 0.142 | 1.000 | 0.298 | 0.998 | 1.000 | 0.995 | 0.886 | 1.000 | 1.000 | 0.999 | 1.000 |
| 6 | 1.000 | 0.976 | 0.181 | 1.000 | 1.000 | 0.546 | 0.999 | 0.998 | 0.367 | 1.000 | 0.686 | 0.063 |
| 7 | 1.000 | 0.994 | 0.932 | 0.992 | 0.673 | 1.000 | 1.000 | 0.999 | 0.408 | 0.996 | 0.192 | 1.000 |
| 8 | 0.566 | 0.994 | 1.000 | 0.570 | 0.998 | 0.998 | 0.996 | 0.193 | 0.997 | 0.993 | 1.000 | 1.000 |
| 9 | 1.000 | 0.855 | 0.763 | 0.996 | 0.360 | 0.998 | 1.000 | 1.000 | 0.996 | 0.994 | 0.576 | 1.000 |
| 10 | 0.519 | 0.448 | 1.000 | 0.992 | 0.998 | 0.663 | 0.586 | 0.276 | 0.114 | 0.728 | 0.170 | 0.063 |
| 11 | 1.000 | 0.062 | 0.049 | 0.892 | 0.762 | 0.522 | 0.390 | 0.255 | 0.241 | 0.994 | 0.999 | 1.000 |
| 12 | 1.000 | 0.779 | 0.497 | 0.992 | 0.488 | 0.531 | 0.183 | 0.745 | 0.996 | 0.607 | 1.000 | 0.893 |
| 13 | 1.000 | 0.994 | 0.756 | 0.840 | 1.000 | 0.208 | 0.997 | 1.000 | 0.306 | 0.993 | 1.000 | 1.000 |
| 14 | 0.295 | 0.996 | 0.152 | 0.992 | 0.038 | 0.524 | 1.000 | 0.933 | 0.995 | 0.996 | 0.999 | 0.380 |
| 15 | 0.469 | 0.996 | 1.000 | 0.992 | 1.000 | 0.406 | 0.368 | 0.909 | 0.332 | 0.418 | 1.000 | 1.000 |
| 16 | 1.000 | 0.097 | 2.97E-03 | 1.64E-06 | 1.73E-04 | 0.999 | 0.997 | 0.085 | 0.184 | 0.361 | 0.999 | 0.589 |
| 17 | 0.064 | 0.994 | 0.515 | 0.992 | 0.744 | 0.047 | 1.000 | 0.999 | 0.820 | 0.993 | 0.279 | 0.218 |
| 18 | 1.000 | 0.994 | 0.333 | 0.997 | 0.999 | 1.000 | 0.995 | 1.000 | 0.831 | 0.996 | 0.853 | 1.000 |
| 19 | 1.000 | 0.374 | 0.755 | 0.993 | 1.000 | 0.998 | 0.270 | 0.999 | 0.016 | 0.704 | 1.000 | 1.000 |
| 20 | 1.000 | 0.994 | 0.225 | 0.101 | 0.434 | 0.786 | 1.000 | 1.000 | 0.994 | 1.000 | 1.000 | 1.000 |
| 21 | 1.000 | 8.45E-03 | 1.000 | 0.397 | 1.000 | 0.998 | 1.000 | 0.378 | 0.374 | 0.993 | 1.000 | 1.000 |
| 22 | 0.454 | 0.997 | 0.845 | 0.206 | 1.000 | 1.000 | 0.991 | 1.000 | 0.109 | 0.993 | 0.110 | 1.000 |
| All | 0.020 | 1.87E-03 | 8.22E-05 | 1.65E-07 | 1.67E-05 | 0.041 | 0.092 | 0.041 | 4.51E-03 | 0.120 | 1.19E-03 | 6.67E-03 |

**Table S6. Counts of pairwise RR effects for alleles that are increasing over time.**

For each pair of traits, the number of markers with positive and negative RR effects are counted for the increasing alleles.

| trait pair | +/+ | +/- | -/+ | -/- | trait pair | +/+ | +/- | -/+ | -/- |
|---|---|---|---|---|---|---|---|---|---|
| FT/LODG | 7,750 | 11,547 | 6,580 | 12,959 | HT/SPWT | 8,336 | 7,027 | 10,862 | 12,611 |
| FT/YLD | 12,996 | 6,301 | 13,251 | 6,288 | HT/TGW | 6,984 | 8,379 | 10,259 | 13,214 |
| FT/HT | 10,552 | 8,745 | 4,811 | 14,728 | HT/EM2 | 7,491 | 7,872 | 12,107 | 11,366 |
| FT/PROT | 8,153 | 11,144 | 6,851 | 12,688 | HT/TILL | 7,068 | 8,295 | 12,408 | 11,065 |
| FT/WK | 6,738 | 12,559 | 11,821 | 7,718 | HT/MAT | 9,382 | 5,981 | 10,916 | 12,557 |
| FT/AWNS | 7,926 | 11,371 | 11,880 | 7,659 | PROT/WK | 6,824 | 8,180 | 11,735 | 12,097 |
| FT/SPWT | 9,474 | 9,823 | 9,724 | 9,815 | PROT/AWNS | 7,855 | 7,149 | 11,951 | 11,881 |
| FT/TGW | 6,972 | 12,325 | 10,271 | 9,268 | PROT/SPWT | 8,430 | 6,574 | 10,768 | 13,064 |
| FT/EM2 | 9,472 | 9,825 | 10,126 | 9,413 | PROT/TGW | 6,380 | 8,624 | 10,863 | 12,969 |
| FT/TILL | 8,408 | 10,889 | 11,068 | 8,471 | PROT/EM2 | 6,887 | 8,117 | 12,711 | 11,121 |
| FT/MAT | 15,617 | 3,680 | 4,681 | 14,858 | PROT/TILL | 7,520 | 7,484 | 11,956 | 11,876 |
| LODG/YLD | 8,729 | 5,601 | 17,518 | 6,988 | PROT/MAT | 8,019 | 6,985 | 12,279 | 11,553 |
| LODG/HT | 8,999 | 5,331 | 6,364 | 18,142 | WK/AWNS | 9,824 | 8,735 | 9,982 | 10,295 |
| LODG/PROT | 5,832 | 8,498 | 9,172 | 15,334 | WK/SPWT | 8,788 | 9,771 | 10,410 | 9,867 |
| LODG/WK | 6,501 | 7,829 | 12,058 | 12,448 | WK/TGW | 8,666 | 9,893 | 8,577 | 11,700 |
| LODG/AWNS | 7,201 | 7,129 | 12,605 | 11,901 | WK/EM2 | 9,439 | 9,120 | 10,159 | 10,118 |
| LODG/SPWT | 7,255 | 7,075 | 11,943 | 12,563 | WK/TILL | 9,156 | 9,403 | 10,320 | 9,957 |
| LODG/TGW | 6,624 | 7,706 | 10,619 | 13,887 | WK/MAT | 8,542 | 10,017 | 11,756 | 8,521 |
| LODG/EM2 | 6,883 | 7,447 | 12,715 | 11,791 | AWNS/SPWT | 10,706 | 9,100 | 8,492 | 10,538 |
| LODG/TILL | 7,411 | 6,919 | 12,065 | 12,441 | AWNS/TGW | 9,840 | 9,966 | 7,403 | 11,627 |
| LODG/MAT | 7,390 | 6,940 | 12,908 | 11,598 | AWNS/EM2 | 10,468 | 9,338 | 9,130 | 9,900 |
| YLD/HT | 9,905 | 16,342 | 5,458 | 7,131 | AWNS/TILL | 11,302 | 8,504 | 8,174 | 10,856 |
| YLD/PROT | 6,336 | 19,911 | 8,668 | 3,921 | AWNS/MAT | 8,454 | 11,352 | 11,844 | 7,186 |
| YLD/WK | 11,905 | 14,342 | 6,654 | 5,935 | SPWT/TGW | 8,588 | 10,610 | 8,655 | 10,983 |
| YLD/AWNS | 13,387 | 12,860 | 6,419 | 6,170 | SPWT/EM2 | 10,571 | 8,627 | 9,027 | 10,611 |
| YLD/SPWT | 13,237 | 13,010 | 5,961 | 6,628 | SPWT/TILL | 9,178 | 10,020 | 10,298 | 9,340 |
| YLD/TGW | 12,504 | 13,743 | 4,739 | 7,850 | SPWT/MAT | 9,157 | 10,041 | 11,141 | 8,497 |
| YLD/EM2 | 13,891 | 12,356 | 5,707 | 6,882 | TGW/EM2 | 6,917 | 10,326 | 12,681 | 8,912 |
| YLD/TILL | 12,962 | 13,285 | 6,514 | 6,075 | TGW/TILL | 9,080 | 8,163 | 10,396 | 11,197 |
| YLD/MAT | 14,055 | 12,192 | 6,243 | 6,346 | TGW/MAT | 7,533 | 9,710 | 12,765 | 8,828 |
| HT/PROT | 7,220 | 8,143 | 7,784 | 15,689 | EM2/TILL | 9,843 | 9,755 | 9,633 | 9,605 |
| HT/WK | 5,846 | 9,517 | 12,713 | 10,760 | EM2/MAT | 9,964 | 9,634 | 10,334 | 8,904 |
| HT/AWNS | 7,036 | 8,327 | 12,770 | 10,703 | TILL/MAT | 9,852 | 9,624 | 10,446 | 8,914 |

**Table S7. Counts of pairwise LASSO effects for alleles that are increasing over time.**

For each pair of traits, the number of markers with positive and negative LASSO effects are counted for the increasing alleles. Significance for each pair of traits (contingency table) is determined from a $\chi^2_{df=1}$ distribution.

| trait pair | +/+ | +/- | -/+ | -/- | $-\log_{10}p$ | trait pair | +/+ | +/- | -/+ | -/- | $-\log_{10}p$ |
|---|---|---|---|---|---|---|---|---|---|---|---|
| FT/LODG | 90 | 223 | 41 | 163 | 1.456 | HT/SPWT | 71 | 63 | 149 | 234 | 2.206 |
| FT/YLD | 270 | 43 | 145 | 59 | 4.435 | HT/TGW | 68 | 66 | 148 | 235 | 1.719 |
| FT/HT | 112 | 201 | 22 | 182 | 9.352 | HT/EM2 | 56 | 78 | 214 | 169 | 2.170 |
| FT/PROT | 77 | 236 | 59 | 145 | 0.491 | HT/TILL | 74 | 60 | 212 | 171 | 0.000 |
| FT/WK | 135 | 178 | 135 | 69 | 6.326 | HT/MAT | 102 | 32 | 223 | 160 | 3.474 |
| FT/AWNS | 139 | 174 | 140 | 64 | 6.959 | PROT/WK | 57 | 79 | 213 | 168 | 2.165 |
| FT/SPWT | 117 | 196 | 103 | 101 | 2.367 | PROT/AWNS | 67 | 69 | 212 | 169 | 0.624 |
| FT/TGW | 120 | 193 | 96 | 108 | 1.215 | PROT/SPWT | 82 | 54 | 138 | 243 | 5.742 |
| FT/EM2 | 151 | 162 | 119 | 85 | 1.506 | PROT/TGW | 64 | 72 | 152 | 229 | 0.754 |
| FT/TILL | 170 | 143 | 116 | 88 | 0.200 | PROT/EM2 | 64 | 72 | 206 | 175 | 0.717 |
| FT/MAT | 268 | 45 | 57 | 147 | 38.908 | PROT/TILL | 77 | 59 | 209 | 172 | 0.097 |
| LODG/YLD | 101 | 30 | 314 | 72 | 0.452 | PROT/MAT | 81 | 55 | 244 | 137 | 0.388 |
| LODG/HT | 75 | 56 | 59 | 327 | 20.083 | WK/AWNS | 145 | 125 | 134 | 113 | 0.013 |
| LODG/PROT | 40 | 91 | 96 | 290 | 0.607 | WK/SPWT | 96 | 174 | 124 | 123 | 2.977 |
| LODG/WK | 64 | 67 | 206 | 180 | 0.368 | WK/TGW | 124 | 146 | 92 | 155 | 1.250 |
| LODG/AWNS | 55 | 76 | 224 | 162 | 2.688 | WK/EM2 | 148 | 122 | 122 | 125 | 0.598 |
| LODG/SPWT | 55 | 76 | 165 | 221 | 0.018 | WK/TILL | 166 | 104 | 120 | 127 | 2.370 |
| LODG/TGW | 67 | 64 | 149 | 237 | 1.801 | WK/MAT | 161 | 109 | 164 | 83 | 0.874 |
| LODG/EM2 | 59 | 72 | 211 | 175 | 1.148 | AWNS/SPWT | 135 | 144 | 85 | 153 | 2.313 |
| LODG/TILL | 77 | 54 | 209 | 177 | 0.385 | AWNS/TGW | 119 | 160 | 97 | 141 | 0.137 |
| LODG/MAT | 83 | 48 | 242 | 144 | 0.011 | AWNS/EM2 | 160 | 119 | 110 | 128 | 1.829 |
| YLD/HT | 106 | 309 | 28 | 74 | 0.103 | AWNS/TILL | 169 | 110 | 117 | 121 | 1.922 |
| YLD/PROT | 74 | 341 | 62 | 40 | 17.486 | AWNS/MAT | 145 | 134 | 180 | 58 | 7.317 |
| YLD/WK | 213 | 202 | 57 | 45 | 0.324 | SPWT/TGW | 85 | 135 | 131 | 166 | 0.607 |
| YLD/AWNS | 223 | 192 | 56 | 46 | 0.036 | SPWT/EM2 | 148 | 72 | 122 | 175 | 8.195 |
| YLD/SPWT | 176 | 239 | 44 | 58 | 0.007 | SPWT/TILL | 105 | 115 | 181 | 116 | 2.427 |
| YLD/TGW | 170 | 245 | 46 | 56 | 0.286 | SPWT/MAT | 119 | 101 | 206 | 91 | 3.269 |
| YLD/EM2 | 228 | 187 | 42 | 60 | 1.765 | TGW/EM2 | 85 | 131 | 185 | 116 | 5.962 |
| YLD/TILL | 230 | 185 | 56 | 46 | 0.000 | TGW/TILL | 122 | 94 | 164 | 137 | 0.144 |
| YLD/MAT | 280 | 135 | 45 | 57 | 4.687 | TGW/MAT | 125 | 91 | 200 | 101 | 1.239 |
| HT/PROT | 56 | 78 | 80 | 303 | 5.408 | EM2/TILL | 146 | 124 | 140 | 107 | 0.213 |
| HT/WK | 34 | 100 | 236 | 147 | 11.996 | EM2/MAT | 149 | 121 | 176 | 71 | 3.643 |
| HT/AWNS | 49 | 85 | 230 | 153 | 5.361 | TILL/MAT | 177 | 109 | 148 | 83 | 0.170 |

**Table S8. Selection gradients, responses, differentials and intensities.**

Selection gradients ($\beta_{sel}$) were estimated from the multivariate breeder's equation of $\Delta Z = G\beta_{sel}$ using the genetic variance-covariance ($G$) calculated from the 12 traits. Selection responses ($\Delta Z$) were estimated from regressing the traits over year of release. Selection differentials ($S$) and intensity ($i$) were calculated from $S = P\beta_{sel}$ and $i = S/\sqrt{diag(P)}$ where $P$ is the phenotypic variance-covariance matrix. The estimated values were partitioned into direct and indirect components due to the direct contributions from selection on the trait itself or indirect contributions from selection on other correlated traits.

| Data | Trait | $\beta_{sel}$ | $\Delta Z$ | | | $S$ | | | $i$ | | |
|---|---|---|---|---|---|---|---|---|---|---|---|
| | | | Total | Direct | Indirect | Total | Direct | Indirect | Total | Direct | Indirect |
| DArT | FT | -0.578 | -0.017 | -1.383 | 1.366 | -0.199 | -2.824 | 2.626 | -0.090 | -1.277 | 1.188 |
| | LODG | 0.877 | -0.033 | 0.085 | -0.118 | 0.156 | 0.323 | -0.167 | 0.257 | 0.532 | -0.275 |
| | YLD | 0.233 | 0.365 | 1.221 | -0.856 | 2.941 | 3.934 | -0.993 | 0.716 | 0.957 | -0.242 |
| | HT | -0.024 | -0.312 | -0.249 | -0.064 | -0.071 | -0.624 | 0.553 | -0.014 | -0.123 | 0.109 |
| | PROT | 1.176 | -0.028 | 0.133 | -0.161 | -0.093 | 0.323 | -0.416 | -0.178 | 0.616 | -0.794 |
| | WK | -0.563 | -0.004 | -0.208 | 0.203 | -0.410 | -0.530 | 0.120 | -0.422 | -0.546 | 0.124 |
| | AWNS | 0.977 | 0.003 | 0.025 | -0.022 | 0.001 | 0.040 | -0.039 | 0.006 | 0.198 | -0.192 |
| | SPWT | -0.210 | -0.001 | -0.100 | 0.099 | -0.288 | -0.622 | 0.335 | -0.167 | -0.362 | 0.194 |
| | TGW | 0.114 | -0.017 | 0.077 | -0.094 | 0.402 | 0.364 | 0.038 | 0.225 | 0.204 | 0.021 |
| | EM2 | 0.023 | 0.101 | 4.211 | -4.110 | 17.103 | 18.234 | -1.131 | 0.604 | 0.644 | -0.040 |
| | TILL | -2.433 | 0.001 | -0.020 | 0.021 | -0.118 | -0.180 | 0.062 | -0.435 | -0.662 | 0.228 |
| | MAT | 2.029 | -0.001 | 0.352 | -0.353 | 0.340 | 0.874 | -0.534 | 0.518 | 1.332 | -0.814 |
| GBS | FT | -0.155 | -0.017 | -0.368 | 0.351 | 0.721 | -0.775 | 1.496 | 0.323 | -0.347 | 0.670 |
| | LODG | -1.116 | -0.033 | -0.090 | 0.058 | -0.335 | -0.412 | 0.076 | -0.552 | -0.678 | 0.126 |
| | YLD | 0.071 | 0.352 | 0.341 | 0.011 | 2.761 | 1.092 | 1.669 | 0.702 | 0.278 | 0.424 |
| | HT | 0.071 | -0.312 | 0.772 | -1.084 | 0.509 | 1.753 | -1.244 | 0.103 | 0.354 | -0.251 |
| | PROT | -0.934 | -0.028 | -0.092 | 0.064 | -0.252 | -0.238 | -0.014 | -0.500 | -0.472 | -0.028 |
| | WK | -0.317 | -0.004 | -0.094 | 0.089 | -0.472 | -0.290 | -0.182 | -0.493 | -0.303 | -0.190 |
| | AWNS | -0.591 | 0.003 | -0.014 | 0.017 | -0.031 | -0.022 | -0.010 | -0.164 | -0.113 | -0.052 |
| | SPWT | 0.471 | -0.001 | 0.266 | -0.267 | 1.269 | 1.344 | -0.075 | 0.752 | 0.796 | -0.044 |
| | TGW | -0.081 | -0.017 | -0.054 | 0.037 | -0.061 | -0.254 | 0.194 | -0.034 | -0.143 | 0.109 |
| | EM2 | -0.010 | 0.101 | -2.529 | 2.630 | -3.661 | -7.747 | 4.086 | -0.132 | -0.279 | 0.147 |
| | TILL | 0.355 | 0.001 | 0.003 | -0.002 | -0.007 | 0.026 | -0.033 | -0.026 | 0.096 | -0.122 |
| | MAT | 0.777 | -0.001 | 0.125 | -0.126 | 0.323 | 0.330 | -0.006 | 0.496 | 0.506 | -0.010 |

**Table S9. Correlations of multivariate breeder's equation variables in simulated data.**

Correlations of variables were calculated from 100 simulations between true/expected and estimated values, and between true/expected and realized values. Estimated values are derived from mixed model G and P matrices while realized values are derived from true G and P matrices. G: genetic variance-covariance (vcov), E: residual vcov, P: phenotypic vcov, Z: selection response, β: selection gradient, S: selection differential, i: selection intensity.

| Variable | | Correlation (true vs. estimated values) | | | Correlation (true vs. realized values) | | |
|---|---|---|---|---|---|---|---|
| | | Trait 1 | Trait 2 | Trait 3 | Trait 1 | Trait 2 | Trait 3 |
| G | Trait 1 | - | - | - | - | - | - |
| | Trait 2 | 0.417 | - | - | - | - | - |
| | Trait 3 | 0.326 | 0.407 | - | - | - | - |
| E | Trait 1 | 0.484 | - | - | - | - | - |
| | Trait 2 | 0.618 | 0.570 | - | - | - | - |
| | Trait 3 | 0.446 | 0.518 | 0.566 | - | - | - |
| Sel1 | Z | - | - | - | 0.144 | 0.857 | 0.721 |
| | Z, direct | 0.230 | 0.346 | 0.314 | 0.069 | 0.611 | 0.449 |
| | Z, indirect | 0.190 | 0.213 | 0.315 | 0.550 | 0.756 | 0.843 |
| | β | 0.402 | 0.360 | 0.331 | 0.069 | 0.611 | 0.449 |
| | S | 0.095 | 0.460 | 0.308 | 0.522 | 0.795 | 0.605 |
| | S, direct | 0.083 | 0.344 | 0.317 | 0.441 | 0.618 | 0.451 |
| | S, indirect | 0.294 | 0.367 | 0.385 | 0.591 | 0.837 | 0.879 |
| | i | 0.289 | 0.476 | 0.325 | 0.155 | 0.791 | 0.603 |
| | i, direct | 0.270 | 0.352 | 0.323 | 0.154 | 0.615 | 0.449 |
| | i, indirect | 0.294 | 0.373 | 0.394 | 0.593 | 0.830 | 0.877 |
| Sel2 | Z | - | - | - | 0.609 | 0.692 | 0.730 |
| | Z, direct | 0.268 | 0.247 | 0.209 | 0.310 | 0.394 | 0.458 |
| | Z, indirect | 0.174 | 0.263 | 0.335 | 0.738 | 0.735 | 0.779 |
| | β | 0.314 | 0.223 | 0.200 | 0.310 | 0.394 | 0.458 |
| | S | 0.304 | 0.182 | 0.292 | 0.637 | 0.614 | 0.627 |
| | S, direct | 0.204 | 0.026 | 0.194 | 0.450 | 0.434 | 0.448 |
| | S, indirect | 0.375 | 0.422 | 0.333 | 0.759 | 0.827 | 0.839 |
| | i | 0.358 | 0.268 | 0.321 | 0.554 | 0.581 | 0.637 |
| | i, direct | 0.257 | 0.112 | 0.196 | 0.360 | 0.405 | 0.453 |
| | i, indirect | 0.351 | 0.421 | 0.334 | 0.753 | 0.822 | 0.831 |
| Sel3 | Z | - | - | - | 0.546 | 0.612 | 0.448 |
| | Z, direct | 0.087 | 0.205 | 0.041 | 0.229 | 0.359 | 0.181 |
| | Z, indirect | 0.122 | 0.101 | 0.355 | 0.731 | 0.543 | 0.677 |
| | β | 0.121 | 0.155 | 0.084 | 0.229 | 0.359 | 0.181 |
| | S | 0.119 | 0.197 | 0.030 | 0.521 | 0.559 | 0.342 |
| | S, direct | 0.052 | 0.073 | -0.015 | 0.313 | 0.409 | 0.209 |
| | S, indirect | 0.390 | 0.287 | 0.309 | 0.810 | 0.694 | 0.740 |
| | i | 0.130 | 0.253 | 0.086 | 0.450 | 0.514 | 0.280 |
| | i, direct | 0.076 | 0.107 | 0.029 | 0.257 | 0.378 | 0.182 |
| | i, indirect | 0.380 | 0.279 | 0.314 | 0.798 | 0.682 | 0.735 |

**Table S10. Stability of the genetic variance-covariance (G) and correlation (g) matrices across years of release and countries of origin.**

The matrices are compared pairwise using Random Skewers (Cheverud and Marroig 2007) and Mantel test (Mantel 1967). For Random Skewers, the results are shown as the correlations between the predicted responses from the two matrices and the p-values from 1,000 permutations. For Mantel test, the results are shown as the correlations between the off-diagonal elements of the two matrices and the p-values from 10,000 permutations. In both tests, a significance indicates similarity between the two matrices.

| Matrix 1 | Matrix 2 | Random Skewers | | Mantel test | |
|---|---|---|---|---|---|
| | | Correlation | P-value | Correlation | P-value |
| $G$ | $G_{DArT}$ | 0.994 | 0 | 0.870 | 1e-4 |
| $g$ | $g_{DArT}$ | 0.941 | 0 | 0.899 | 1e-4 |
| $G$ | $G_{DE}$ | 0.924 | 0 | 0.543 | 0.003 |
| $G$ | $G_{FR}$ | 0.979 | 0 | -0.001 | 0.920 |
| $G$ | $G_{UK}$ | 0.974 | 0 | 0.615 | 0.002 |
| $G$ | $G_{y1}$ | 0.984 | 0 | 0.788 | 1e-4 |
| $G$ | $G_{y2}$ | 0.875 | 0 | 0.195 | 0.108 |
| $G$ | $G_{y3}$ | 0.978 | 0 | 0.300 | 0.038 |
| $G_{DE}$ | $G_{FR}$ | 0.887 | 0 | -0.484 | 0.003 |
| $G_{DE}$ | $G_{UK}$ | 0.891 | 0 | -0.013 | 0.432 |
| $G_{FR}$ | $G_{UK}$ | 0.939 | 0 | -0.203 | 0.111 |
| $G_{y1}$ | $G_{y2}$ | 0.856 | 0 | -0.303 | 0.081 |
| $G_{y1}$ | $G_{y3}$ | 0.977 | 0 | 0.438 | 0.010 |
| $G_{y2}$ | $G_{y3}$ | 0.886 | 0 | -0.331 | 0.029 |
| $g$ | $g_{DE}$ | 0.625 | 0.191 | 0.449 | 0.002 |
| $g$ | $g_{FR}$ | 0.876 | 0 | 0.825 | 1e-4 |
| $g$ | $g_{UK}$ | 0.510 | 0.852 | 0.280 | 0.037 |
| $g$ | $g_{y1}$ | 0.761 | 0 | 0.700 | 1e-4 |
| $g$ | $g_{y2}$ | 0.755 | 0 | 0.624 | 1e-4 |
| $g$ | $g_{y3}$ | 0.660 | 0.049 | 0.693 | 1e-4 |
| $g_{DE}$ | $g_{FR}$ | 0.456 | 0.966 | 0.223 | 0.105 |
| $g_{DE}$ | $g_{UK}$ | 0.271 | 1 | 0.018 | 0.913 |
| $g_{FR}$ | $g_{UK}$ | 0.368 | 1 | 0.108 | 0.430 |
| $g_{y1}$ | $g_{y2}$ | 0.434 | 0.982 | 0.223 | 0.068 |
| $g_{y1}$ | $g_{y3}$ | 0.491 | 0.912 | 0.433 | 1e-4 |
| $g_{y2}$ | $g_{y3}$ | 0.473 | 0.950 | 0.437 | 3e-4 |

**Table S11. Genetic correlation and variance-covariance matrix for 12 traits in the TG panel.**

Genetic correlations, variances (highlighted in grey) and covariances are shown in the top right triangle, diagonal and bottom left triangle, respectively.

| Marker | Trait | FT | LODG | YLD | HT | PROT | WK | AWNS | SPWT | TGW | EM2 | TILL | MAT |
|---|---|---|---|---|---|---|---|---|---|---|---|---|---|
| DArT | FT | 2.394 | 0.226 | 0.074 | 0.409 | -0.070 | -0.510 | -0.215 | -0.254 | -0.476 | -0.039 | 0.059 | 0.867 |
| | LODG | 0.109 | 0.097 | -0.198 | 0.596 | 0.128 | -0.309 | 0.262 | 0.438 | -0.085 | -0.188 | 0.628 | 0.052 |
| | YLD | 0.262 | -0.141 | 5.241 | -0.314 | -0.768 | 0.184 | 0.041 | -0.420 | 0.054 | 0.033 | -0.062 | -0.009 |
| | HT | 2.034 | 0.597 | -2.309 | 10.328 | 0.352 | -0.491 | 0.006 | 0.455 | -0.063 | -0.112 | 0.086 | 0.198 |
| | PROT | -0.036 | 0.013 | -0.591 | 0.381 | 0.113 | -0.228 | 0.024 | 0.643 | 0.061 | -0.181 | 0.151 | -0.053 |
| | WK | -0.479 | -0.059 | 0.256 | -0.959 | -0.047 | 0.369 | 0.056 | -0.134 | 0.121 | 0.275 | 0.224 | -0.187 |
| | AWNS | -0.053 | 0.013 | 0.015 | 0.003 | 0.001 | 0.005 | 0.026 | 0.454 | 0.270 | 0.058 | 0.234 | -0.403 |
| | SPWT | -0.271 | 0.094 | -0.665 | 1.011 | 0.150 | -0.056 | 0.051 | 0.477 | -0.150 | 0.295 | -0.135 | -0.499 |
| | TGW | -0.603 | -0.022 | 0.100 | -0.165 | 0.017 | 0.060 | 0.036 | -0.085 | 0.669 | -0.517 | 0.329 | -0.433 |
| | EM2 | -0.826 | -0.798 | 1.013 | -4.916 | -0.828 | 2.275 | 0.127 | 2.770 | -5.755 | 185.22 | -0.210 | -0.130 |
| | TILL | 0.008 | 0.018 | -0.013 | 0.025 | 0.005 | 0.012 | 0.003 | -0.008 | 0.024 | -0.257 | 0.008 | 0.176 |
| | MAT | 0.559 | 0.007 | -0.009 | 0.266 | -0.007 | -0.047 | -0.027 | -0.144 | -0.148 | -0.738 | 0.007 | 0.174 |
| GBS | FT | 2.367 | 0.498 | -0.077 | 0.388 | 0.053 | -0.432 | -0.284 | -0.205 | -0.492 | -0.065 | -0.009 | 0.808 |
| | LODG | 0.218 | 0.081 | -0.205 | 0.723 | 0.048 | -0.315 | 0.094 | 0.235 | -0.522 | 0.072 | 0.415 | 0.102 |
| | YLD | -0.260 | -0.128 | 4.830 | -0.461 | -0.726 | 0.259 | 0.144 | -0.214 | 0.190 | 0.138 | 0.081 | -0.057 |
| | HT | 1.964 | 0.677 | -3.333 | 10.826 | 0.395 | -0.302 | -0.116 | 0.305 | -0.235 | -0.042 | -0.069 | -0.033 |
| | PROT | 0.026 | 0.004 | -0.500 | 0.408 | 0.098 | -0.285 | -0.079 | 0.498 | -0.096 | -0.202 | -0.140 | -0.047 |
| | WK | -0.361 | -0.049 | 0.309 | -0.540 | -0.049 | 0.295 | 0.020 | -0.021 | 0.130 | 0.253 | -0.111 | -0.095 |
| | AWNS | -0.068 | 0.004 | 0.049 | -0.059 | -0.004 | 0.002 | 0.024 | 0.405 | 0.319 | -0.093 | 0.380 | -0.296 |
| | SPWT | -0.237 | 0.050 | -0.353 | 0.755 | 0.117 | -0.009 | 0.047 | 0.565 | -0.293 | 0.310 | 0.023 | -0.510 |
| | TGW | -0.618 | -0.122 | 0.342 | -0.631 | -0.025 | 0.058 | 0.041 | -0.180 | 0.668 | -0.529 | 0.301 | -0.579 |
| | EM2 | -1.585 | 0.326 | 4.813 | -2.199 | -1.005 | 2.176 | -0.228 | 3.686 | -6.846 | 250.92 | -0.034 | -0.019 |
| | TILL | -0.001 | 0.011 | 0.017 | -0.022 | -0.004 | -0.006 | 0.006 | 0.002 | 0.024 | -0.051 | 0.009 | 0.302 |
| | MAT | 0.500 | 0.012 | -0.050 | -0.044 | -0.006 | -0.021 | -0.019 | -0.154 | -0.190 | -0.118 | 0.012 | 0.162 |

### Supplementary Figures

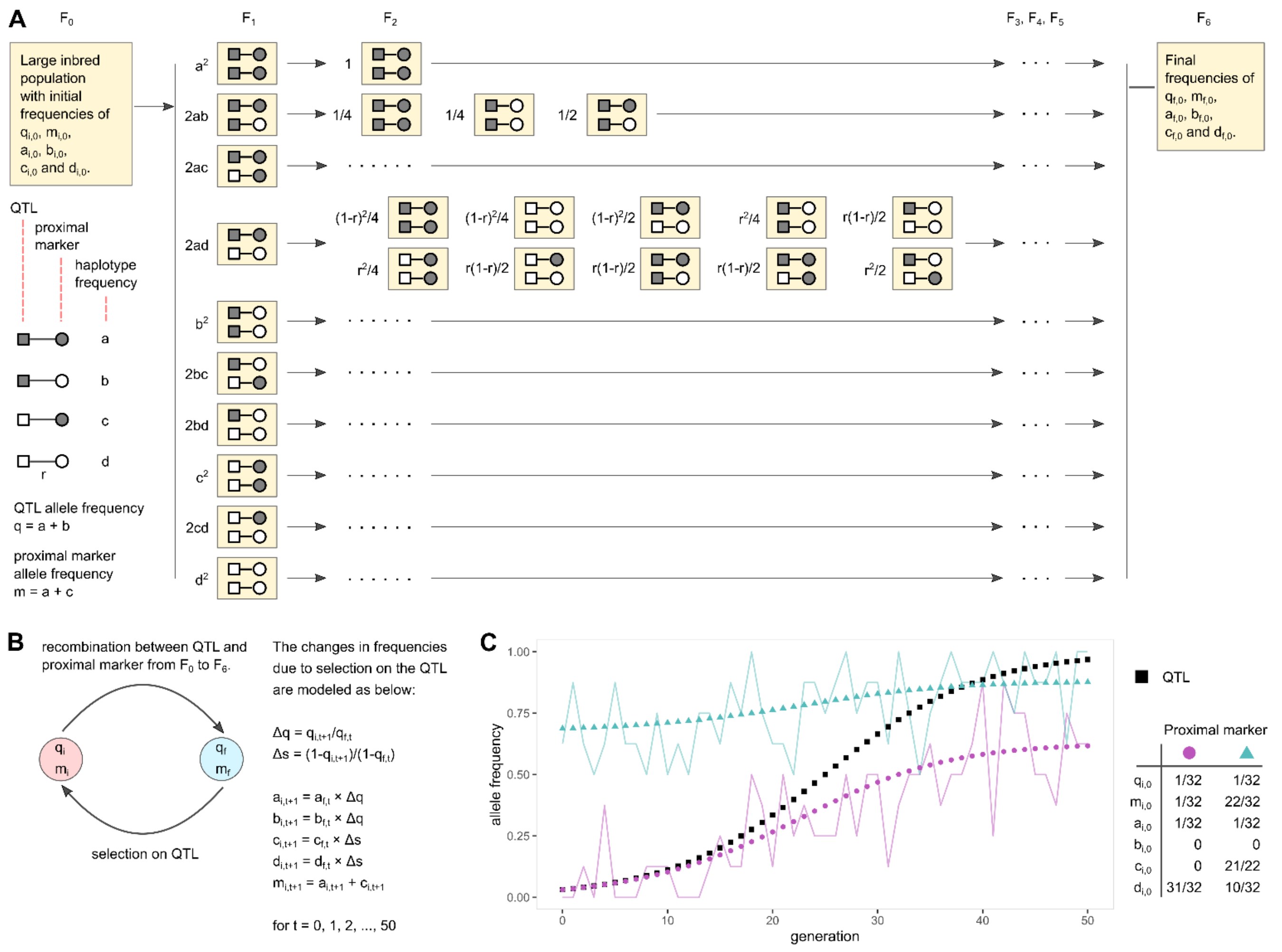


**Figure S1. Modelling of hitch-hiking effect due to selection on QTL.**

We simulated selection on a QTL and observed the possible hitch-hiking effect in the proximal marker. We used the simulation results to determine the extent of RALLY detection limit in marker linked to QTLs under selection. [A] Given the initial frequencies of the QTL allele ($q_i$), proximal marker allele ($m_i$) and QTL-marker haplotypes ($a_i$, $b_i$, $c_i$ and $d_i$) in the parents, we determined the expected frequencies $q_f$, $m_f$, $a_f$, $b_f$, $c_f$ and $d_f$ in the $F_6$ individuals. We tested distances of 1, 2, …, 10 cM between the QTL and proximal marker. [B] By modeling selection directly on the QTL as an increase in QTL allele frequency, we computed the correlated changes in proximal marker allele and QTL-marker haplotypes frequencies. [C] An example of a QTL with an initial frequency of 1/32 rising to 31/32 over 50 generations of selection under a logistic model. Two proximal markers that are 1 cM away from the QTL are considered. Initially, the proximal marker allele shown as purple circle is completely linked to the QTL allele while the proximal marker allele shown as blue triangle is weakly linked to the QTL allele. The line plots show the proximal marker allele frequencies from random sampling of 8 individuals per generation. Under the same sample size, the first marker is more likely to be significant under a logistic regression model than the second marker which is largely invariant over time.

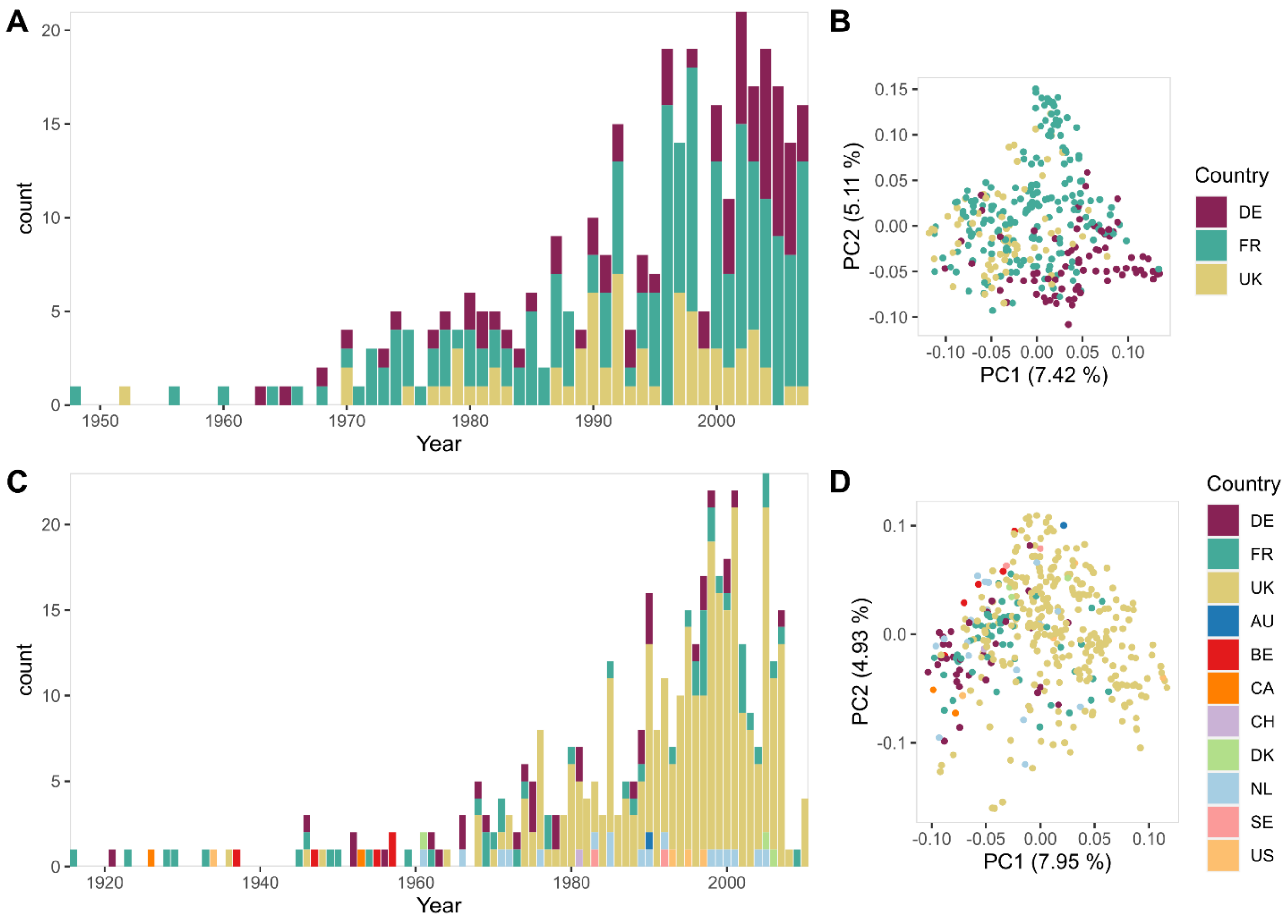


**Figure S2. Distributions of 333 TG and 403 WAGTAIL winter wheat varieties according to years of release and countries of origin.**

[**A**] Counts of TG varieties ranging from 1948 to 2007. [**B**] Plot of first two principal components using TG GBS dataset. [**C**] Counts of WAGTAIL varieties ranging from 1916 to 2010. [**D**] Plot of first two principal components using WAGTAIL dataset.

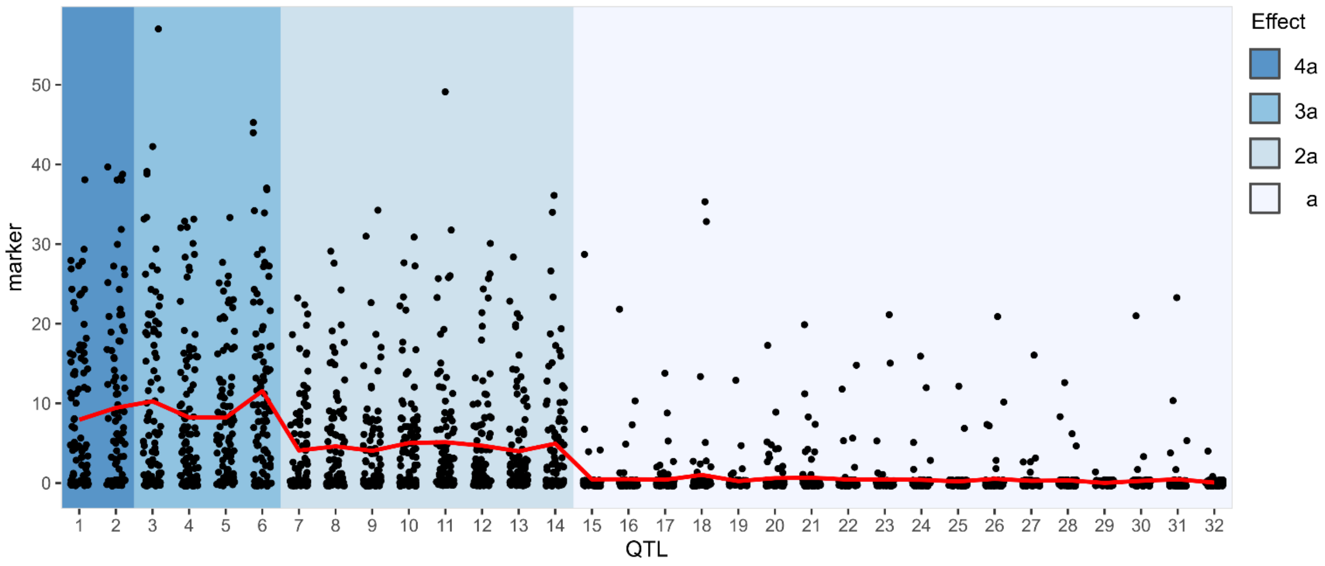


**Figure S3. Number of significant markers across different simulated QTL frequencies and effects in the selected population.**

The QTLs are simulated such that the initial frequencies are 1/32 (QTL-1,2), 2/32 (QTL-3,4), …, and 16/32 (QTL-31,32). Each point represents the number of significant markers within 5 cM of the QTL in a simulation. The red line shows the mean number of significant markers across all 32 QTLs. The number of significant markers drops as the QTL initial frequency increases and effect decreases.

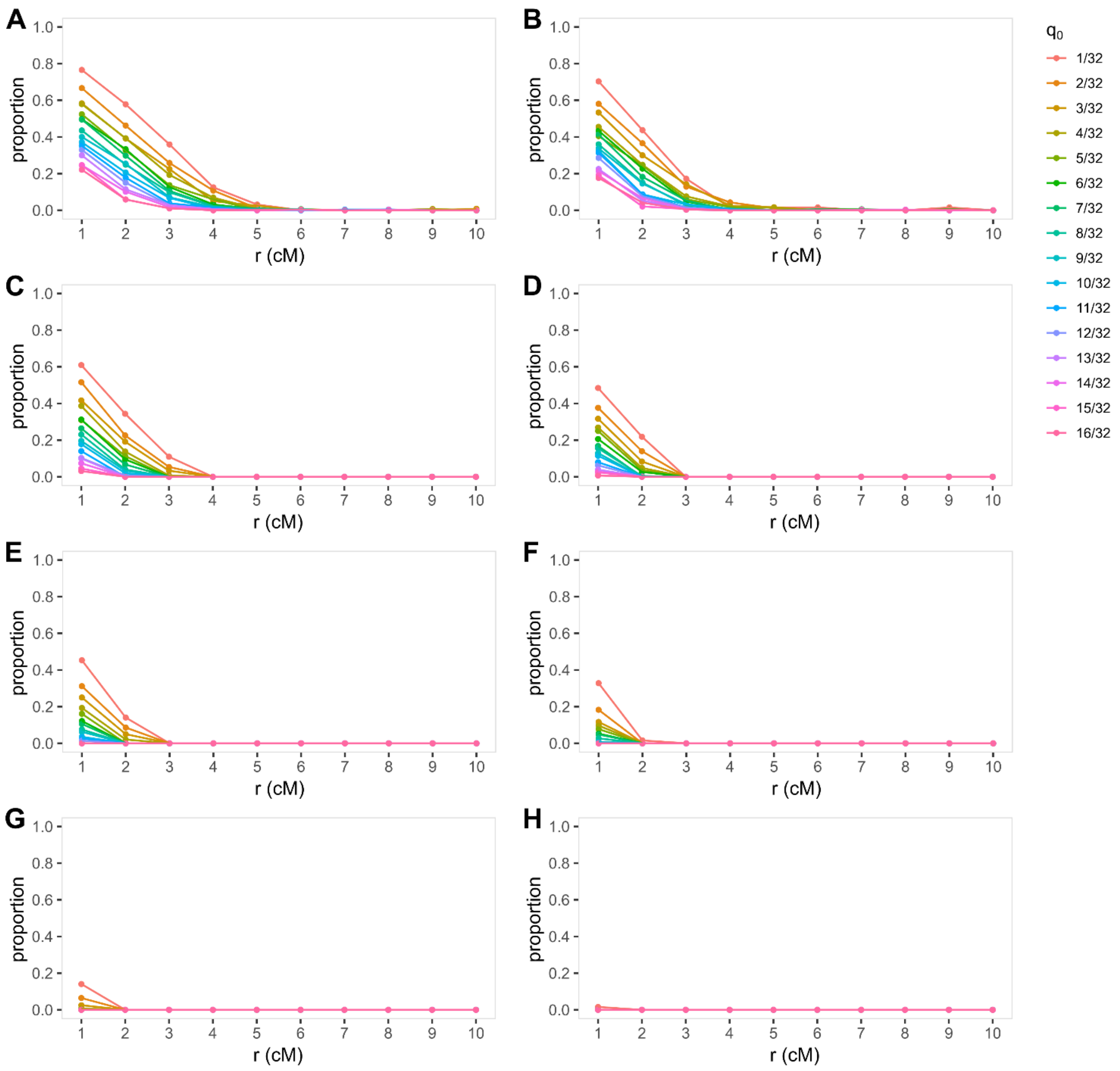


**Figure S4. Detection limit of RALLY.**

We tested the significances of proximal markers at distances of 1 to 10 cM away from a QTL under selection. All possible initial QTL-marker haplotypes were considered for each initial QTL allele frequency ($q_0$), and the proportions of haplotypes that led to significant markers were plotted. The QTL allele frequencies in [**A,C,E,G**] increase under a logistic model while the QTL allele frequencies in [**B,D,F,H**] increase under a linear model. The results are separated by markers that are [**A,B**] significant in at least 1 out of 100 simulations, [**C,D**] significant in at least 10 out of 100 simulations, [**E,F**] significant in at least 50 out of 100 simulations and [**G,H**] significant in all 100 simulations.

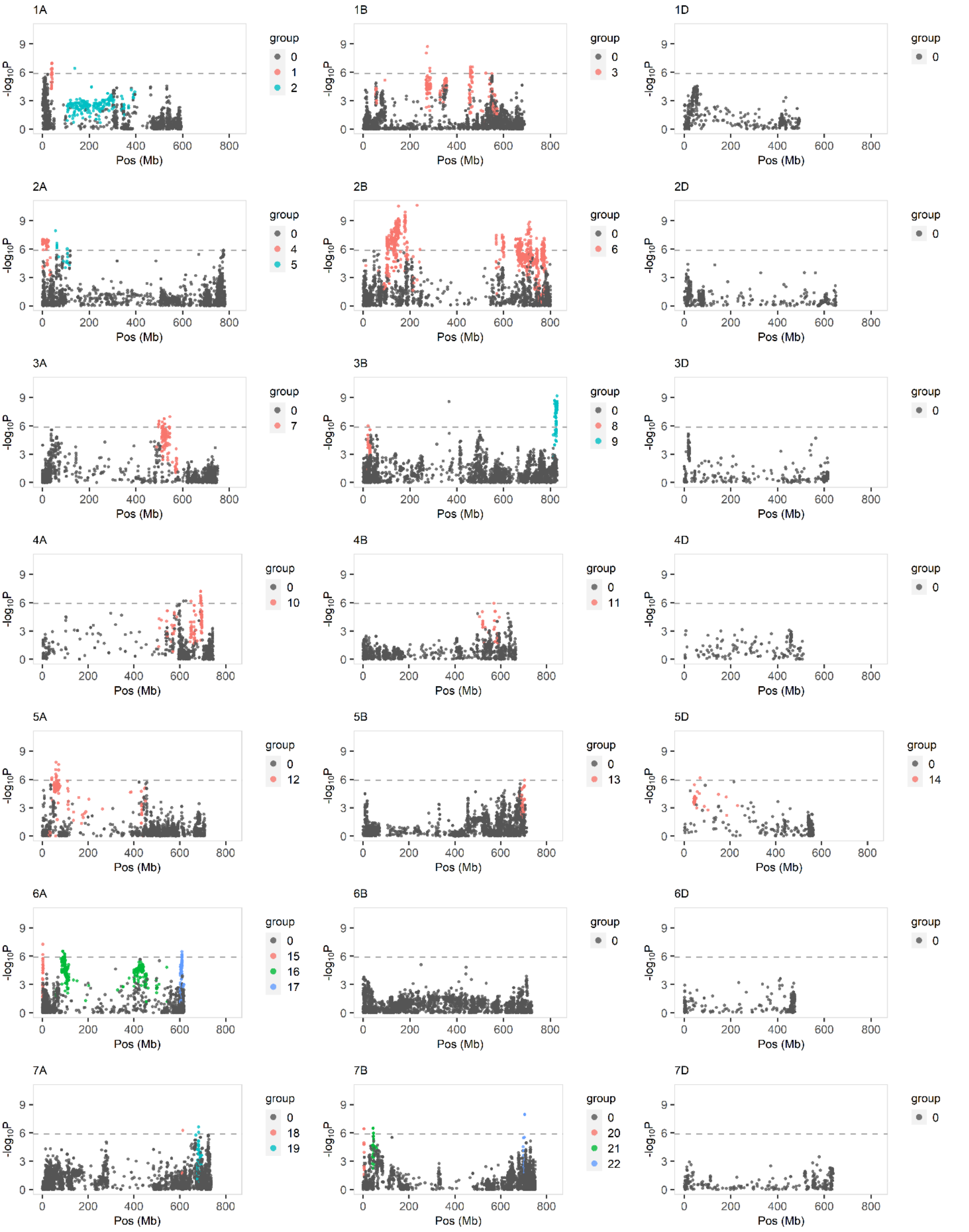


**Figure S5. RALLY QSL groups in TG panel.**

RALLY peaks and their linked markers ($r^2 > 0.2$) are grouped and highlighted. The dashed horizontal line represents the Bonferroni threshold of 0.05.

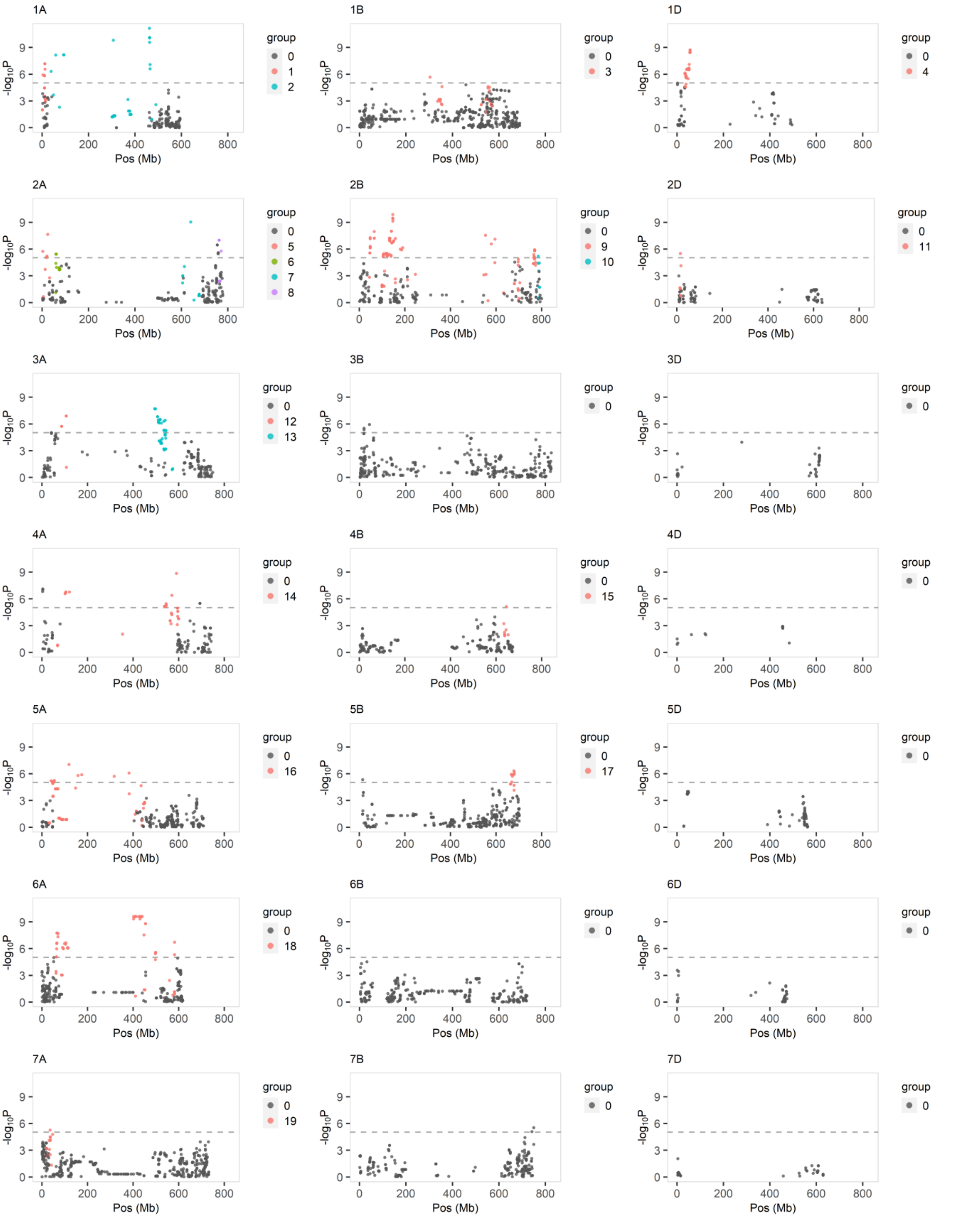


**Figure S6. RALLY QSL groups in WAGTAIL panel.**

RALLY peaks and their linked markers ($r^2 > 0.2$) are grouped and highlighted. The dashed horizontal line represents the Bonferroni threshold of 0.05.

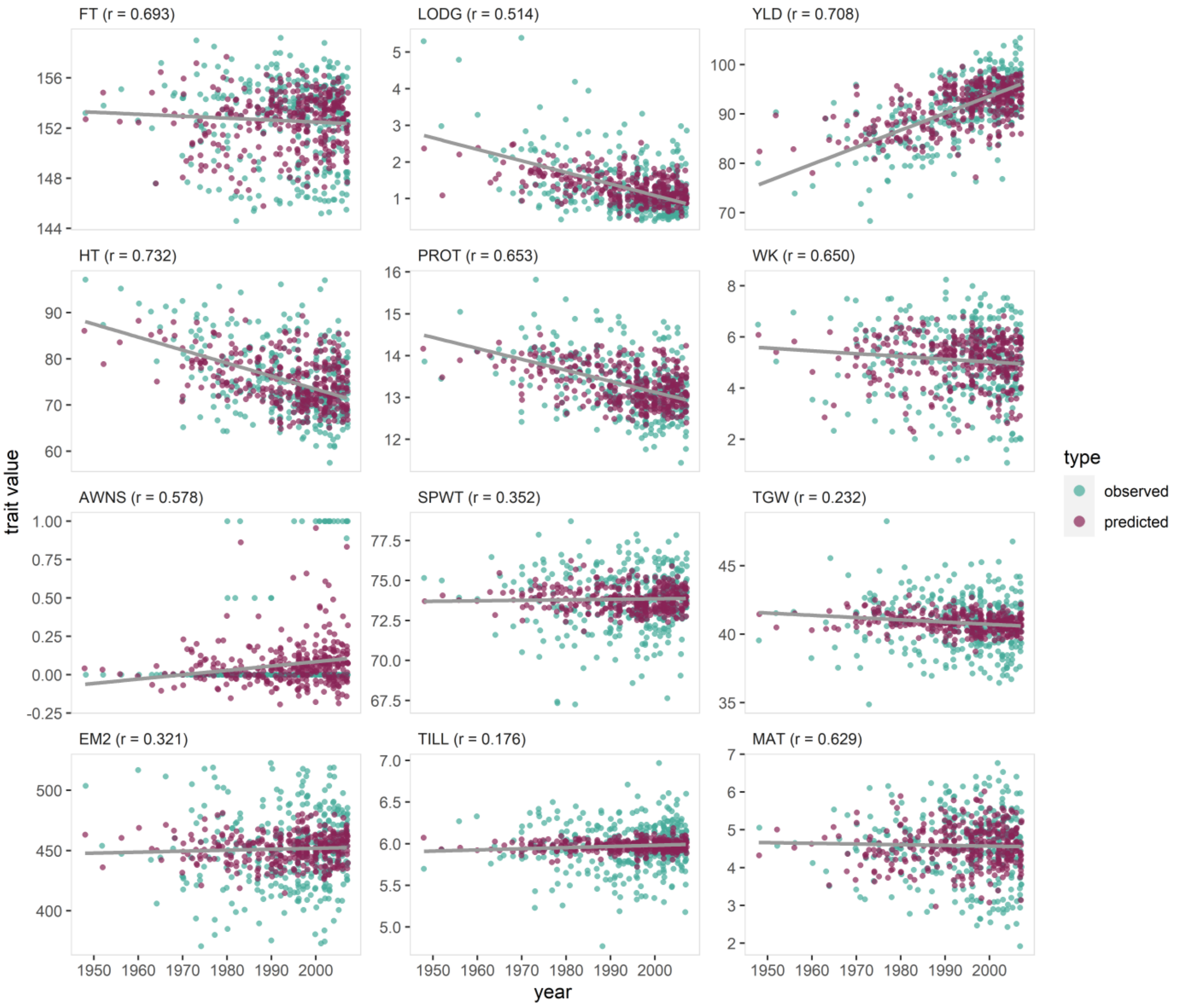


**Figure S7. Predicted trait values using Ridge Regression (RR).**

The traits are predicted using 5-fold cross validations (CVs). The predicted and observed trait values are plotted against the year of release. The correlations between predicted and observed trait values are shown at the top of each subplot. The gray linear regression line is fitted on the observed trait values to summarize the change in phenotypic trait values over the years.

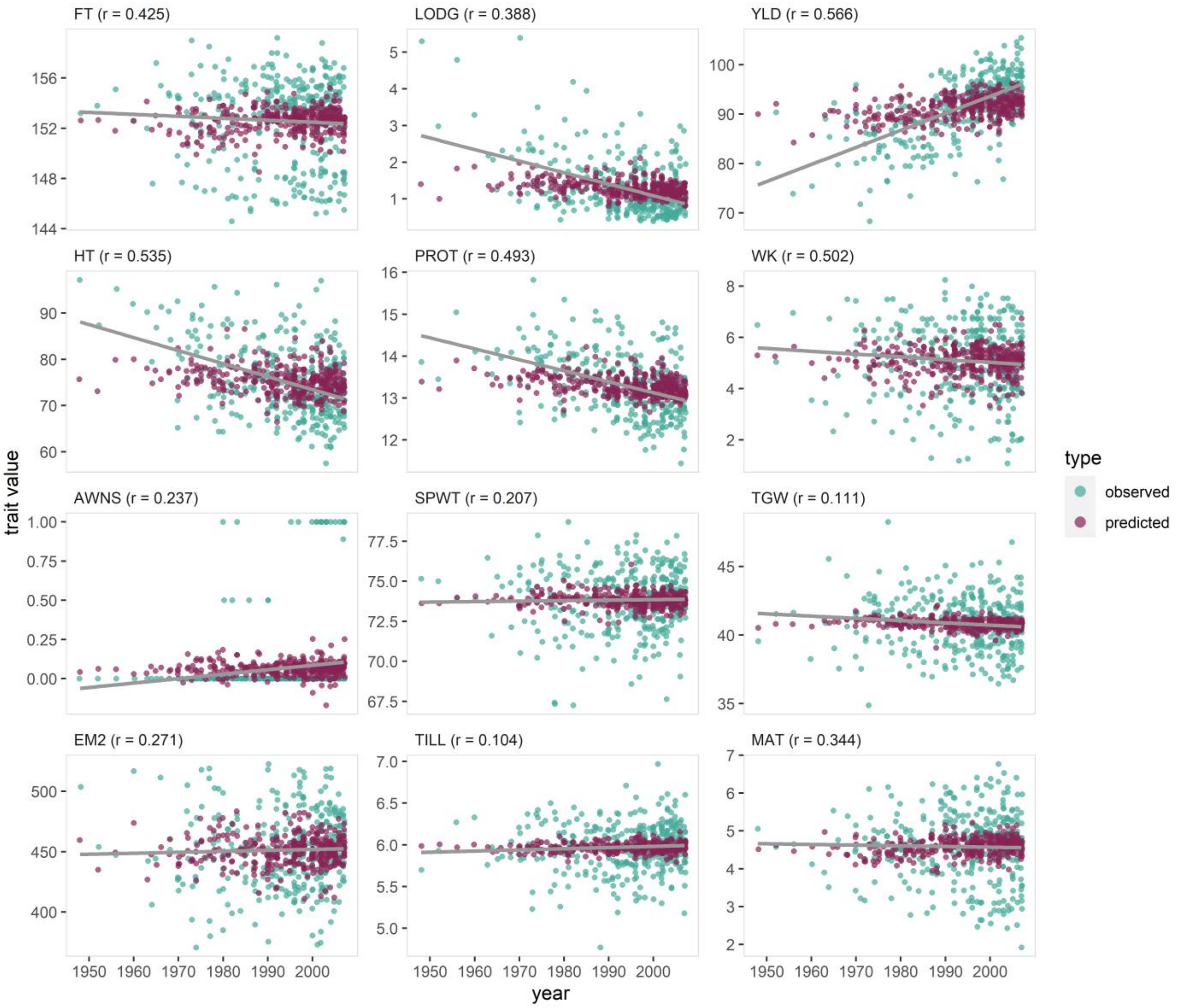


**Figure S8. Predicted trait values using LASSO.**

The traits are predicted using 5-fold cross validations (CVs). The predicted and observed trait values are plotted against the year of release. The correlations between predicted and observed trait values are shown at the top of each subplot. The gray linear regression line is fitted on the observed trait values to summarize the change in phenotypic trait values over the years.

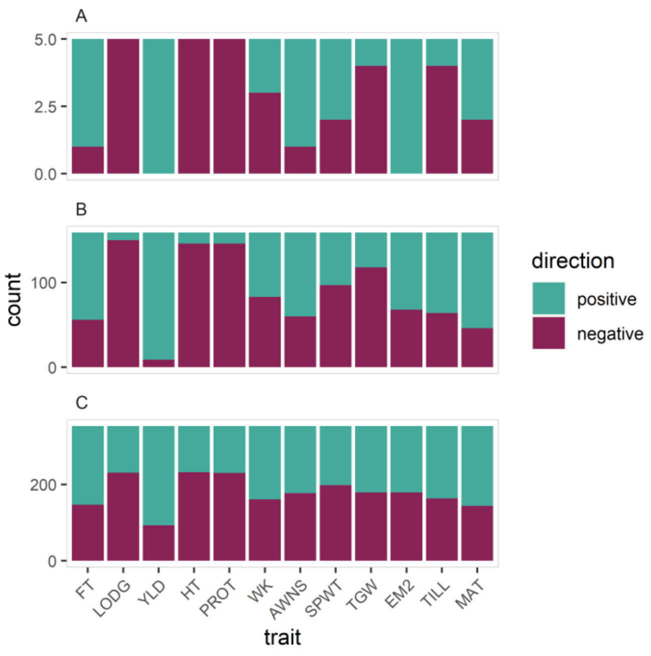


**Figure S9. Distribution of the positive and negative LASSO effects in the increasing alleles.**

[A] Markers with RALLY P-values of lower than the Bonferroni corrected threshold. [B] Markers with RALLY P-values between 0.05 and the Bonferroni corrected threshold. [C] Markers with RALLY P-values of higher than 0.05.

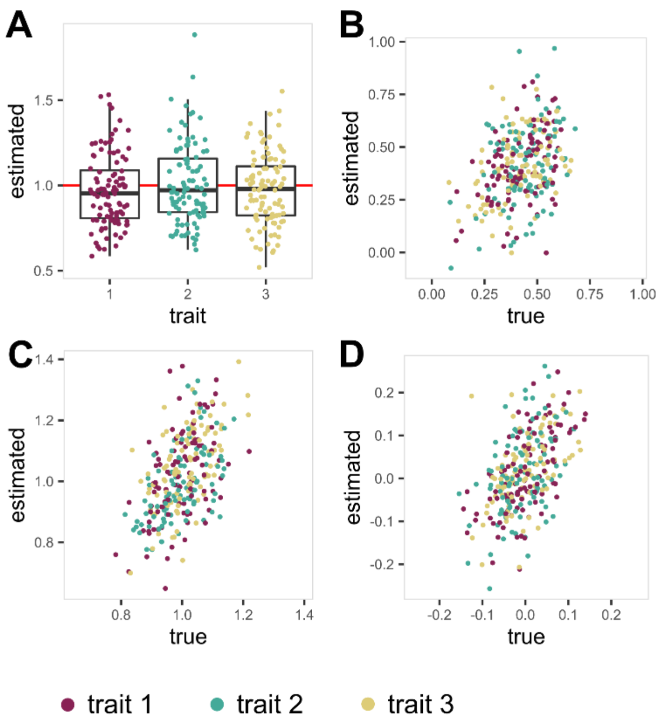


**Figure S10. Genetic and residual variances-covariances in three simulated traits.**

The estimated values from mixed linear model are shown in the Y-axis and the true simulated values are shown in the X-axis in [**B**-**D**]. Similar Y-axis is shown in [**A**] but since the true simulated genetic variances are always 1 (horizontal red line), the traits are shown in the X-axis instead. Each point represents a single simulation with 100 simulations in total. [**A**] Genetic variances, [**B**] Genetic covariances, [**C**] Residual variances, and [**D**] Residual covariances.

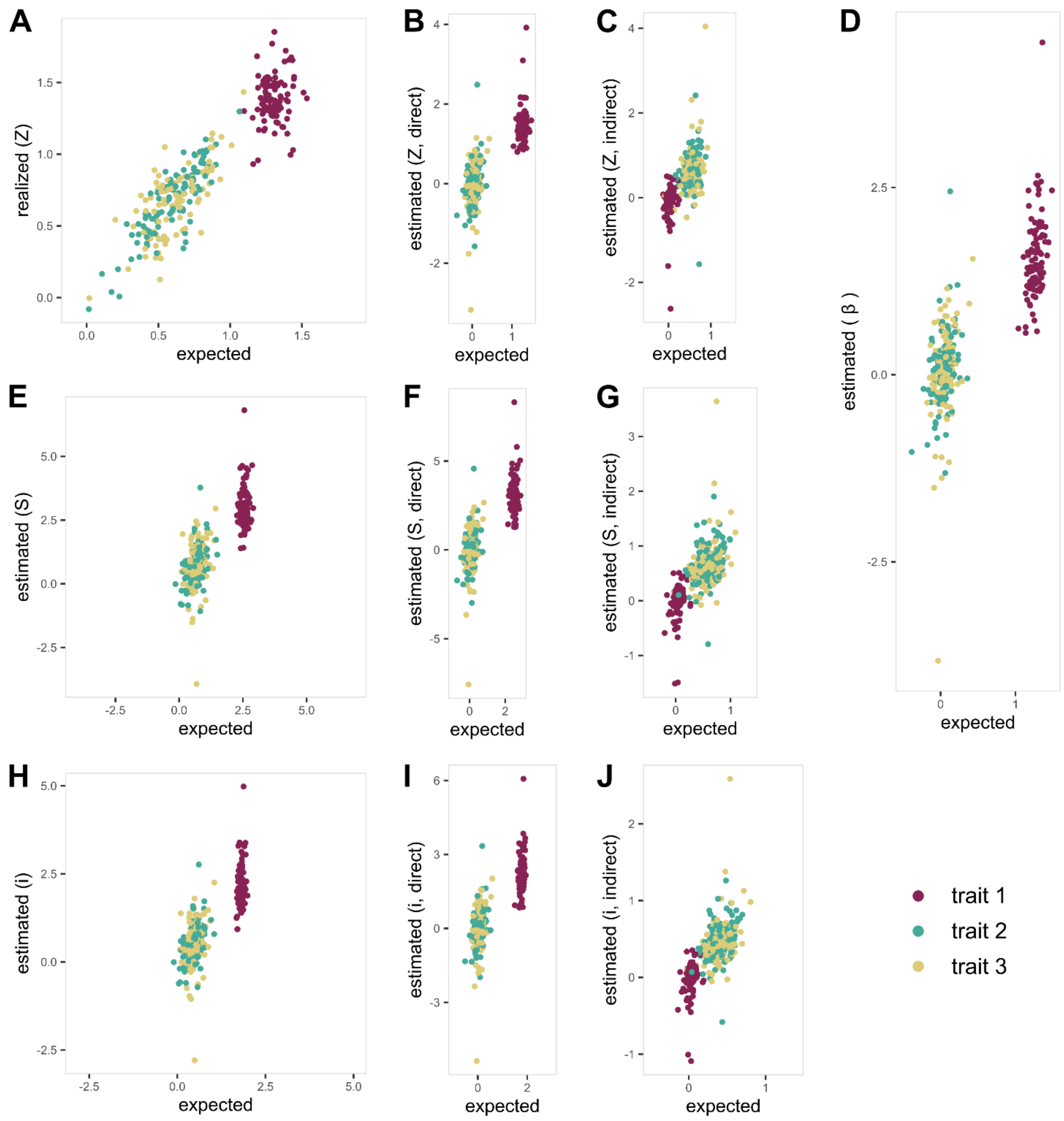


**Figure S11. Estimated multivariate selection parameters in Sel1.**

All estimated multivariate selection parameters are shown in the Y-axis and the true/expected values are shown in the X-axis. Note that selection responses are realized (*same as Figure S12A*) and do not depend on any estimated values from mixed linear model. [**A**] Selection responses, partitioned into [**B**] direct and [**C**] indirect. [**D**] Selection gradient. [**E**] Selection differential, partitioned into [**F**] direct and [**G**] indirect. [**H**] Selection intensity, partitioned into [**I**] direct and [**J**] indirect.

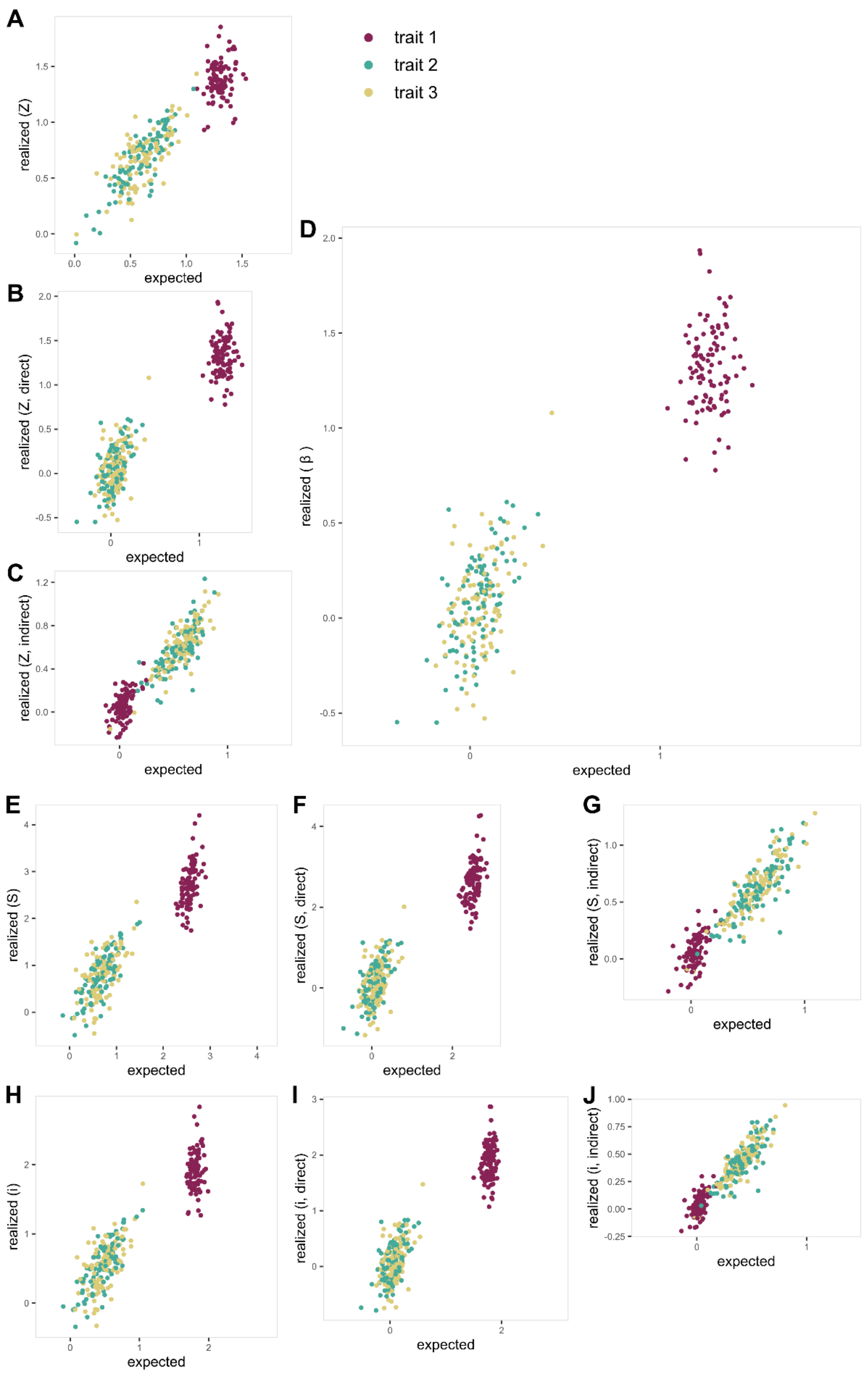


**Figure S12. Realized multivariate selection parameters in Sel1.**

All realized multivariate selection parameters are shown in the Y-axis and the true/expected values are shown in the X-axis. [**A**] Selection responses, partitioned into [**B**] direct and [**C**] indirect. [**D**] Selection gradient. [**E**] Selection differential, partitioned into [**F**] direct and [**G**] indirect. [**H**] Selection intensity, partitioned into [**I**] direct and [**J**] indirect.

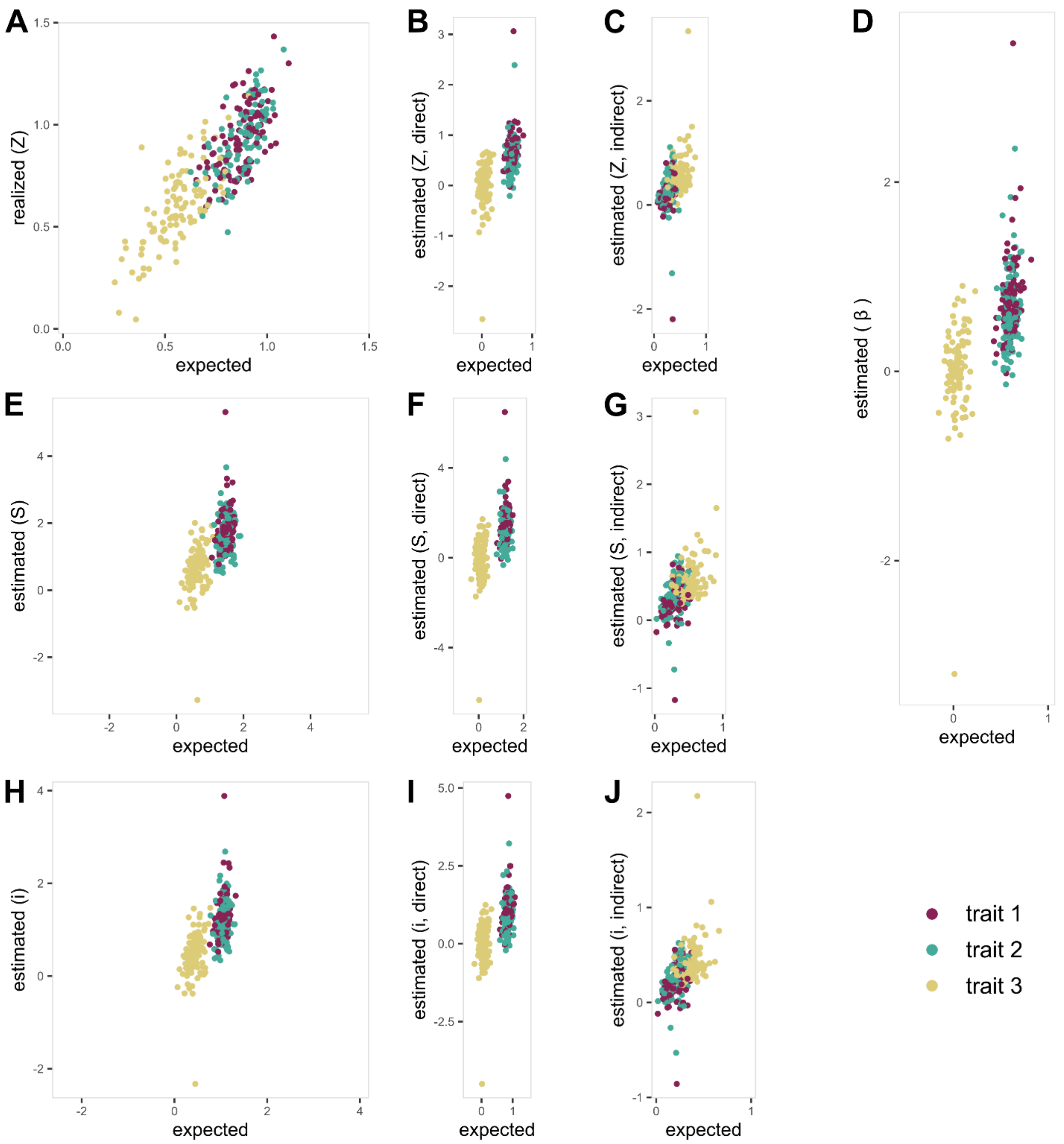


**Figure S13. Estimated multivariate selection parameters in Sel2.**

All estimated multivariate selection parameters are shown in the Y-axis and the true/expected values are shown in the X-axis. Note that selection responses are realized (*same as Figure S14A*) and do not depend on any estimated values from mixed linear model. [**A**] Selection responses, partitioned into [**B**] direct and [**C**] indirect. [**D**] Selection gradient. [**E**] Selection differential, partitioned into [**F**] direct and [**G**] indirect. [**H**] Selection intensity, partitioned into [**I**] direct and [**J**] indirect.

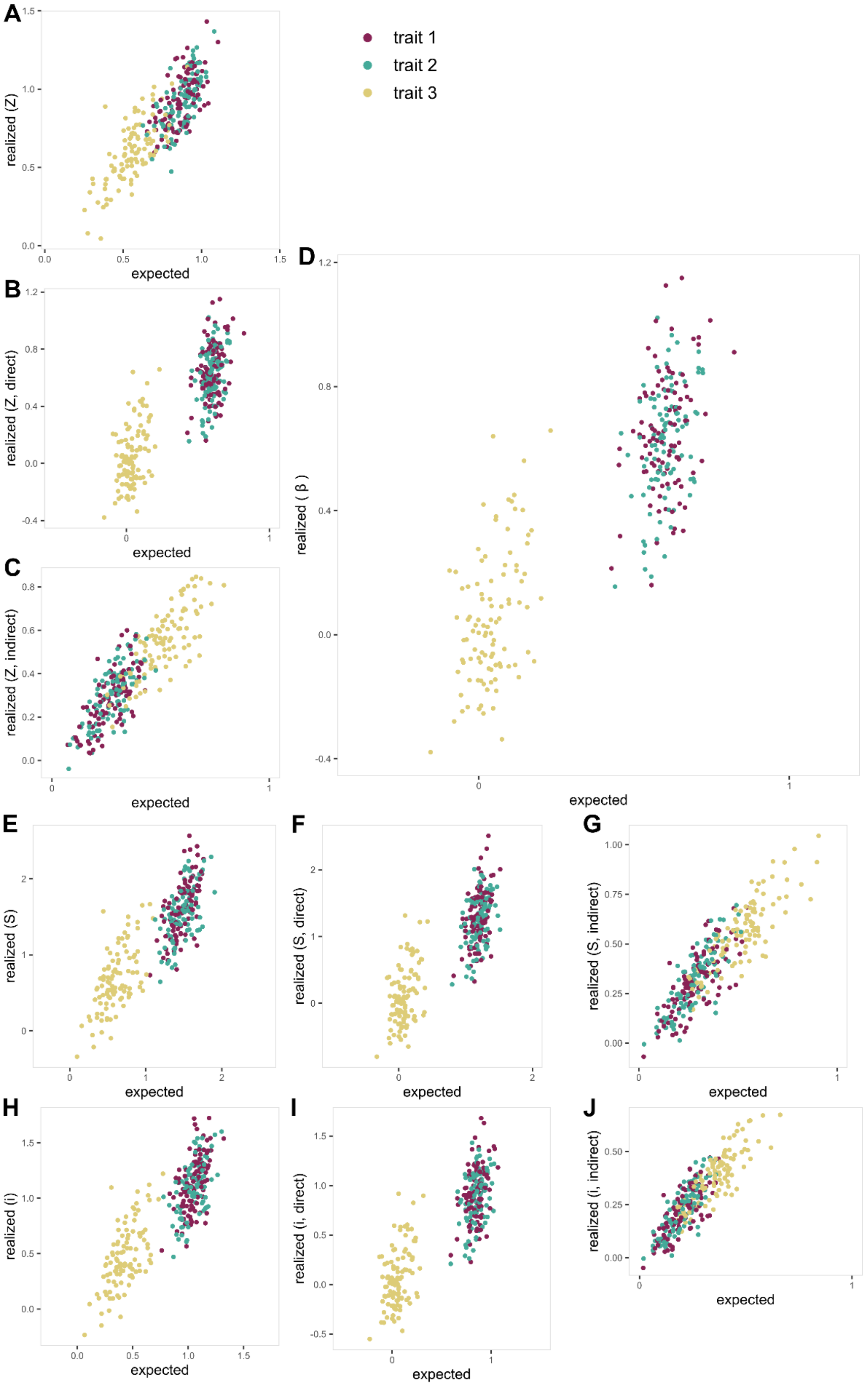


**Figure S14. Realized multivariate selection parameters in Sel2.**

All realized multivariate selection parameters are shown in the Y-axis and the true/expected values are shown in the X-axis. [**A**] Selection responses, partitioned into [**B**] direct and [**C**] indirect. [**D**] Selection gradient. [**E**] Selection differential, partitioned into [**F**] direct and [**G**] indirect. [**H**] Selection intensity, partitioned into [**I**] direct and [**J**] indirect.

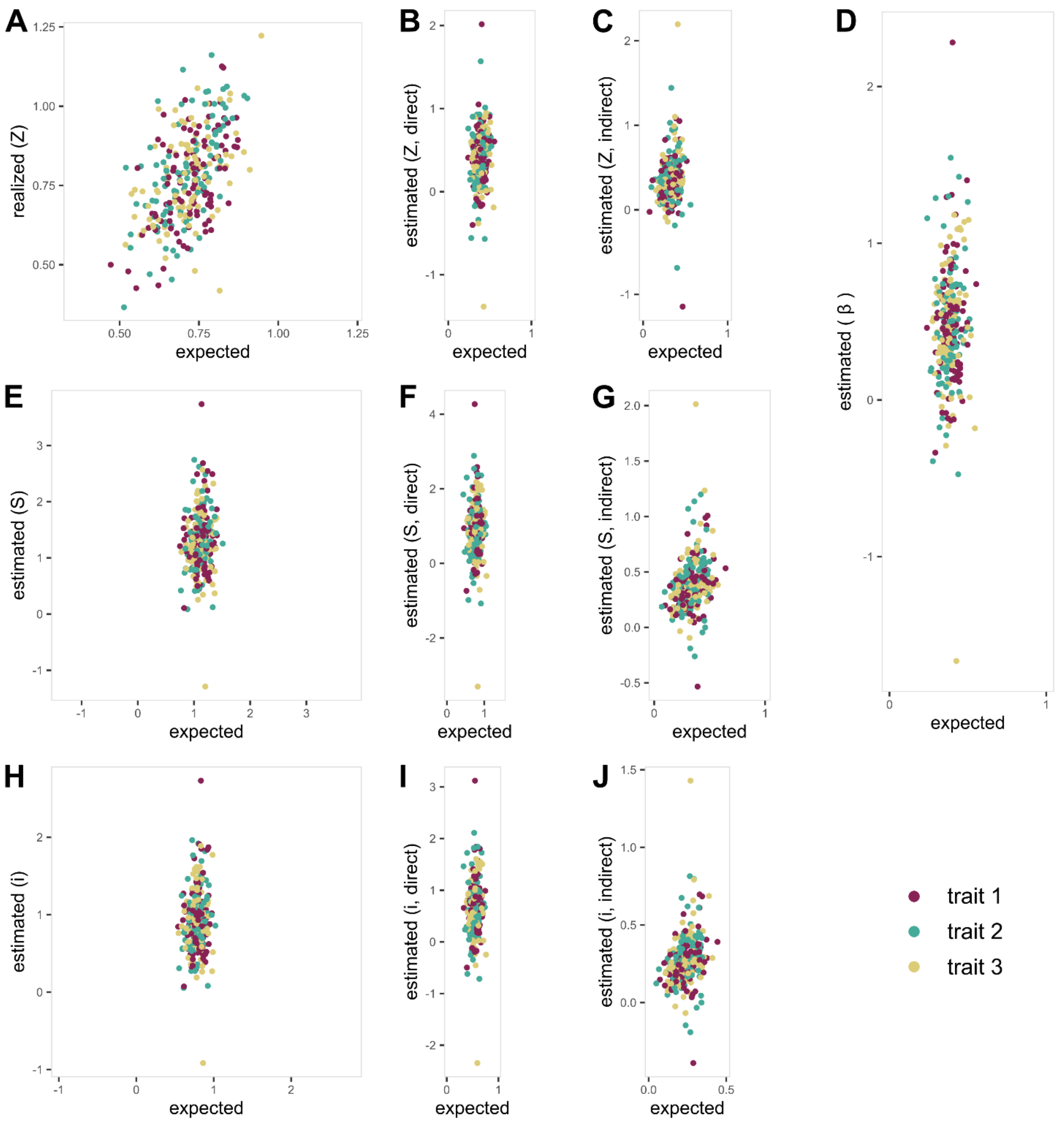


**Figure S15. Estimated multivariate selection parameters in Sel3.**

All estimated multivariate selection parameters are shown in the Y-axis and the true/expected values are shown in the X-axis. Note that selection responses are realized (*same as Figure S16A*) and do not depend on any estimated values from mixed linear model. [**A**] Selection responses, partitioned into [**B**] direct and [**C**] indirect. [**D**] Selection gradient. [**E**] Selection differential, partitioned into [**F**] direct and [**G**] indirect. [**H**] Selection intensity, partitioned into [**I**] direct and [**J**] indirect.

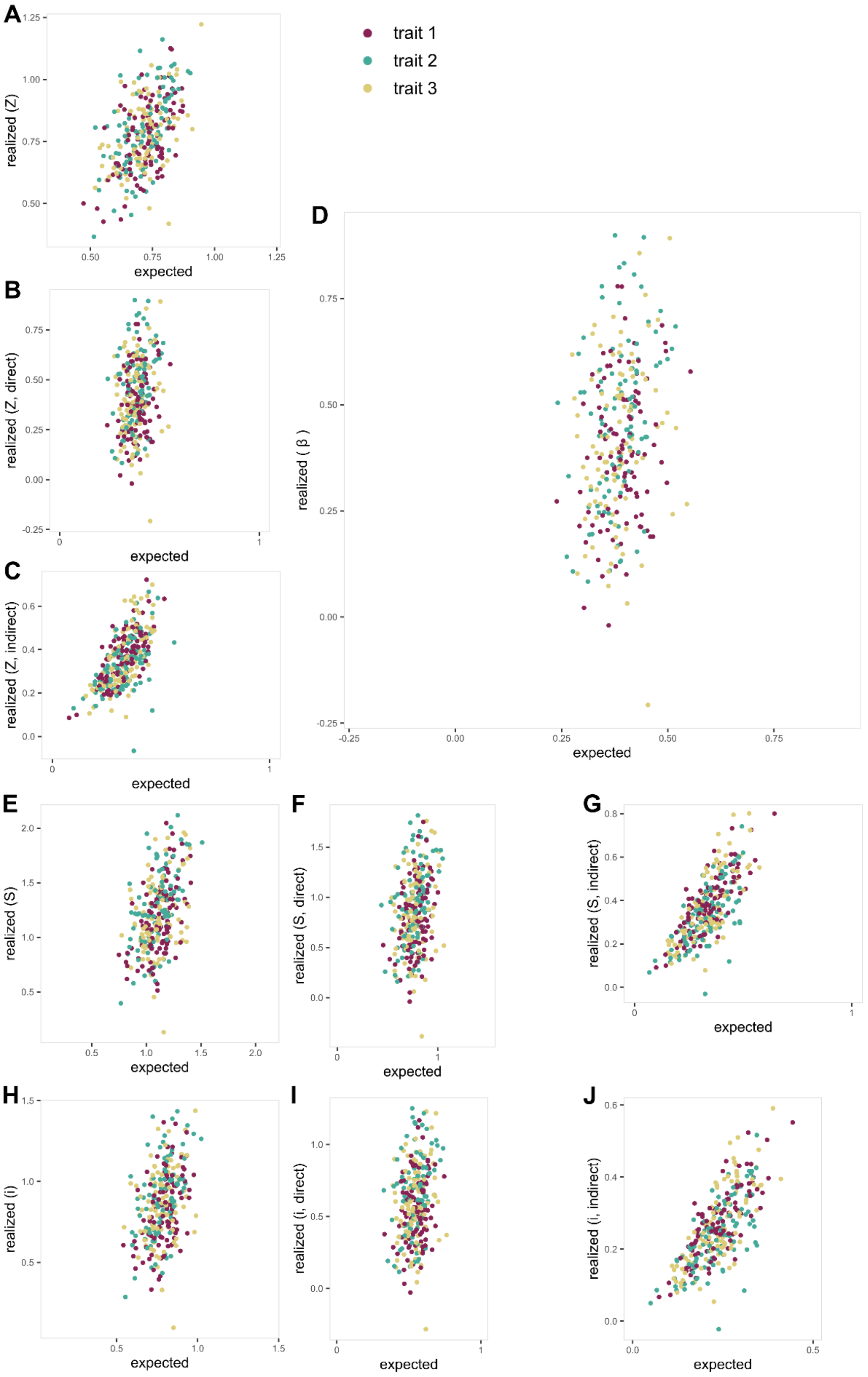


**Figure S16. Realized multivariate selection parameters in Sel3.**

All realized multivariate selection parameters are shown in the Y-axis and the true/expected values are shown in the X-axis. [**A**] Selection responses, partitioned into [**B**] direct and [**C**] indirect. [**D**] Selection gradient. [**E**] Selection differential, partitioned into [**F**] direct and [**G**] indirect. [**H**] Selection intensity, partitioned into [**I**] direct and [**J**] indirect.
